# Cryptic mitochondrial ageing coincides with mid-late life and is pathophysiologically informative in single cells across tissues and species

**DOI:** 10.1101/2023.07.04.547509

**Authors:** Alistair P. Green, Florian Klimm, Aidan S. Marshall, Rein Leetmaa, Juvid Aryaman, Aurora Gómez-Durán, Patrick F. Chinnery, Nick S. Jones

**Affiliations:** Department of Mathematics & Centre for the Mathematics of Precision Healthcare, Imperial College London, South Kensington, London SW7 2AZ, United Kingdom; Department of Clinical Neuroscience & Medical Research Council Mitochondrial Biology Unit, School of Clinical Medicine, University of Cambridge, Cambridge CB2 0QQ, UK; MitoPhenomics Lab, Centro Singular de Investigación en Medicina Molecular y Enfermedades Crónicas (CiMUS), Universidade de Santiago de Compostela, Campus Vida Avenida Barcelona, s/n, 15782, A Coruña (Spain); I-X Centre for AI in Science, Imperial White City Campus, 84 Wood Lane, London W12 7SL, UK

**Author notes:** Both authors contributed equally.

## Abstract

Ageing is associated with a range of chronic diseases and has diverse hallmarks. Mitochondrial dysfunction is implicated in ageing, and mouse-models with artificially enhanced mitochondrial DNA (mtDNA) mutation rates show accelerated ageing. A scarcely studied aspect of ageing, because it is invisible in aggregate analyses, is the accumulation of somatic mtDNA mutations which are unique to single cells (cryptic mutations). We find evidence of cryptic mtDNA mutations from diverse single-cell datasets, from three species, and discover: cryptic mutations constitute the vast majority of mtDNA mutations in aged post-mitotic tissues, that they can avoid selection, that their accumulation is consonant with theory, hits high levels coinciding with species specific mid-late life, and that their presence covaries with a majority of the hallmarks of ageing including protein misfolding and ER stress. We identify mechanistic links to ER stress experimentally and further give evidence that aged brain cells with high levels of cryptic mutations show markers of neurodegeneration and that calorie restriction slows the accumulation of cryptic mutations.

## Introduction

Diseases of later life represent a formidable human challenge (*1*). Existing hallmarks of ageing point to disparate but interconnected classes of causal factors leaving the search for unifying factors in humans open (*2, 3*). Somatic theories of ageing implicate DNA mutation as a possible cause — but nuclear mutation levels are possibly too low to be fully explanatory in post-mitotic tissue (*2*–*4*). Mitochondria and mtDNA mutation have been implicated in ageing (*5, 6*). MtDNA mutations certainly increase in number with age in tissue and cellular aggregates (*7*–*9*) with mutations predominantly created during mtDNA replication (*10*). There is emerging evidence that pathological mtDNA mutations are harboured at high levels in immune cells (*11*), and that these mutations can be linked to a cell phenotype (*12*). MtDNA mutator mice and cardiac and skin twinkle mutants (*13* –*16*) suggest that very high levels of mtDNA mutation can yield pro-geroid phenotypes (and this can be reversed (*16*)) but shorter-lived organisms and haploid mutator organisms yield a more nuanced picture (*17* –*20*). There is a separate debate about the relative role of mutations that are inherited or developmental and *de novo* mutations (*21*) with evidence pointing to selective effects in inherited and developmental mutations (*22*). Recently there has been an explosion of ageing-related single-cell transcriptomics data, though how the age of single cells is defined is unclear (*23, 24*) with mixed evidence suggesting a link between ageing and gene-expression variance (*24*). While single-cell transcriptomics has been shown to allow variant calling in nuclear and mtDNA (*25, 26*) there have been no efforts to link single cell data to mtDNA ageing.

Drawing on single cell data from over 124,000 cells, three mammalian species and seven tissues, using a mix of single-cell RNA and ATAC sequencing (scRNA-seq, scATAC-seq), we link ageing in post-mitotic tissues to an understudied type of mtDNA mutation (cryptic: those mtDNA mutations which are unique to single cells in a sample) which is invisible in aggregate. We give evidence that these mutations constitute the dominant fraction of tissue mtDNA mutations, accumulate in a manner which coincides with species lifespan and are consonant with new theory we develop. This theory predicts that mutations will reach functionally relevant heteroplasmies within human lifespan, and infers an mtDNA mutation rate consistent with existing literature values. Through new experiments and informatics we find evidence that cryptic mtDNA mutations are linked to the expression of genes linked to disrupted proteostasis and immune/inflammatory response. Looking across rat, human, and mouse we find links between cryptic mutation and multiple hallmarks of ageing. We further find evidence that in aged neurons the presence of cryptic mutations correlates with markers of neurodegeneration and that caloric restriction slows the accumulation of cryptic mutations.

### Cryptic mutation is predominant and its accumulation coincides with lifespan

It has been shown that scRNA-seq can be used for mtDNA mutation identification (*25, 27, 28*), and we further corroborate the validity of this technique (see Supplementary Discussion S6). We leverage mutational information gained from scRNA-seq and scATAC-seq to study the presence of mtDNA mutations at different ages.

The mtDNA heteroplasmy *h* of a mutation is the proportion of mtDNA molecules in a cell that bear that mutation. Empirically, we assign a heteroplasmy value to each mutation based on the fraction of reads carrying that mutation (see Eq. 1 in Methods). In the following we exploit distributional and comparative approaches to ensure robustness to inevitable errors in sequencing and variant inference: in particular, we consider distributions of cryptic heteroplasmies where each mutant site is found in only one cell amongst all cells from a given donor and has an associated heteroplasmy. A distribution of cryptic heteroplasmies can be found from a collection of cells by recording the heteroplasmies of all cryptic mtDNA mutations found in each cell in the sample and building a histogram showing the frequencies with which different heteroplasmies are observed. We term this distribution of cryptic heteroplasmies the ‘cryptic’ site frequency spectrum (cSFS) and it is a natural object from population genetics. By taking the distribution of these heteroplasmies, we can find the probability that a cryptic mutation, picked at random from the set of all cryptic mtDNA mutations in these cells, has a particular heteroplasmy (we use an extended notion of the site frequency spectrum and include homoplasmic (100 % heteroplasmic) mutants (examples in Figs. 1c, f)).

**Figure 1.**
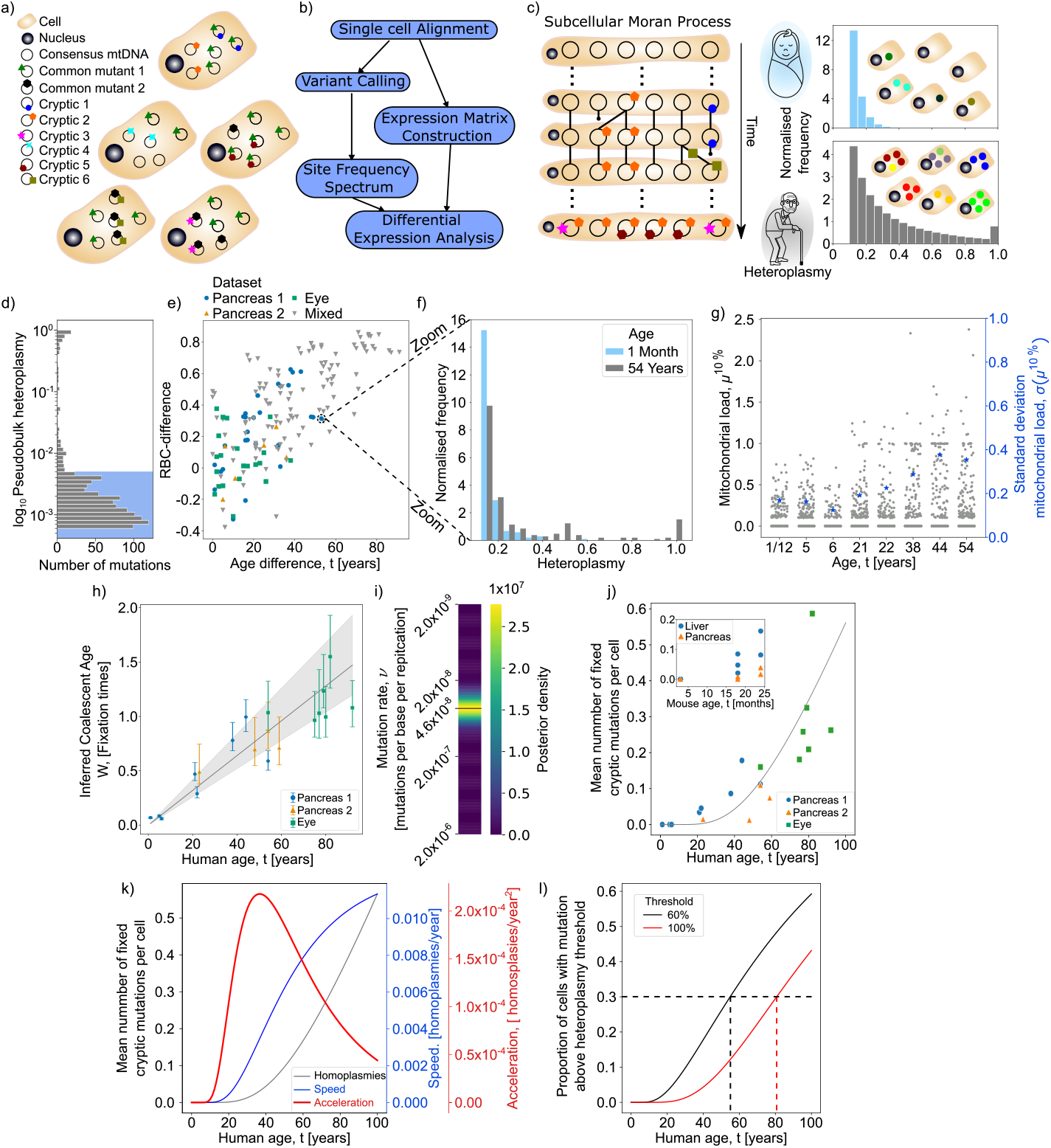
Cryptic mtDNA mutation is a predominant form of mutation, reaches physiologically relevant levels in later life and evolves in a clock-like manner consonant with theory. (a) Cells in tissues carry two mutant types — those common to multiple cells in the tissue, and those unique to a single cell. (b) We exploit scRNA-seq and scATAC-seq on tissues taken from individuals of varying ages and pool variants from every cell allowing us to not only construct an expression matrix, but to examine the site frequency spectrum of a tissue and link changes in expression of single cells to inferred mtDNA mutant load. (c) Modelling sub-cellular population genetics with an out-of-equilibrium infinite-sites Moran model we can predict how the normalised site frequency spectrum of cryptic mtDNA mutations evolves over a lifetime and predict an accumulation of high heteroplasmy mutations in aged individuals. (d) Single-cell sequencing is necessary to reveal true mutant load of a tissue. 90.8 % of mtDNA mutations are at a pseudobulk heteroplasmy *h <* 0.5 % (marked in blue) and so would not be reliably detected in most bulk experiments (Data taken from all mutations found in (*29*)) (e-f) The cryptic site frequency spectrum (cSFS) evolves with time. We see that for 19 individuals across 3 human datasets taken from different tissues and differing sequencing techniques, the rank biserial correlation distance (RBC-difference, a measure of how likely a mutation sampled from one spectrum will have a higher heteroplasmy than a mutation sampled from another) between two spectra increases with the age difference between the individuals (Spearman correlation *r ≈* 0.70 and *p <* 10^*−*26^). By looking directly at an example pair of spectra (f) we see that, as predicted by our theory, there is an accumulation of mutants at high heteroplasmies (for a breakdown of the number of cells and mutations represented in the cSFS see table S2). (g) The mitochondrial load *μ*^10 %^ of potentially pathogenic, cryptic mutations increases with age in human pancreas cells. We show the mitochondrial load *μ*^10 %^ for all eight donors in Ref. (*29*) and observe an increase with age (Spearman correlation *r ≈* 0.04 and *p <* 0.049) and observe an even stronger effect for larger heteroplasmy thresholds (see Supplementary Discussion S8.5) The standard deviation *σ*(*μ*^10 %^) of the mitochondrial load also increases with age (blue asterisks; Spearman correlation *r ≈* 0.88 and *p <* 0.005). (h) We present the 95 % credible interval for each donor’s mitochondrial age as a proportion of the expected time to fixation, and show the maximum a posteriori (MAP) estimate for regression, along with the 95 % credible interval of the median. (i) The posterior on the inferred mutation rate per base per replication is shown under the assumption of 1000 mtDNA per cell, with the MAP estimate highlighted as 4.6 *×* 10^*−*8^. (j) The number of homoplasmic mutations per cell increases with age in humans (Spearman correlation *r ≈* 0.89 and *p <* 10^*−*7^). Also shown is a line generated with the MAP parameters found from fitting the data to our model (Supplementary discussion S2). We see a similar accumulation of homoplasmic mutations in mice in samples from 2 tissues and 16 mice (Spearman correlation *r ≈* 0.85 and *p <* 10^*−*4^). (k) We look at the first and second derivatives of the MAP estimate for number of homoplasmic mutations against time. These can be roughly equated to the speed and acceleration of ageing. The peak in acceleration of ageing occurs at around 40 years old. (l) Using the MAP estimate we look at the proportion of cells carrying a mutation above a certain threshold. After around 20 years cells begin to have mutations above a 60 % heteroplasmy threshold and by age 40 an appreciable fraction of cells are carrying a mutation with heteroplasmy above 60 % and cells have begun to accumulate mutations at homoplasmy (100 % heteroplasmy). By the age of 80 nearly 30 % of cells are predicted to have a homoplasmic mutations and nearly half are predicted to have mutations at a heteroplasmy above 60 %.

We first examine five single-cell RNA-seq datasets covering two species, three tissue types, and three sequencing techniques to demonstrate the wide relevance of our results (*29* –*32*). With single-cell-level data we see how mutations which would be detectable at *h ≥* 0.5 % heteroplasmy in a bulk sequencing experiment make up only *∼* 9 % of the mutations in a tissue (Fig. 1d). Of the *∼* 91 % of mutations that are at *h <* 0.5 % in bulk heteroplasmy almost all (*∼* 94 %) are only found in single cells.

We hypothesise, supported by theory (discussed later), that the cSFS will evolve with time gradually spreading to higher heteroplasmies, Fig. 1c. We compare the cSFS from human donors of different ages to see how it evolves through life and find that, consistent with the hypothesis, the further apart in age two individuals are, the larger the difference in their cSFS, as measured by the rank biserial correlation difference, RBC-difference, between pairs (see Figs. 1e-f each data point is a comparison between all the cells of two individuals, see Supplementary Fig. S22 for alternative metric comparisons; these results are preserved when restricting to a single cell type (see Supplementary Fig. S23a). The RBC-difference of two cSFSs (A and B) is a measure of how likely a mutation sampled from A has a higher heteroplasmy than a mutation sampled from B (see Methods). To exclude possible errors from library preparation and sequencing, we only consider cryptic mutations with heteroplasmy *h >* 10 % (see Methods and Supplementary Discussion S6).

Next, we identify cells in a single-cell human pancreas data set (*29*) which carry mutations that are likely dysfunctional. For this, we compute each cell’s mtDNA load of cryptic mutations *μ*^10 %^ as the sum of the heteroplasmies above 10 % of all cryptic mutations which are not synonymous protein coding mutants (see Eq. 2 in Methods). We find that the mtDNA mutant load *μ*^10 %^ increases with age (see Fig. 1g). Furthermore, we observe that the standard deviation of the mtDNA mutant load increases with age, indicating a increasing age-associated cellular heterogeneity. The correlation *Corr*(*μ*^*h* %^, *t*) between age *t* and mtDNA mutant load *μ*^*h* %^ is significant for all heteroplasmy thresholds *h >* 10 % (see Supplementary Figs. S25, S26), indicating that the accumulation of cryptic mutations occurs already at a low-heteroplasmy threshold but is observable across a wide range of heteroplasmies.

To confirm that these results reflect the underlying mtDNA, we also examined a 10x single-nucleus ATAC-seq dataset from the aged human brain, offering an orthogonal sequencing modality, which supports our hypothesis that older individuals have a cryptic site frequency spectrum which is shifted to higher heteroplasmies (see Supplementary Fig. S17).

### Cryptic mutations reach physiologically relevant levels in a manner consonant with theory

Using an established model from population genetics, the Moran process (*33*), we derived a formula for how the cSFS should evolve with time if a cell starts with no cryptic mutations and gradually acquires them over a lifetime, Fig. 1c (see Methods and Supplementary Discussions S1 and S2 for details). The formula predicts that the cSFS histogram should have an amount of mutation at higher heteroplasmies that increases with age (Fig. 1c) and which has a characteristic age at which high-heteroplasmies are reached. In the model mtDNA replicate and are eliminated in a manner that keeps the population constant; *de novo* mutations occur with a fixed rate and mutations spread through the population because of random birth and death. For the Moran process, in the long-time limit, the expected time for the set of mtDNA in a cell to have a single common ancestor is proportional to the product of the number of mtDNA in the cell and the half-life of the mtDNA population: rapid birth-death or small populations lead to faster fixation of mutations.

We used Bayesian inference to fit our model for the cSFS to the human datasets. In brief, our model fits a ‘mitochondrial age’, *W*, and scaled mutation rate, Θ, to a set of cells from a donor tissue. We find that the inferred mitochondrial age increases with the chronological age of individuals: providing a novel biological age marker for tissues (Fig. 1h). By making an assumption about the mtDNA copy number of cells, *N*, we can convert the inferred scaled mutation rate Θ to a mutation rate per base per replication, *ν* using the equation Θ = *Nν*. Under the assumption of 1000 mtDNA per cell, we infer a maximum a posteriori (MAP) estimate of the median mutation rate per base per replication of 4.6 *×* 10^*−*8^ (Fig. 1i, see Supplementary Discussion S2 for details and full fitting results including showing that the trends are recapitulated using only alpha cells from Ref. (*29*)).

A core model prediction is that the heteroplasmies of the cSFS increase until they reach a steady state, while the average number of homoplasmies is constantly increasing. At steady-state, heteroplasmic mutations are continually being lost or fixing at homoplasmy, while homoplasmic mutations accumulate, since they have no avenue to be lost from the population (*33*). Experimental data in both human and mouse show a non-linear increase in homoplasmies with age, even across datasets and tissue types (Fig. 1j, this trend is also seen when restricted to a single cell type, see Supplementary Fig. S23b). Most strikingly, the data shows that after 80 years of age in humans, over 20% of cells carry a mutation with heteroplasmy *h >* 95 %. We provide a line indicative of our theory, produced using the MAP estimate for ageing rate and mutation rate found using our model (see Supplementary Discussion S2 for details and full fitting results). Notably shorter lived animals (mouse inset (Fig. 1j)) also reach high levels of homoplasmy much faster: reaching high numbers of homoplasmic cells coincides with these organism’s lifespan. We also examine the first and second derivatives of the fit for number of homoplasmic mutations (Fig. 1k). These can be thought of as both the speed and acceleration mtDNA ageing respectively. The shape of the speed of cryptic mtDNA ageing is roughly sigmoidal, with the ageing speed only beginning to increase after around 20 years. The acceleration of ageing peaks at around 40 years old when many of the more obvious signs of ageing have begun to present themselves. As noted the heteroplasmies in the cell (unlike the homoplasmies) do reach equilibrium and this timescale (using the same parameters as in Fig. 1k) coincides with lifespan Fig. 1j. We can also look at the fraction of cells carrying a mutation above a certain heteroplasmy threshold. Again using the MAP estimate of model parameters we see that by 50 years old, over 30 % of cells are expected to carry at least one mutation at a heteroplasmy *>* 60 %. The time at which 30 % of cells are expected to carry at least one mutation at homoplasmy is 70 years old (Fig. 1m). These two thresholds demonstrate the multiple timescales at which mtDNA mutations can accumulate in tissues. We note that a third timescale is implied by this evidence: by age 20 mutations at frequencies ¡10 % have mostly reached their life-long levels (Supplementary Fig. S24).

### Cryptic mutations, unlike other types, can expand neutrally

Access to the cSFS of single cells allows us to examine selection against pathogenic mutations in more depth than the non-synonymous/synonymous mutant ratio. We can assign each protein coding mutation a pathology class of either synonymous, low pathogenicity, or high pathogenicity (see Methods) and then examine the cSFS of the three pathology classes. By comparing this with a more conventional measure of selection, the non-synonymous/synonymous ratio, we can look in more detail at whether selection effects are dependent on the heteroplasmy of mutations. We perform this analysis on cryptic mutations taken from a high-quality full-length scRNA-seq data of healthy pancreas cells (*29*) and find that, for mutations above 10 % heteroplasmy, there is no evidence of a significant shift in the cSFS of low or high pathogenicity mutations when compared to synonymous mutants (Fig. 2a), and the non-synonymous/synonymous ratio is not significantly shifted from 1: mutations that reach a heteroplasmy of *>* 10 % do not show evidence of selection. We repeat this analysis considering all non-cryptic mutations and find that as well as having a non-synonymous/synonymous ratio significantly shifted below 1, the SFS of the average heteroplasmy of non-cryptic, highly pathogenic mutations are also significantly shifted to lower heteroplasmies, which is consistent with evidence of selection happening along the human germline and in development in a way which is modulated by the degree of pathogenicity the mutation causes (*34*) (Fig. 2b). For comparative purposes we perform this analysis on scATAC-seq data for cultured cell lines (*35*) and find that, analogously, the SFS of all mutations with both high and low pathogenicity are significantly shifted to lower heteroplasmies compared to synonymous ones, indicating selection against high heteroplasmy pathogenic mutants (Fig. 2c).

**Figure 2.**
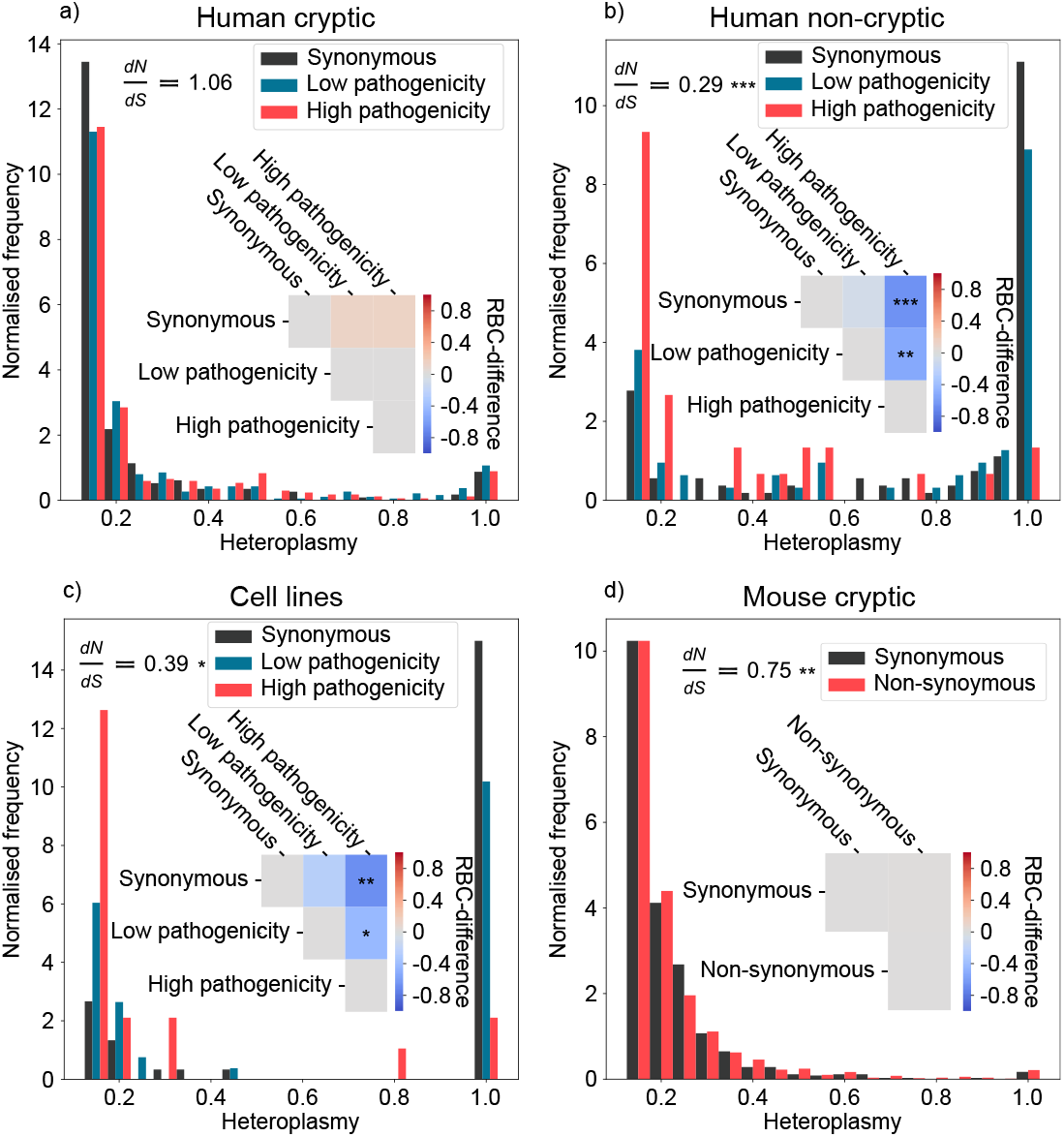
Cryptic mtDNA mutations, unlike other types, can expand neutrally. We show the SFS for mutations classified by their potential pathology, as well as the RBC-difference (see Methods) which shows the magnitude of difference between these SFSs (for a breakdown of the number of cells and mutations represented in each SFS see table S2). (a) All cryptic mutations taken from healthy human pancreas tissue (*29*) show no significant evidence of selection in either the non-synonymous/synonymous ratio or in the RBC-difference between cSFSs, indicating these mutations are not under selective pressure. (b) When we consider all non-cryptic mutations, we see a non-synonymous/synonymous ratio significantly less than 1 (Fisher’s exact test *p <* 10^*−*8^) as well as a significant shift of the SFS of high pathogenicity mutations as compared to synonymous and low pathogenicity mutations (Mann-Whitney U test *p <* 10^*−*6^, 10^*−*3^ respectively), though we do not see evidence for selection against low pathogenicity mutations compared to synonymous, suggesting that mutations occurring on the germline or during development experience selective pressure based on the level of dysfunction they cause. (c) The SFS for all mutations found in cells from culture (*35*) show a similar selection pattern to non-cryptic mutations taken from tissue samples. They have a non-synonymous/synonymous ratio significantly less than 1 (*p <* 10^*−*3^) and the spectra of high pathogenicity mutations show a significant shift towards lower heteroplasmies when compared to synonymous and low pathogenicity mutations (*p <* 10^*−*4^, 0.05 respectively). (d) Though we cannot score the potential pathogenicity of non-synonymous mutations in mice, we look at the difference in the cSFS of synonymous and non-synonymous mutations finding no evidence of selection. We do, however, find a significant lack of non-synonymous mutations at heteroplasmies *h >* 10 % (*p <* 10^*−*4^).

Though we cannot score the potential pathogenicity of mtDNA mutations in mice, we can compare the cSFS of synonymous and non-synonymous mutations above 10 % from both liver and pancreas (Fig. 2d): we find no evidence of a shift in the cSFS between these two classes (as we did in Fig. 2a). We do, however, discover a significant shift in the non-synonymous/synonymous ratio below 1, and find evidence for analogous behaviour in human brain tissue (see Supplementary Fig. S17). This is consistent with a model where, for some tissues, some non-synonymous mutations undergo strong negative selection at heteroplasmies *<* 10 % (perhaps selective mitophagy) but where mutations that evade this selective mechanism expand neutrally in a manner independent of their heteroplasmy.

These results, taken with the life-time evolution of the cSFS (Fig. 1d,f,h), point to the potential for accumulation of cryptic pathogenic mtDNA mutations through life, causing mosaic dysfunction in post-mitotic tissues.

### Cryptic mutation links to cellular phenotype in a manner consonant with markers of ageing pathophysiology

The number of cells with evidence of high-heteroplasmy cryptic mutations increases nonlinearly with age. To identify which genes’ expression levels might be perturbed by the presence of cryptic mutations, we perform a differentially expressed genes (DEG) analysis: comparing cells with detected cryptic mutations which are not synonymous above 10% heteroplasmy and those without (see Methods) for the full-length scRNA-seq pancreas data (*29*). After multiple-testing correction, we find 1342 genes significantly differentially expressed (see Fig. 3a), consonant with a possible large-scale transcriptional perturbation induced by cryptic mutations. First, as expected, we find mitochondrially-encoded (e.g., MT-CO2, MT-RNR1) and nuclearly-encoded (e.g, NDUFA1, NDUFA13) OXPHOS genes upregulated, which is an established response to an impaired energy production (*36*). Second, we identify genes associated with innate immune signalling and altered proteostasis (HSPA5, HSPA13, HSP90B1, and YME1L1 (*37*)) downregulated and key inflammatory cytokine MIF upregulated (*38*).

**Figure 3.**
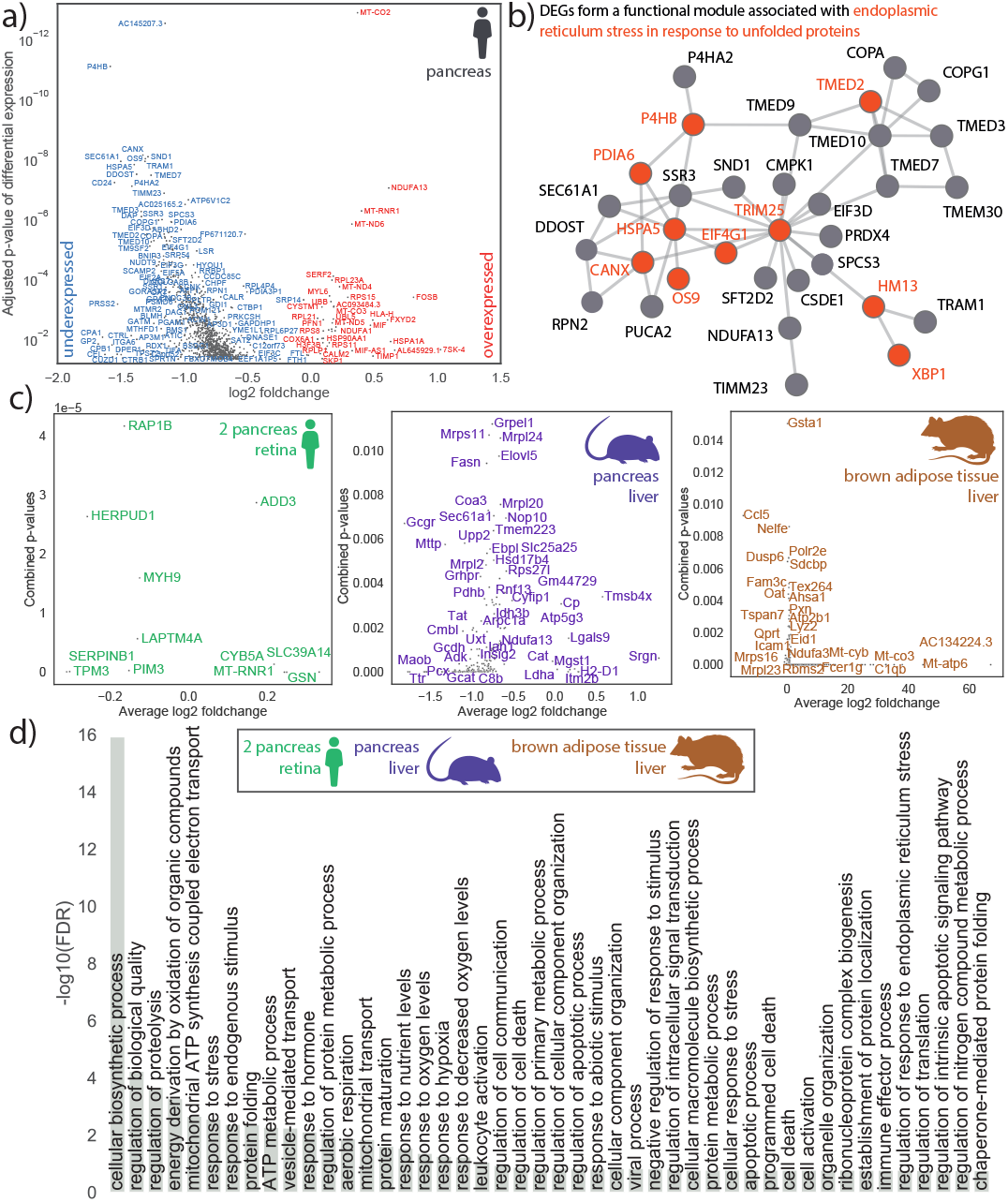
Single-cell transcriptional hallmarks of ageing covary with cryptic mutations. (a) Volcano plot of DEGs in cells with cryptic mutations which are not synonymous above 10% heteroplasmy, and those without, in the human pancreas data (*29*). (b) Genes whose expression covaries with the presence of mutations form a functional protein-interaction module which is enriched with markers of stress response. To obtain the functional module, we combine the *p*-values of differential expression with the curated protein interaction network from BioGRID by computing a maximum-weight connected subgraph with scPPIN (*41*). (c) DEGs in cells with cryptic mutations which are not synonymous for human, mice, and rat. (d) Selected GO terms that are enriched in the DEGs of at least six of the seven data sets (2 human pancreas, human retina, mouse pancreas, mouse liver, rat liver, rat brown adipose tissue, see Data section for details). We order the GO terms by the false discovery rate (FDR) in the data from Ref. (*29*).

Surprisingly, we identify an altered expression of long noncoding RNAs (lncRNAs), such as AC145207.3 and MIF-AS1, which have been hypothesised to play a role in ageing (*39*) and cancer cell proliferation (*40*). While we here aggregate data from all donors to extract even subtle changes in gene expression, we also observe a perturbation of gene expression at the level of single donors or cell types (see Supplementary Discussions S3 and S4).

Since proteins fulfil their biological functions through interaction, we then use scPPIN (*41*) to integrate the *p*-values of differential expression with protein–protein interaction data and obtain a mutation-linked functional module consisting of 33 proteins (see Fig. 3b). This module is associated with endoplasmic reticulum (ER) stress in response to unfolded proteins and is consistent with mtDNA mutations yielding misfolded proteins, which trigger an ER stress response (as is known for ageing-associated diseases of various tissues (*42, 43*), including Alzheimer’s disease). We find multiple Transmembrane emp24 domain-containing proteins to be perturbed, which hints at a dysregulated immune response (*44*). TRIM25, a ubiquitin ligase that regulates the innate immune response, is at the centre of the module, highlighting the interplay between mtDNA mutation, immune response, and ER stress (*45* –*47*).

We perform an analogous DEG analysis for seven scRNA-seq data sets from three different mammals (human, mice, and rat) and four different tissues, and identify in each of them that the cryptic mutations are linked to gene expression changes (see Fig. 3c and Methods). We identify biological pathways linked to the presence of cryptic mutations across organisms by performing a GO-term enrichment analysis with PANTHER (*48*), highlighting terms that are enriched across at least six of seven data sets (see Fig. 3d). Cryptic mutations coincide with a perturbation to the regulation of biological quality, response to stress, ER-stress, viral response, leukocyte activation, apoptosis, hypoxia and proteolysis, and protein-folding. Combined with an enrichment of immune effector process these terms are consistent with an immune response triggered by cryptic mutations: in line with recent findings linking neo-epitopes to *de novo* mtDNA-mutations (*45, 47, 49*). An enrichment of response to nutrient levels indicates that these processes might interplay with dietary interventions, a hypothesis that we test in the following section. Beyond genomic instability (a hallmark of ageing) the transcriptional discrepancies between cells with cryptic mtDNA load and those without are consonant with four further hallmarks of ageing (loss of proteostasis, deregulated nutrient-sensing, mitochondrial dysfunction, and altered intercellular communication).

Since the cryptic mutations we observe are diverse (and unique to each cell in the sample) we expect them to each create distinctive modulations to gene expression: nonetheless, Fig 3d suggests common patterns, which can be accounted for as follows. In mitochondrial physiology it is uncontroversial that mitochondrial-disease related mtDNA mutations (e.g., LHON, MELAS) can cause changes in gene expression (we find LHON-associated mutations in 39 of the cells in the long-read human pancreas data (*29*)). The changes in Fig. 3d are consistent with the perturbations already known to be created by mitochondrial-disease mutations: changes related to hypoxia, ETC, ER-stress, and protein folding (*50*). It is known that Complex I mutations have a marked effect in mitochondrial disease (*51*) – we find that mutations in the mt-ND4 and mt-ND5 genes are strongly associated with changes in cellular gene expression. The former being linked to ‘response to unfolded protein’ and the latter linked to ‘ATP synthesis coupled electron transport’ (see Supplementary Fig. S36). This points to a picture of ageing as partly a mosaic of different single-cell mitochondrial diseases.

Following our observations regarding complex I mutations we developed select cell lines containing mtDNA mutations (Fig. 4a, with background matched controls) that we had separately identified as cryptic complex I mutations in our analysis of single cell data: these lines thus allow us to functionally explore the effect of exemplar cryptic mutations (while carefully controlling for other mtDNA mutations). We observe mutation-specific ETC (Fig. 4b) and mitochondrial ATP deficiency (Fig. 4c) with elevated ROS levels (Fig. 4d) and bioenergetic differences (Fig. 4e). Transcriptomic analysis shows a pronounced effect on gene-expression (Fig. 4f-g). Pathway enrichment analysis yields evidence for ER-stress and altered proteostasis (Fig. 4g, e.g. AFT6, XBP1(S) and IRE1alpha) consonant with our previous observation (Fig. 3d) and we corroborated this by finding experimental evidence for the activation of Eukaryotic translation initation factor 2 alpha a key kinase that responds to ER stress (*52*)(Fig. 4h).

**Figure 4.**
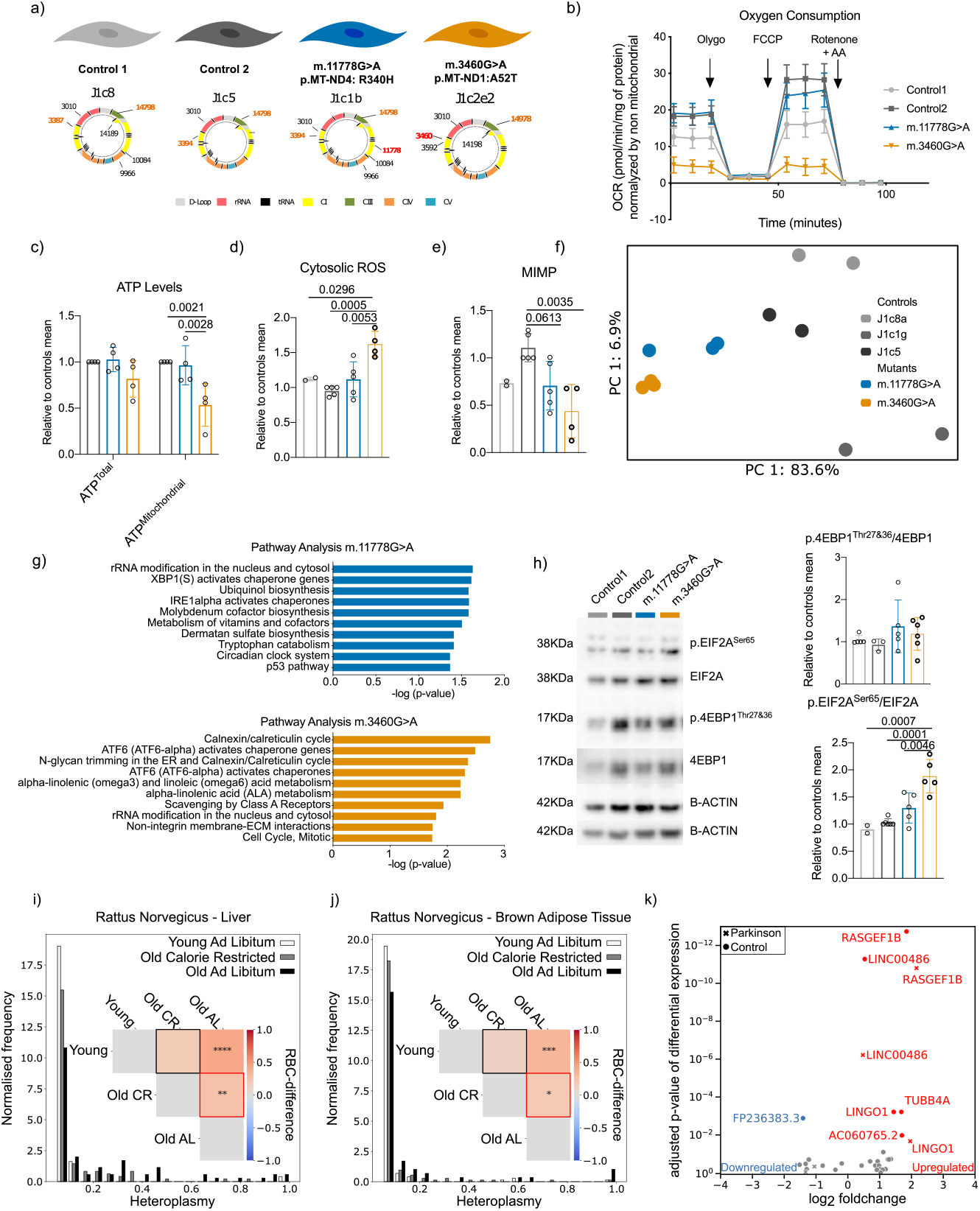
Evidence that cryptic mtDNA mutations can have functional effects including ER-stress, have heteroplasmy levels that can be controlled for therapeutic benefit and are connected to neurodegeneration-linked genes. (a) Graphical representation of the mtDNAs of the model. The mitochondrial population background and location of the mtDNA non-synonymous variants are shown. Full mtDNA sequences is included in supplementary table S6. (b) Oxygen consumption rate (OCR) from a representative experiment. The Complex V inhibitor oligomycin (Oligo), the uncoupler carbonyl cyanide 4-(trifluoromethoxy)phenylhydrazone (FCCP), and the Complex I and III inhibitors antimycin (AA) and rotenone (Rotenone) were added sequentially at the indicated time points. (c) ATP levels quantification for total and mitochondrial (glycolysis inhibited by incubation with d-dexy-glucose instead of glucose). (d) Representative example of FACS determination for cytosolic ROS and quantification of the levels. (e) Representative example of FACS determination for MIMP and quantification of the levels. (f) PCA representation of all the DEG genes in both comparisons (DEGs resulting from pair to pair comparisons are included in Extended Data). (g) Mutation specific pathway analysis (full report is included in Extended Data). (h) Analysis of the EIF2A and 4EBP1 activation. Example of immunoblots. The blots were immuno-detected using an anti-EIF2A, anti-p.EIF2ASer51, anti-4EBP1, anti-p.4EBP1Thr37&46 and *β*-actin as loading control. Quantification of immune-detected bands for p.EIF2ASer51 and EIF2A as well as anti-p.4EBP1Thr37&46 and anti-4EBP1corrected with values from the loading control (*β*-actin) in each cell line. Activation of the kinases is calculated as the ratio of phosphorylated/ non-phosphorylated. Bars and lines represent mean *±* s.d. of the biological replicates (n = 4) in all the cell lines. Values are represented relative to the average of control cell lines unless indicated. Statistical testing was performed by using a 1-way-ANOVA test followed by Holm-Sidak’s multiple comparisons test unless stated otherwise. Exact p-values corrected for multiple comparisons are indicated. (i-j) We find the cSFS for rats in three groups, young ad libitum (Y-AL), old ad libitum (O-AL) and old calorically restricted (O-CR), for variants called at a 5% heteroplasmy. This is displayed for liver in (i) and in (j) for brown adipose tissue. Pairwise RBC-difference (see Methods) between each group’s cSFS indicates that caloric restriction slows the accumulation of cryptic mutation. This comparison was done between all pairings of the three groups and is displayed in the inset in (i) for liver and in (j) for brown adipose tissue. Mutations in the O-CR are marginally more likely to be at a higher heteroplasmy than the Y-AL in both the liver and the brown adipose tissue (*p >* 0.05). Compared to this, mutations in the O-AL are far more likely to be at a higher heteroplasmy than Y-AL mice (Mann-Whitney U Test *p <* 0.0001 & *p <* 0.001 Bonferroni corrected in liver and brown adipose tissue respectively). Critically, mutations in O-AL rats are statistically significantly more likely to be at a higher heteroplasmy than mutations found in O-CR (Mann-Whitney U test *p <* 0.01 & *p <* 0.05 Bonferroni corrected in liver and brown adipose tissue respectively). Mean heteroplasmies of observed liver mutations in Y-AL, O-CR, O-AL respectively are 0.0846 *±* 0.0025, 0.115*±*0.0013, 0.228*±*0.0035. In BAT the means are 0.0796*±*0.00068, 0.0909*±*0.00072, 0.144*±*0.0035. (k) High-heteroplasmy mutational load *μ*^95 %^ in single brain cells coincides with differential expression of neurodegeneration-linked genes.

### Implications for calorie restriction and disease

Given the possible causal link between mtDNA mutation and ageing we asked first whether levels of cryptic mtDNA mutations could be controlled through an established anti-ageing technique, caloric restriction, and second, whether we could identify links between cryptic mutation and neurodegeneration (via single-nucleus-RNA-seq).

#### Caloric restriction

Caloric restriction has been recognised as one of the most effective interventions to promote longevity, and combat ageing (*53*). We use scRNA-seq data from young and old *ad libitum* fed rats (Y-AL and O-AL, respectively) and old calorically restricted rats (O-CR) (*54*) to obtain cSFS of each group’s liver and brown adipose tissue. Using the RBC-difference (see Methods) we observed that (as anticipated in Fig. 1f) cryptic mutations are of higher heteroplasmy in the O-AL than Y-AL (Fig. 4i, j), in both liver (*p <* 0.0001, Bonferroni corrected) and brown adipose tissue (*p <* 0.001, Bonferroni corrected). Likewise, mutations in O-AL are higher in heteroplasmy than in O-CR, in both liver (*p <* 0.01, Bonferroni corrected) and brown adipose tissue (*p <* 0.05, Bonferroni corrected). By contrast, the mutations in O-CR rats are not significantly different in heteroplasmy to those found in Y-AL rats. These findings suggest that caloric restriction slows the rate of increase of average cryptic heteroplasmy, this mechanism might be an explanatory factor for the observed longevity in calorically restricted organisms. Caloric restriction can increase the number of mtDNA molecules in rat livers (*55*): our theory suggests that increasing mtDNA-copy-number slows the rate of increase of mean heteroplasmy (see Supplementary discussion S1). An analysis of genes that are differentially expressed in rat cells with cryptic mutations highlighted the differential expression of cystatin C and Apolipoprotein E and enriched GO terms include ‘apoptotic mitochondrial changes’ and ‘ageing’.

#### Disease

Mitochondrial dysfunction is implicated in neurodegenerative diseases such as Parkinson’s disease (PD) (*56*) and Alzheimer’s disease (AD) (*57*). We use snRNA-seq (*58*) to investigate whether there is evidence for a perturbation of gene expression in the presence of high-heteroplasmy mtDNA mutations. As snRNA-seq provides sparse coverage of mtDNA transcripts (see Fig. S9) we reduce the minimum coverage for calling heteroplasmies to 10 reads and, to remove potentially falsely called variants, only consider cryptic mutations with a heteroplasmy of at least 95 % to identify DEGs in PD and control group, separately (see Fig **??**k). We find three genes significantly upregulated in cells with cryptic mtDNA mutations in both groups: LINGO1 has been associated with various neurodegenerative diseases by inhibiting regeneration in the nervous system (*59*) and the Guanine nucleotide exchange factor RAP2A has been associated with a population of excitatory neurons in AD (*60*). The lncRNA LINC00486 is overexpressed in cells with cryptic mutations and has been associated with common bipolar disorder (*61*). For an analysis of AD data that also identifies lncRNAs, specifically MALAT1 see Supplementary Information S5. Further evidence that disease can modulate mitochondrial parameters emerged when we found different mitochondrial ageing rates for diabetic and healthy pancreas tissue (for full results see Supplemetary Information S2.6).

## Conclusion

We find evidence that an understudied type of single-cell mutation, cryptic mtDNA mutations, while invisible in aggregate, are predictive of age and markers of ageing and show pathologically-relevant levels of heteroplasmy at middle age and late life. We find evidence that, in post-mitotic tissues, cryptic mutations can evade negative selection, expanding neutrally, and are linked to 5 of 9 hallmarks of ageing (*2*) (genomic instability, loss of proteostasis, deregulated nutrient-sensing, mitochondrial dysfunction, and altered intercellular communication); specific proteins point to pathways involving mitoprotein-folding, the ER stress, and immune responses (*62*), and we corroborated this with experiment. The gene-expression changes we observe are consonant with a mosaic combination of the established changes caused by well-studied mitochondrial-disease-associated mtDNA mutations (*50*). We conclude with evidence that an anti-ageing therapy can reduce the rate of accumulation of cryptic mtDNA mutations.

## Methods

We construct an analysis pipeline that allows us to identify mtDNA mutations in single cells from single-cell sequencing data. The pipeline enables the analysis of single-cell RNA-seq data (e.g., Smart-seq2, CEL-Seq2, 10x Genomics’ Chromium™ Single Cell 3’ Solution, and 10x Genomics’ Chromium™ Single Cell 3’ Solution (single-nuclei protocol), and ATAC-seq) but can be adapted to analyse other sequencing data and is a collection of custom-made Shell and Python code.

### Data access

#### Sequencing data

We download publicly available sequencing data from the Gene Expression Omnibus with the SRA Toolkit, specifically the fasterq-dump function. The raw read data is in the form of .`fastq` files. For data stored on AWS we use to AWS CLI to copy .`fastq` files to a local drive.

#### Reference genome

We download reference genome files from https://www.ensembl.org/ or https://www.gencodegenes.org/. Specifically, we download a *gene annotation file* .`gtf` and a *DNA sequence file* .`fa` for each organism. For each organism, we use the following versions.

**Table 1:**
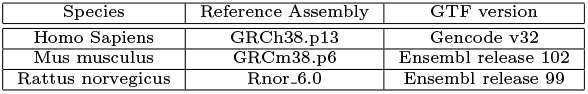
Reference Genome Versions

### Alignment, demultiplexing, and UMI counting

To align raw reads to a reference genome, we use the STAR aligner (version 2.7.5c) (*63*). Specifically, for multiplexed data we use STARsolo, which takes as input the fastq files containing all cells and returns (1) aligned reads in a `.bam` file and also (2) a UMI counts matrix `.tsv`. For non-multiplexed datasets (such as Smart-Seq2) STARsolo appends the sample name to the reads of each cells `.fastq` file and then creates an expression matrix for all cells simultaneously, as well as a `.bam` file which can then be split by sample name for variant calling on each cell.

For full length datasets, we constructed an expression matrix for the whole experiment with STARsolo, and we created the aligned `.bam` file using STAR for each cell separately. We processed CEL-seq2 `.fastq` files using UMI-Tools (version 1.0.1) (*64*). Cellular barcodes of reads were extracted from barcode `.fastq` files and placed into the read names of genomic reads using umi tools extract. We then aligned reads to the reference genome using STAR with default settings, outputting an aligned `.bam` file. featureCounts was then used to tag reads with the gene they map to. We processed this `.bam` file by using the umi tools count function to produce an expression matrix. We used the expression matrix and aligned `.bam` files for DEG analysis and variant calling, respectively, in the downstream analysis.

### Variant calling

We called variants using a custom variant calling pipeline. We attempt to call variants on all cells passing the first round of quality control on the expression matrix. For these cells we import the aligned cellular `.bam` files using pysam (v0.15.3) (*65*) and search for mutations. To call mutations we first drop reads which are not uniquely aligned to the mitochondrial genome to avoid heteroplasmy calls caused by NUMTs in the reference genome, then we search the remaining reads to find mismatches between observed bases and the reference base. We only considered genome positions if over 200 reads align to the position with a base quality score above 30. Furthermore, we excluded cells from our analysis if the number of positions passing quality control fell outside of a log-normal distribution, or if the number of positions passing quality control was below 300, so as to exclude cells where we are unlikely to be able to detect any mutations. To calculate heteroplasmy (*h*_*i*_) of a base *i ∈* [*A, C, G, T*] we take the ratio of the number of reads assigned to that base (*N*_*i*_) to the total number of reads at that position:

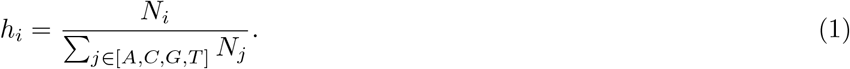

This definition of heteroplasmy allows for the possibility that two different mutations can occur at the same position on the genome in a cell, which occurs in less than 5 % of cells throughout our analysis. To aid comparison between the UMI and non-UMI data, we did not perform deduplication on any dataset. However, we find that when comparisons between heteroplasmy calls are made with and without deduplication on UMI data, strong agreement was observed (see Supplementary Discussion S6.3). Then we classified mutations found in only one cell of a donor/sample at *h*_*i*_ *>* 5 % as ‘cryptic’. We classified any mutations which were common to more than three donors from a given dataset as ‘common mutations’ and excluded them from further analysis to avoid common RNA variants being used in further analysis. We also exclude any mutation on a site that was not covered with sufficient depth in at least ten cells from a donor (thereby excluding sites with systematically small numbers of reads). After classification we keep only mutations with *h*_*i*_ *>* 10 % to exclude possible PCR or sequencing errors (see Supplementary Discussion S6). For rat sequencing data this was done at 5% to enable greater variant discovery due to the much lower coverage of the mitochondrial genome of that data set (see Supplementary Figure S20).

### Mitochondrial information

For human cells, we use the HmtVar (*66*) database to characterise mtDNA mutations. Specifically, we identify whether mutations result in a non-synonymous amino acid substitution, and classify their pathogenicity by using the MutPred score (*67*). MutPred is based on the effect of the substitution on protein structure and function, such that it categorises a non-synonomous mutations as either ‘low pathogenicity’ or ‘high pathogenicity’. Combing this information, we obtain three categories of human mutations: ‘synonymous’, ‘low pathogenicity’, and ‘high pathogenicity’. For mouse cells pathology scoring are unavailable for the majority of mitochondrial encoded proteins, so we classify mutations as either ‘synonymous’ or ‘non-synonymous’.

### Mitochondrial mutation statistics

To quantify the mitochondrial mutation of single cells, we compute different statistics. Given a cell with *m* mutations at heteroplasmies **h** = (*h*_1_, *h*_2_, …, *h*_*m*_) with *h*_*j*_ *∈* [0, 1], we define the *mitochondrial load* as:

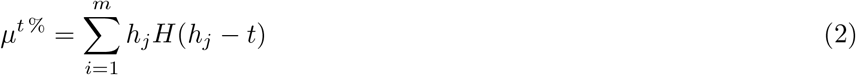

where *H*(*h*_*j*_ *− t*) indicates the Heaviside step function, such that only heteroplasmies greater than the threshold *t* contribute to the mutation load. By default, we only count cryptic mutations, which are not synonymous, above 10 % and indicate this for simplicity by *μ*.

### Quality control

Using the count matrix produced from STARsolo, we exclude cells as recommended by current best practices (*68*). For the analysis of the expression matrices, we use scanpy. We filter each expression matrix using three covariates: the total counts per cell, total genes per cell, and the percentage of reads aligned to the mitochondrial genome. These quality covariates are examined for outlier peaks that are filtered out by thresholding. We determined these threshold separately for each data set to account for quality differences. This quality control procedure allows us to establish cells with unexpectedly low read depth or high fraction of mitochondrial content, indicating that mRNA leaked from the cell during membrane permeabilization, or those with high read depth which could be doublets. We only keep cells, which pass both filtering steps, (1) the variant calling filtering (as discussed in subsection ‘Variant calling’) and (2) the expression matrix filtering (as discussed in this subsection).

### Gene expression analysis

We use scanpy for single cell gene expression analysis (*69*) and perform standard preprocessing steps. In addition to the filtering (as outlined in the ‘Quality Control’ subsection), we normalised to 10,000 reads per cell and log-transformed the counts. We use a Wilcoxon rank-sum test with significance threshold of 0.05 for DEG discovery and use the Benjamini– Hochberg procedure to obtain multiple-testing corrected *p*-values. For gene-ontology enrichment and pathway analysis, we use PANTHER (*70*). To combine the *p*-value of differential expression with a protein interaction network, we use scPPIN (*41*) and visualise it with Netwulf (*71*). For bulk sequencing data pathway analyses were performed using the WEB-based GEne SeT AnaLysis Toolkit following their instructions online.

### Mitochondrial bioenergetics characterization

Oxygen consumption was performed as previously described (*72*). For oxygen-consumption modifications, briefly, 20 × 104 cells per well were seeded 8–12 h before measuring basal respiration, leaking respiration, maximal respiratory capacity, and non-mitochondrial respiration (NMR). Respiration levels were determined by adding 1 *μ*M oligomycin (leaking respiration), 0.75 and 1.5 *μ*M carbonyl cyanide-p-trifluoromethoxyphenylhydrazone (maximal respiratory capacity) and 1 *μ*M rotenone–antimycin (NMR), respectively. Data were corrected with values from NMR and expressed as pmol of oxygen per min per mg protein.

### Determination of mitochondrial inner membrane potential and cytoplasmic ROS

Determination of mitochondrial inner membrane potential (MIMP) and cytosolic ROS were performed as previously described (*72*). Determination of MIMP was carried out using Tetramethylrhodamine, methyl ester at 20 nM in DMSO in parallel with mitochondrial mass detection using MitoTracker Green (20 nM in DMSO). Cytosolic ROS levels were measured using 2′,7′-dichlorofluorescin diacetate at 9 *μ*M in DMSO. All reagents were purchased from Invitrogen. Fluorescence-activated detection was carried using a BD LSRFortessa cell analyzer from BD. A total of 20,000 events were recorded, and doublet discrimination was carried out using FSC-Height and Area in the FlowJo software.

### Determination of ATP levels

ATP levels were determined with previously established methods (*73*). ATP levels were measured four times in three independent experiments using the CellTiter-Glo Luminescent Cell Viability Assay (Promega) according to the manufacturer’s instructions. Briefly, 10,000 cells per well were seeded, and the medium was changed 48 h before measurement. After that time, cells were lysed, and lysates were incubated with luciferin and luciferase reagents. Samples were measured using a NovoStar MBG Labtech microplate luminometer, and results correspond to the protein quantity measured in a parallel plate.

### Electrophoresis and western blot analysis

Electrophoresis and WB analysis was performed as previously described (*72*). Total protein extracts were prepared according to each protein’s solubility. Mitochondrial proteins were prepared using 2 % dodecyl-maltoside in PBS including protease inhibitors. Protein extracted for kinase phosphorylation analysis was extracted using the PathScan Sandwich ELISA Lysis buffer from Cell Signaling. In any case, protein extracts were loaded on NuPAGE Bis-Tris Precast mini/midi Protein gels with MES (Invitrogen. Electrophoresis was carried out following the manufacturer’s instructions. The SeeBlue Plus2 Pre-stained Protein Standard from Invitrogen was used in each electrophoresis as protein size markers. Separated proteins were transferred to polyvinylidene fluoride membranes. The resulting blots were probed overnight at 4^*°*^C with primary antibodies at the appropriate concentration following the manufacturer’s instructions with minor adaptations. The following antibodies were used; anti-EIF2A, Cell signaling #9722, 1:500; p. anti-p.EIF2ASer51, Cell signaling #972, 1:500, anti-4EBP1, Abcam ab2606, 1:500; anti-p.4EBP1Thr37&46, Cell signaling #9459, 1:500; anti-*β*-actin, Sigma-Aldrich A5441, 1:2000. After the primary antibody, blots were incubated for 1 h with secondary antibodies conjugated with horseradish peroxidase, and signals were immunodetected using an Amersham Imager 600 and/or medical X-Ray Film blu (Agfa). The bands for each antibody were quantified, aligned and cropped using the Fiji ImageJ 2.3.0/1.53q program, and the O.D. was used as a value for statistical purposes. To avoid interblot variation, one cell line was used as an internal control, and O.D. values corrected for *β*-actin levels were shown as relative to the internal control in each case.

### Rank biserial correlation difference

The rank biserial correlation difference, *r*, is a rank correlation. To illustrate its interpretation, consider 2 sets of heteroplasmic mtDNA mutations, called *G*_1_ and *G*_2_. State the hypothesis that mutations in *G*_1_ are greater in heteroplasmy than mutations in *G*_2_. r is an effect size measuring the degree of support for this hypothesis by considering all possible pairings between the mutations in *G*_1_ to the mutations in *G*_2_. Let f be the proportion of all such pairs of mutations in which mutations in *G*_1_ have a greater heteroplasmy than mutations from *G*_2_. Let u be the proportion of pairings in which *G*_2_ mutations have a greater heteroplasmy than the mutation from *G*_1_. The rank biserial correlation is simply defined as r = f - u, and can range from -1 to 1. *r* = *−*1 showing all mutations from *G*_2_ have a higher heteroplasmy than mutations from *G*_1_, negating the hypothesis. *r* = 1, indicates a positive effect size for the hypothesis indicating that all mutations in *G*_1_ are greater in heteroplasmy than those in *G*_2_. *r* = 0 indicates the mutations in *G*_1_ are observed to have a lower heteroplasmy than *G*_2_ mutations as often as they to have a higher heteroplasmy (*74*).

### The Moran model

The theory that we use to model mtDNA population in a post-mitotic cell is a fixed population size birth–death model with mutation known as the Moran model (*75*): it is arguably one of the simplest models that could be chosen. We consider that, at the start of the dynamics, no mutations (unique to each cell) are present in the system: as such, the site frequency spectrum of the system can be seen as out of equilibrium. In order for the population to remain constant at *N*, birth and death events are linked such that every time a randomly chosen mtDNA is replicated, another is randomly chosen to die (the same mtDNA can replicate and then die). As the half-life of mtDNA is much longer than the time taken for replication, we let the time between birth–death events be distributed as 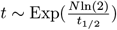, where *t*_1*/*2_ is the half life of mtDNA. During every replication there is a chance that *m* mutations occur along the length of the replicated mtDNA *L*_mtDNA_ due to errors from POLG, and we model this using a binomial distribution of the number of errors *m ∼* Binomial (*L*_mtDNA_, *ν*) where *ν* is the probability of mutation per base of POLG. Through this model, any mutation which enters the system can either be lost, or spread to higher heteroplasmies until it fixes through mtDNA turnover. We make the simplifying assumption that every mutation is unique, and neglect the possibility of back mutation. As the system initially has no mutations present, the mean heteroplasmy of a randomly chosen mutation increases with age. To move beyond a forwards in time simulation of the cellular population of mtDNA we make use of an equivalent backwards in time process known as the Kingman coalescent (*76*). Full details of this theory, including discussion of how the theoretical site frequency spectrum relates to our observed cryptic site frequency spectrum, are given in Supplementary Discussion 1.

## Acknowledgement

We thank the systems and signals group at Imperial College London for discussions.

## Author contributions

NSJ conceived and designed the study. AG, FK, ASM, JA and NSJ analysed the data. AG and NSJ developed theory. All authors interpreted the findings and wrote the manuscript. AG and FK contributed equally.

## Data availability

All analysed data is publicly available from the Gene Expression Omnibus (GEO) website or Amazon Web Services (AWS). See Table 2 for accession codes.

**Table 2:** Summary of the public data sets analysed in this study. Overall, we analyse sequencing data of more than 120,000 cells in three different species and five different tissues, as well as, cell-line data.

| Reference | Species | Tissues | Age range | No. of cells | Accession number | Protocol |
| --- | --- | --- | --- | --- | --- | --- |
| M. Enge, *et al.* (29) | Human | Pancreas | 1 mo-54 yrs | 2,544 | GSE81547 | Smart-seq2 |
| M.J. Muraro, *et al.* (30) | Human | Pancreas | 23-59 yrs | 12,200 | GSE85241 | CEL-seq2 |
| A.P. Voigt, *et al.* (31) | Human | Retinal Epithelium and Choroid | 54-92 yrs | 8,217 | GSE135922 | 10x |
| N. Almanzar, *et al.* (32) | Mouse | Pancreas, Liver | 3-24 mo | 9,380 | AWS | Smart-seq2 |
| J. Buenrostro, *et al.* (35) | Human | Cell lines | N/A | 1,344 | GSE65360 | scATAC-seq |
| S. Ma, *et al.* (54) | Rat | Brown Adipose Tissue, Liver | 5-27 mo | 29,165 | GSE137869 | 10x |
| S. Smajić, *et al.* (58) | Human | Midbrain | 66-93 yrs | > 41, 000 | GSE157783 | 10x (single-nucleus) |
| A. Grubman, *et al.* (77) | Human | Entorhinal cortex | 67-91 yrs | 13,214 | GSE138852 | 10x (single-nucleus) |
| M.R. Corces, *et al.* (78) | Human | Brain tissue | 38-95 yrs | 4,835 | GSE147672 | 10x ATAC (single-nucleus) |
| J. Camunas-Soler, *et al.* (79) | Human | Pancreas | 23-72 yrs | 3789 | GSE124742 | Smart-seq2 |
| | | | | $\Sigma = 124, 242$ | | |

### Cell lines

Cell lines were grown in DMEM containing glucose (4.5 g l-1), pyruvate (0.11 g/l) and FBS (5 %) without antibiotics at 37^*°*^C with 5 % CO2. Four cell lines were used 2 controls and 2 cell lines carrying mtDNA complex I mutations. All cybrids were obtained from cybrid pools after the selection process (*73*). MtDNA sequences of all cell lines can be found in GenBank, and their mtDNA accession numbers are included in table S6. Replication of these sequences have been carried out in this work through analysis of the full mtDNA sequence from the RNA-seq data.

## Code availability

The custom code used for our analysis is available on GitHub at https://github.com/SystemsAndSignalsGroup/Mito-Ageing.

## Supplementary Information

## S1 Supplementary Discussion: Truncated coalescent theory

### S1.1 Moran model and coalescent theory

In this supplementary discussion, we construct a mathematical model that allows us to describe and predict the accumulation of somatic mtDNA mutations throughout ageing. We seek an analytic formula describing the expected time-evolution of the expected number of mutations at every heteroplasmy (the site frequency spectrum, SFS). We extend the typical notion of the SFS to include mutations at 100 % heteroplasmy (homoplasmy) using tools from population genetics (see (*S1*) for an introduction).

A cell’s mtDNA copy number is regulated by mitophagy and replication in order to satisfy cellular energy requirements (*S2*). A simple forwards-in-time model for the mtDNA population of a single post-mitotic cell is a birth–death model with mutation assuming a fixed population size (*S3*). Crucially, in the following, we consider the scenario that, at some start time, *W*, no mutations (unique to that cell) are present in the system: a model for post-mitotic *de novo* mutation. This means that mutations gradually accumulate in the cell and the SFS, is out of equilibrium, and evolves from its form at start-time *W* to the present. This means that mutations gradually accumulate in the cell and the SFS, is out of equilibrium, and evolves from its form at start-time *W* to the present. In the Moran model, birth–death events, hereafter Moran events, are linked so that at all times the population size *N* remains fixed, and every time an event occurs one copy of the mtDNA is randomly chosen for replication and another is randomly chosen for death (Fig. S1a). As replication can occur on a much shorter timescale, compared to the half-life of mtDNA, we let the exponential rate of Moran events be 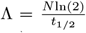, where *t*_1*/*2_ is the half life of mtDNA. During replication of an mtDNA, we assign a probability of mutation *P* (mut) = *u* which can be linked to the error rate *ν* per replication per base of POLG by assuming a mutation can occur during replication on any base with equal probability. Under this assumption the number *m* of mutations during replication of mtDNA with *B*_mtDNA_ bases is binomially distributed, *m ∼* Binomial (*B*_mtDNA_, *ν*), and so yields a probability of an mtDNA being mutated after replication *u* = 1 *− P* (*m* = 0). For a sufficiently small base mutation rate *ν, u* = *νB*_mtDNA_.

By using the Moran model as our forward-in-time model for mtDNA in a single cell, we can formulate a backwards-in-time equivalent process based on the *Kingman coalescent* (*S4*) to derive the properties of an out of equilibrium SFS. The Kingman coalescent focuses on deriving the tree structure which relates a sample of *n* individuals from a population *N*. Consider a single Moran event, one member of the population of *n* individuals is chosen to replicate producing two offspring, and another to die (this can be the same member). The only way for a pair of lineages to coalesce is if they both belong to the two offspring of the Moran event, and if we randomly select two members of the population just after this event, the probability that both are offspring is 2*/N* ^2^ giving us the coalescence probability. Extending this to a sample of *k* individuals, the probability that any two of them are offspring of the event is *k*(*k −* 1)*/N* ^2^. Using the time distribution between Moran events, multiplied by the probability that a Moran event resulted in a coalescence tell us that the time it takes *k* lineages to coalesce to *k −* 1 lineages is exponentially distributed with a rate of Λ*k*(*k −* 1)*/N* ^2^ where Λ is the rate Moran events. It is typical in coalescent theory to re-scale time such that the coalescence rate of *k* lineages is 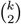, and we work in this re-scaled time when working with the coalescent throughout this derivation.

We wish to produce a theory describing the evolution of the SFS over time supposing that cells initially have no mutations present. We do this by adapting the standard Kingman coalescent with mutation in the following way:

- The coalescent process runs as standard for times *t < W*, and
- At time *t* = *W* all remaining coalescence occurs.

This modification is the same as assuming that for all times *t ≥ W*, there is no turnover in a population of identical DNA sequences, only for a Moran process with mutation to switch on at *t* = *W* and begin introducing mutations into the population. It is also equivalent analytically to the case where there are Moran dynamics, but zero mutation rate happening at *t ≥ W* (Fig. S1a-e).. We call this modified coalescent the *W-Truncated coalescent*, and any quantities with a superscript *W* refer to this process, while quantities without a superscript are results from standard coalescent theory. To convert any of the following results into proper time *τ*, we substitute 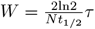.

**Supplementary Figure S1:**
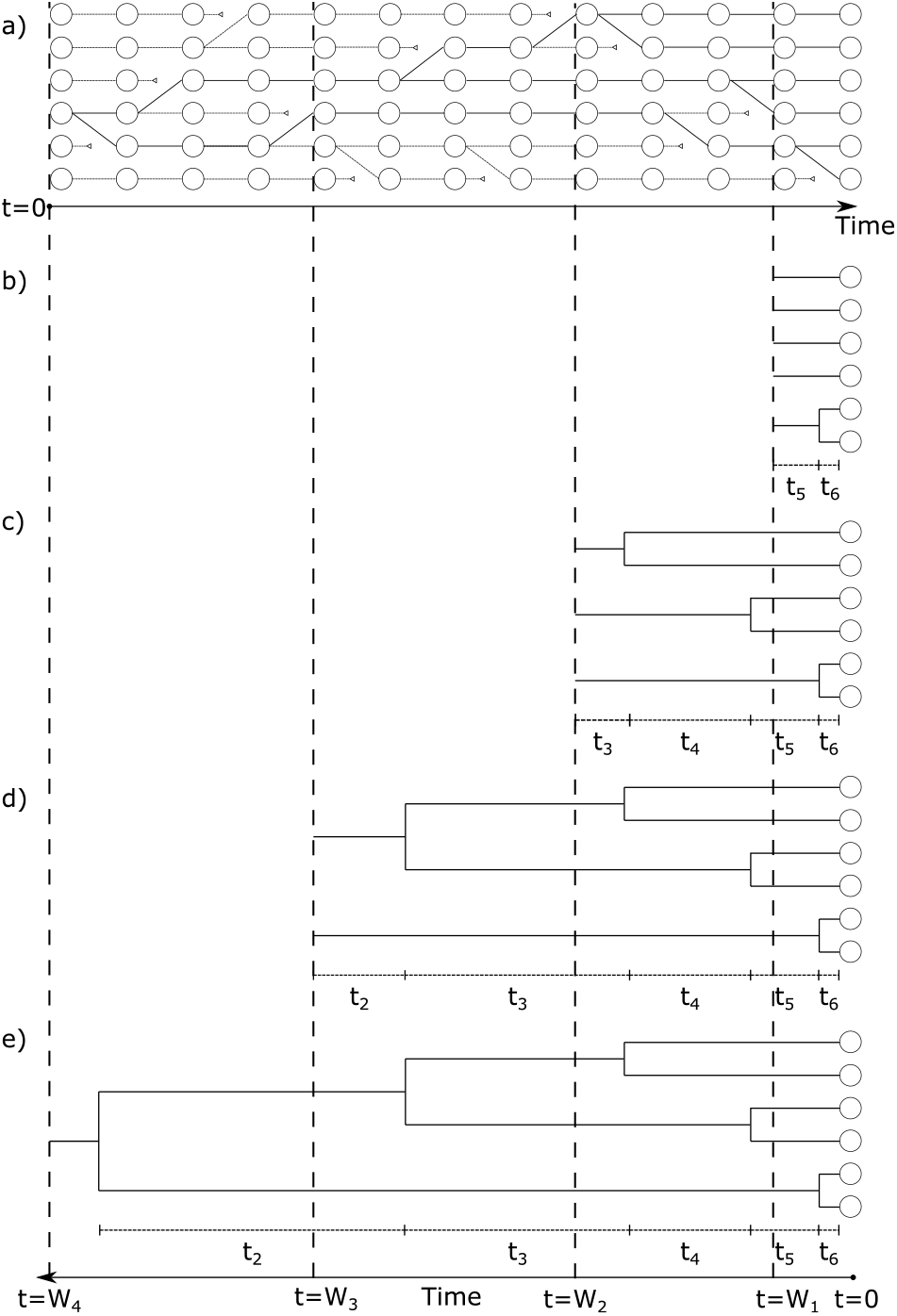
The dynamics of the W-truncated coalescent. (a) A Moran process where at every time step one member of the population is chosen to replicate and another chosen to die. (b-e) A Moran process can run for an arbitrary period of time, but for the W-truncated coalescent we are only interested in the dynamics backwards-in-time up to a time *W*. When working backwards in time we call the length of time the Moran process was allowed to run for the ‘truncation time’, as it is at this point we stop all remaining dynamics of the coalescent process. By increasing the truncation time *W* from *W*_1_ to *W*_4_, we look progressively further backwards in time along the Moran process shown in (a). The resulting coalescent trees drawn from samples at different time points are clearly different, with earlier samples resulting in shorter coalescent trees with coalescent times *t*_*k*_ dependent on the truncation time *W*. At the very short *W* s shown in (b-d), not enough time has passed for all coalescent events to have occurred, and we see that for shorter *W* s there are far fewer coalescent times. Increasing *W* further in (e) has allowed all coalescent events to have occurred, and only after this has happened can homoplasmic mutations begin to be seen in the sampled population.

To find the form of the SFS we need to first find the distribution of coalescence times in the *W-Truncated coalescent*. We do this through finding 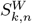, the random variable giving the time at which *k* lineages coalesce in the *W-Truncated coalescent* process with a sample size of *n*. From this we can find the coalescence times using the expectation of the difference between time *k* and *k* + 1 lineages coalesce: 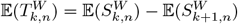.

### S1.2 Distribution of 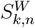

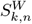 is distributed as in the conventional coalescent (the random variable *S*_*k,n*_) for *t < W* but for any *t ≥ W* all coalescence happens at *W*.

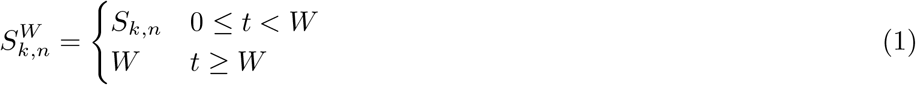

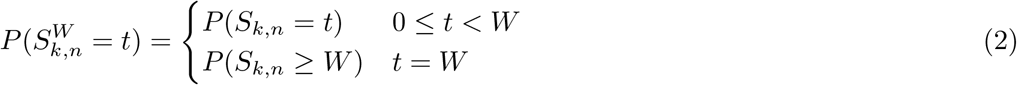

*S*_*k,n*_ is hypo-exponentially distributed so:

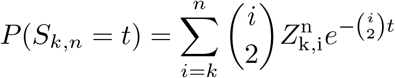

where

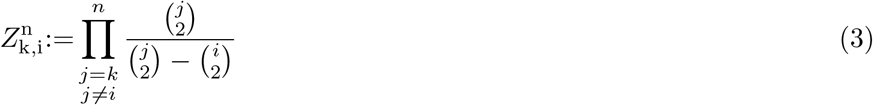

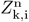 has the identities:

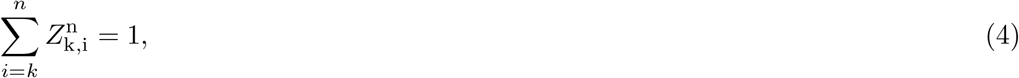

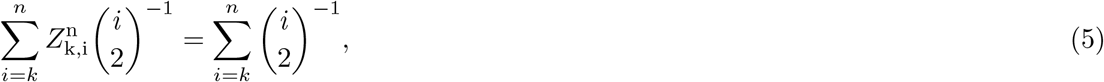

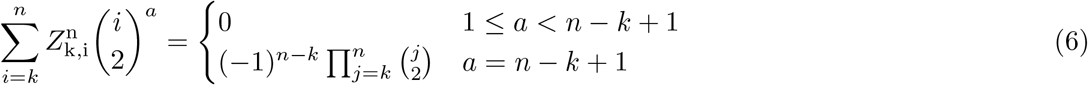

Continuing the derivation, we look for *P* (*S*_*k,n*_ *≥ W*):

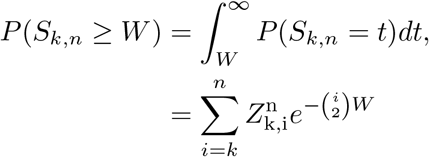

This gives us the distribution of times to *k*^*th*^ coalescence as:

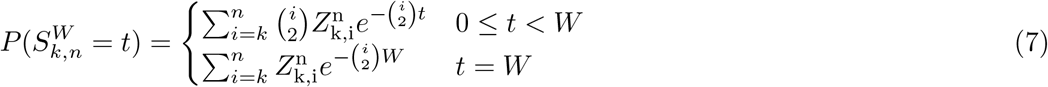

### S1.3 Expectation of 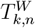

The difference between the time *k* and *k* + 1 lineages coalesce is defined as 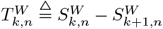

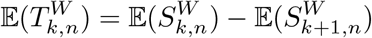

We first compute the expectation of the distribution of times to *k*^*th*^ coalescence as

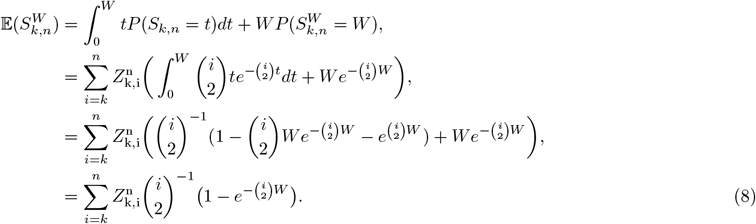

Eq. 8 can be simplified further using Eq. 5 leaving:

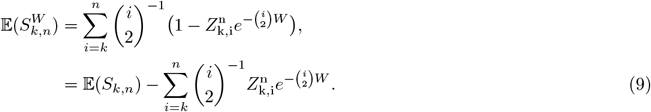

From Eq. 9 follows the anticipated behaviour in the limits of *W*

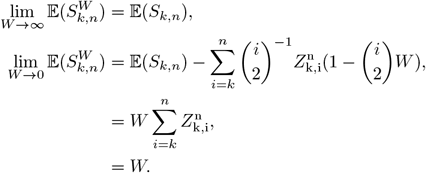

As we allow *W → ∞* the *W-Truncated coalescent* converges to results expected under standard coalescent theory, and when *W →* 0 the time to *k*^*th*^ coalescence is limited by W.

We now expand 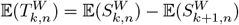 and obtain

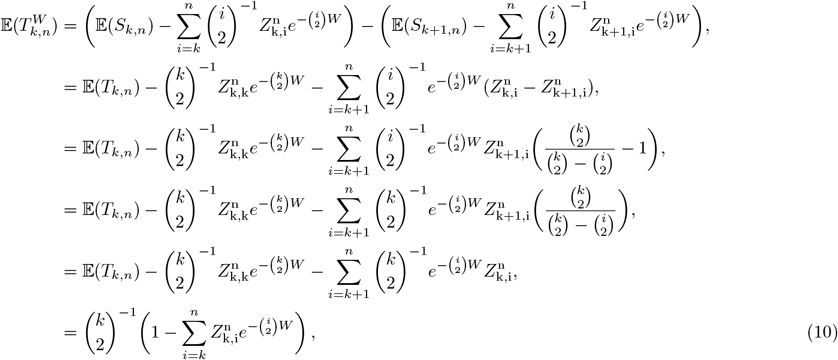

where we use the result from conventional coalescent theory that 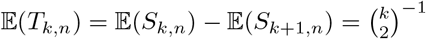

### S1.4 Expected branch lengths

We aim to get a distribution of the expected number of mutations found at a given heteroplasmy. To do this we first need to find the expected branch length of all branches in the tree which carry *b* descendants 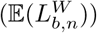. At a level with *k* lineages, we wish to know the probability that any one of those lineages will carry *b* descendants where 1 *≤ b ≤ n − k* + 1. To do this we consider the vector of the number of descendants of the *k* lineage, 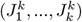 with 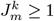. This vector is uniformly distributed over all vectors where 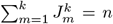. The number of such vectors is 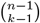. The number of these vectors for which a single lineage *m* carries *b* descendants is equal to the number of ways the remaining *n − b* descendants can be split into *k −* 1 non-empty groups, which is a known result from combinatorics of 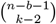. This leaves the probability of a lineage carrying *b* descendants 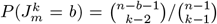. Given there are *k* lineages at this level, and the level is present for a time 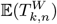, we find the contribution from all levels to the length of all branches carrying *b* descendants to be:

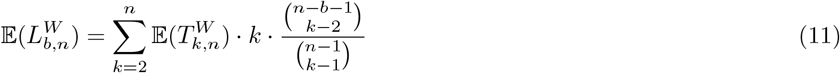

Substituting in the expected times to coalescence gives us a formula of

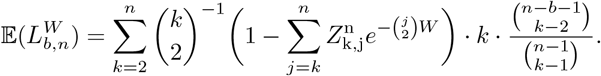

Using the identity

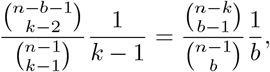

we can get the most simplified form of the 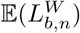:

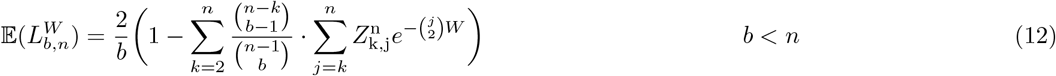

To find the extended site frequency spectrum we must identify the length of the ‘root’ of the coalescent tree 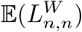. This is done by finding the difference between 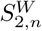 and *W*.

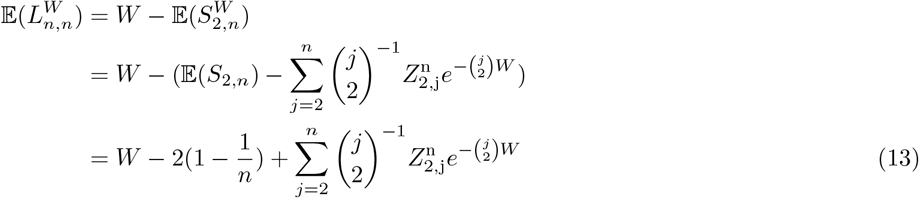

Combining these results we get:

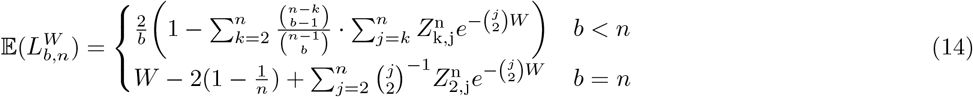

We see from equation 14 that while the expected branch length for heteroplasmic mutations reach a steady state as *W → ∞*, the equivalent for homoplasmic mutations will constantly increase as time goes on. This is because homoplasmic mutations have no avenue to be lost from the population due to the infinite sites assumption, whereas heteroplasmic mutations are either lost or fix at homoplasmy. This constant accumulation of homoplasmies is what allows us to use the extended site frequency spectrum to infer a tissue’s age when its heteroplasmies have already reached equilibrium.

### S1.5 Normalised extended site frequency spectrum

To find the full site frequency spectrum we first need to find the expected branch length of all branches in the tree which carry *i* descendants. Following the formula from Durrett Chapter 2 (Section 2.2) (*S1*) for 𝔼(*L*_*i,n*_) and substituting Eq. 10 we can find 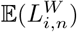 the expected branch length of all branches carrying *i* descendants in a sample of size *n* after time *W* (*i < n*).

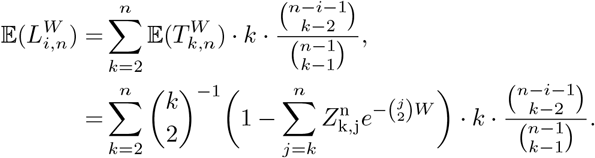

Using the identity:

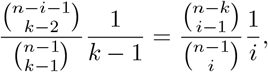

we can get the most simplified form of the 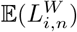:

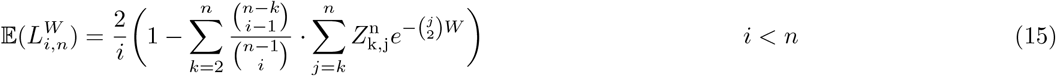

To find the extended site frequency spectrum we must identify the length of the ‘root’ of the coalescent tree 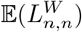. This is done by finding the difference between 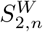 and *W*.

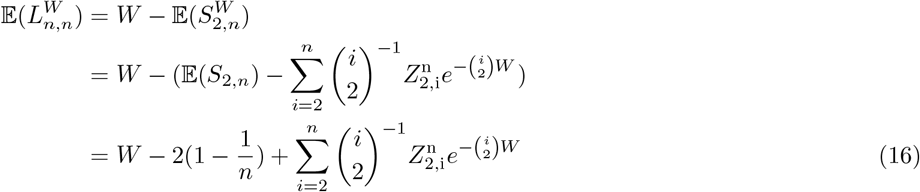

Combining these results we get:

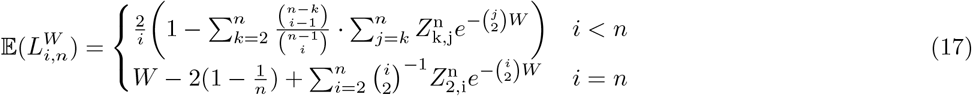

The normalized extended site frequency spectrum is then found using:

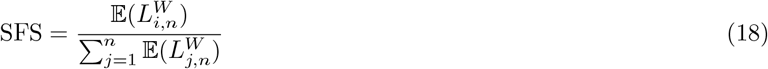

### S1.6 Distribution of number of mutations in a single cell

The distribution of the number of mutations *M* that occur on a given length of a coalescent tree, *L*, is distributed as a Poisson with rate *θL*, where *θ* = *Nu/*2. We wish to find the probability of a given number of mutations at a heteroplasmy *b/n, M*_*b,n*_ given the truncation time, *W*, and the mutation rate, *θ*, which can be found by integrating over the probability distribution of all potential branch lengths of the tree.

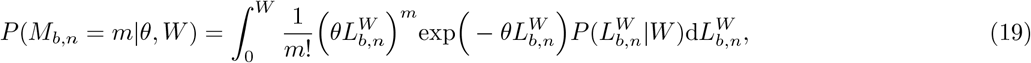

where 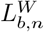 is the total branch length of the coalescent tree which carries *b* descendants. Doing this integral analytically is computationally intractable, so we turn to Monte-Carlo integration to approximate the integral. In brief, we sample *S* coalescent trees under the truncated coalescent process parameterized by *W*, take the lengths 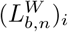 from these samples and evaluate:

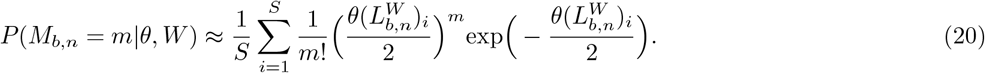

### S1.7 Multiple cells

Now that we have *P* (*M*_*b,n*_ = *m*|*θ, W*) for a single cell we can use this for our likelihood of a set of *C* cells carrying 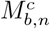 mutations each. If *C*_*m,b*_ is the number of cells with *m* mutations at heteroplasmy *b/n*, then:

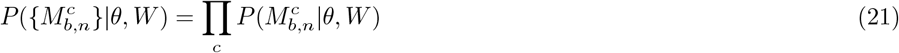

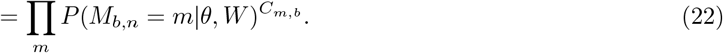

Looking at the number of mutations across all heteroplasmies we find the likelihood of data *D* to be:

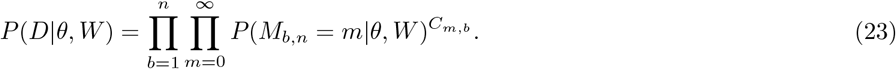

### S1.8 Hierarchical Bayesian Inference

In order to account for potential inter-individual differences in mtDNA copy number, turnover rate, and mutation rate, as well as the variable number of bases surveyed in each sequencing experiment, we apply a hierarchical model structure to our inference. The conversion between proper time and the truncation time *W* is 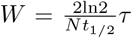. To account for differences between the copy number and mtDNA half-life of different individuals we will attempt to infer the mitochondrial ageing rate 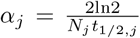 for each individual *j*, such that an individual of age *τ*_*j*_ has a mitochondrial age *W*_*j*_ = *α*_*j*_*τ*_*j*_. We must also account for differences in copy number, *N*_*j*_, mutation rate per base per replication, *ν*_*j*_, and number of bases surveyed *B*_*j*_ which will all contribute to the coalescent mutation rate *θ*_*j*_ = *N*_*j*_*ν*_*j*_*B*_*j*_*/*2. We will attempt to infer a parameter Θ_*j*_ = *N*_*j*_*ν*_*j*_*/*2 for each individual.

The hierarchical structure of the model will be as follows: each individual has two target inference parameters *α*_*j*_, Θ_*j*_ which are drawn from population distributions *α*_*j*_ *∼* LogNormal(*μ*_*α*_, *σ*_*α*_), Θ_*j*_ *∼* LogNormal(*μ*_Θ_, *σ*_Θ_). The parameters *μ*_*α*_ and *μ*_Θ_ are the expected values of *α*_*j*_ and Θ_*j*_, while *σ*_*α*_ and *σ*_Θ_ are the standard deviations of the logarithm of these two random variables.

For a set of *K* donors *X*_1_, …, *X*_*K*_, each with *C*_*d*_ cells each, our final joint posterior for the hyperparameters will be:

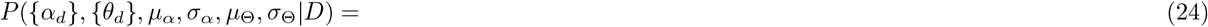

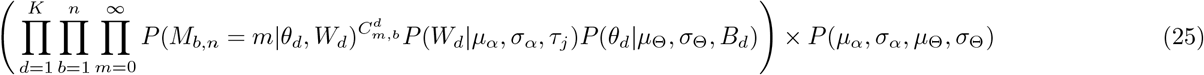

### S1.9 Comparison to the cryptic site frequency spectrum

The similarity between the cryptic mutations we identify from experiment and the model of *de novo* mutations that occur post-mitotically is affected by any replication those cells experience throughout an organisms life, as well as by tissue sampling effects. We limit ourselves to studying tissue types which undergo relatively slow turnover: the Enge dataset (*S5*) is made up from majority alpha, acinar, ductal, and beta cells, so as from (*S6*) we expect ductal cells to be largely non-proliferative, and alpha, acinar and beta cells to be 83 %, 24 % and 61 % non-proliferative respectively at the time points we consider. Ref. (*S6*) uses neurons as the reference slow tissue; we consider neural datasets (*S7* –*S9*). Our last dataset (*S10*) is taken from the retinal epithelium, which has cellular lifespan equivalent to that of neurons (*S6*).

In the event that a cell has undergone additional rounds of cellular replication (and yet has no sister cells contained within the sample) the cell will have a site frequency spectrum with a higher average heteroplasmy than a cell with the same mtDNA birth-death rate and no cellular replication: for the cell that replicates, the theory we outline above is still broadly applicable, but the interpretation of the parameters associated with turnover will be different.

As cryptic mutations are defined as those that occur in only one cell *within the dataset*, we could classify a mutation which occurs late in development or after a cell turnover event as cryptic if we only sample one cell in which this mutation occurs. Our sample-specific definition of cryptic also prevents us from considering post-mitotic *de novo* mutations which independently recur multiple times in an individual, however, in the datasets used through this paper *<* 1 % of mutations will be recurrent (simplifying and assuming the mutation probability is constant along the length of mtDNA).

## S2 Supplementary Discussion: Model inferences

### S2.1 Comparison to simulation

We now show that the model specified above can properly predict the mitochondrial age, *W*, and mutation rate, *θ*, from data generated using simulations of the Moran model using the parameters: *t*_1*/*2_ = 15 days (*S11, S12*), *ν* = 4 *×* 10^*−*8^ per base per replication (*S13*) and *N* = 1000 (*S14*). We do not initially impose a hierarchical structure on our model, as we first wish to verify that our simple likelihood can recapitulate simulation parameters. Simulating 300 different cells for different lengths of time, and sampling 200 mtDNA from each cell, we apply the model to the resulting SFS with a uniform prior for *W ∈* [0, 5] and *θ ∈* [10^*−*2^, 10^2^]. Given that when we analyse real data we filter our heteroplasmies below 10 %, we apply the same filter to this simulated data. This makes the final likelihood:

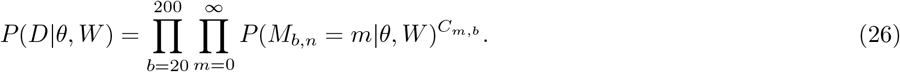

We see in tables S1 and S2 that the true *W* and *θ* lie inside the 95 % credible interval of the marginal posteriors inferred for *W* and *θ* for each time point. The full marginals for *W* and *θ* are shown in Fig. S2 and we also show the SFS predicted from the maximum a posteriori (MAP) estimate of *W* compared to the simulated (Figure S3). From this we can have confidence that our inference can credibly infer parameters from realistic amounts of data.

**Table S1:** The credible intervals and MAP estimate of *W* for different ages and their associated *W* values.

| Age Simulated (Years) | True $W$ Value | 95 % Credible Interval | MAP Estimate $W$ |
| --- | --- | --- | --- |
| 5 | 0.17 | (0.07, 0.18) | 0.12 |
| 10 | 0.34 | (0.20, 0.36) | 0.27 |
| 25 | 0.84 | (0.64, 0.86) | 0.78 |
| 50 | 1.68 | (1.5, 1.94) | 1.67 |
| 75 | 2.52 | (2.23, 2.77) | 2.49 |
| 100 | 3.36 | (3.32, 4.08) | 3.65 |

**Table S2:** The credible intervals and MAP estimate of *θ* for different ages and their associated *θ* value.

| Age Simulated (Years) | True $\theta$ Value | 95 % Credible Interval | MAP Estimate $\theta$ |
| --- | --- | --- | --- |
| 5 | 0.33 | (0.21, 2.04) | 0.58 |
| 10 | 0.33 | (0.27, 0.65) | 0.42 |
| 25 | 0.33 | (0.28, 0.41) | 0.34 |
| 50 | 0.33 | (0.28, 0.35) | 0.32 |
| 75 | 0.33 | (0.30, 0.38) | 0.35 |
| 100 | 0.33 | (0.28, 0.35) | 0.32 |

### S2.2 Hierarchical simulation comparison

To confirm that the parameters of the hierarchical model as outlined in section S1.8 can be recovered using inference, we now simulate 10 donors with different ages between 10 and 100 years old, with parameters *α*_*d*_ *∼* LogNormal(*μ*_*α*_, *σ*_*α*_), Θ_*d*_ *∼* LogNormal(*μ*_Θ_, *σ*_Θ_) where *μ*_*α*_ = 0.03, *σ*_*α*_ = 0.3, *μ*_Θ_ = 10^*−*5^, and *σ*_Θ_ = 0.5. Each donor had a random number of bases from the genome sampled, *B*_*d*_ *∈* Uniform(9000, 11000), and a random number of cells *C*_*d*_ *∈* Uniform(200, 400). We find that the model is able to infer the hyperparameters, as well as the individual donor parameters (see table S3, Fig. S4).

**Table S3:** The credible intervals and MAP estimate for the hyperparameters. The true hyperparameters are contained in the 95 % confidence interval in all cases.

| Parameter | True Value | 95 % Credible Interval | MAP Estimate |
| --- | --- | --- | --- |
| $\mu_\alpha$ | 0.03 | (0.028, 0.038) | 0.033 |
| $\sigma_\alpha$ | 0.3 | (0.06, 0.42) | 0.15 |
| $\mu_\Theta$ | $10^{-5}$ | $(6.61 \times 10^{-6}, 1.51 \times 10^{-5})$ | $9.55 \times 10^{-6}$ |
| $\sigma_\Theta$ | 0.5 | (0.28, 0.35) | 0.32 |

**Supplementary Figure S2:**
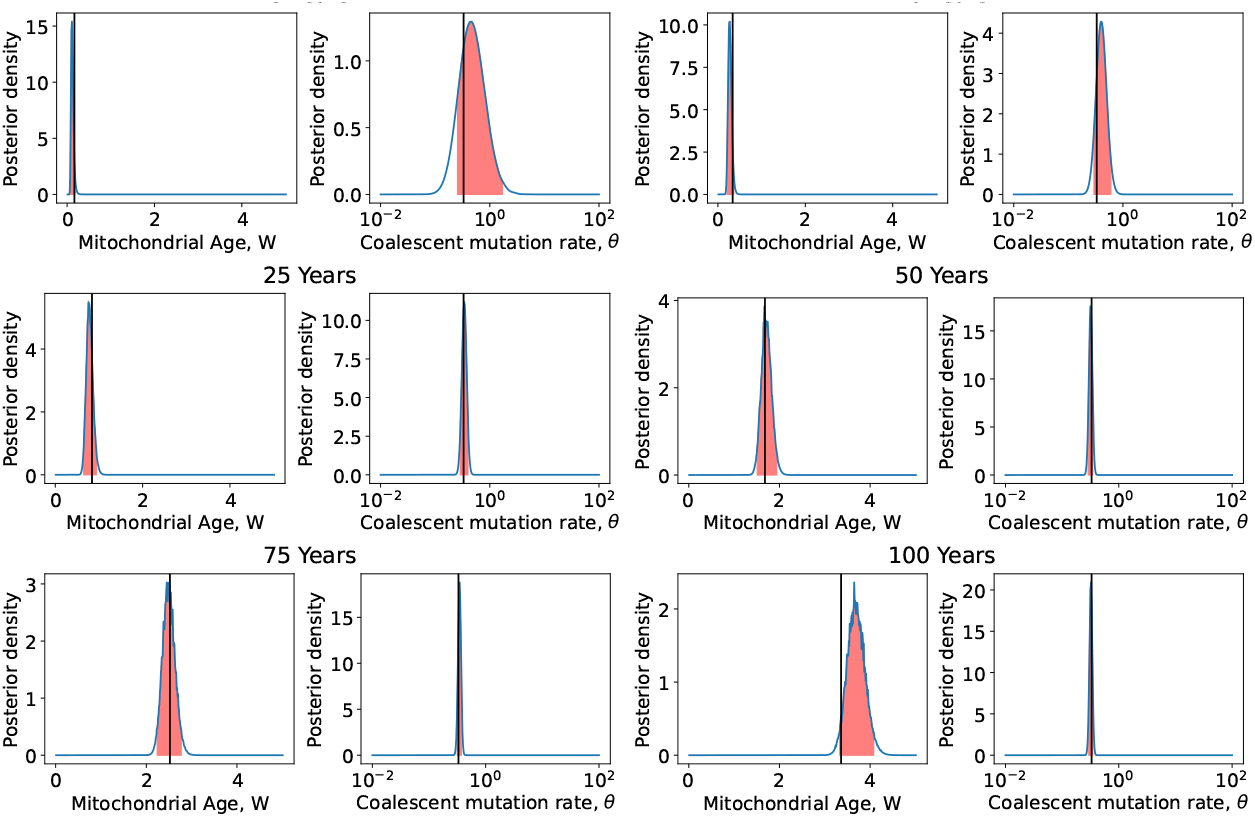
The posterior distribution of the mitochondrial age *W* and coalescent mutation rate *θ* found for each time point. Also shown with the black vertical line is the true values. We find for every donor that the true *W* and *θ* values lie inside the 95 % credible interval, shown in red.

**Supplementary Figure S3:**
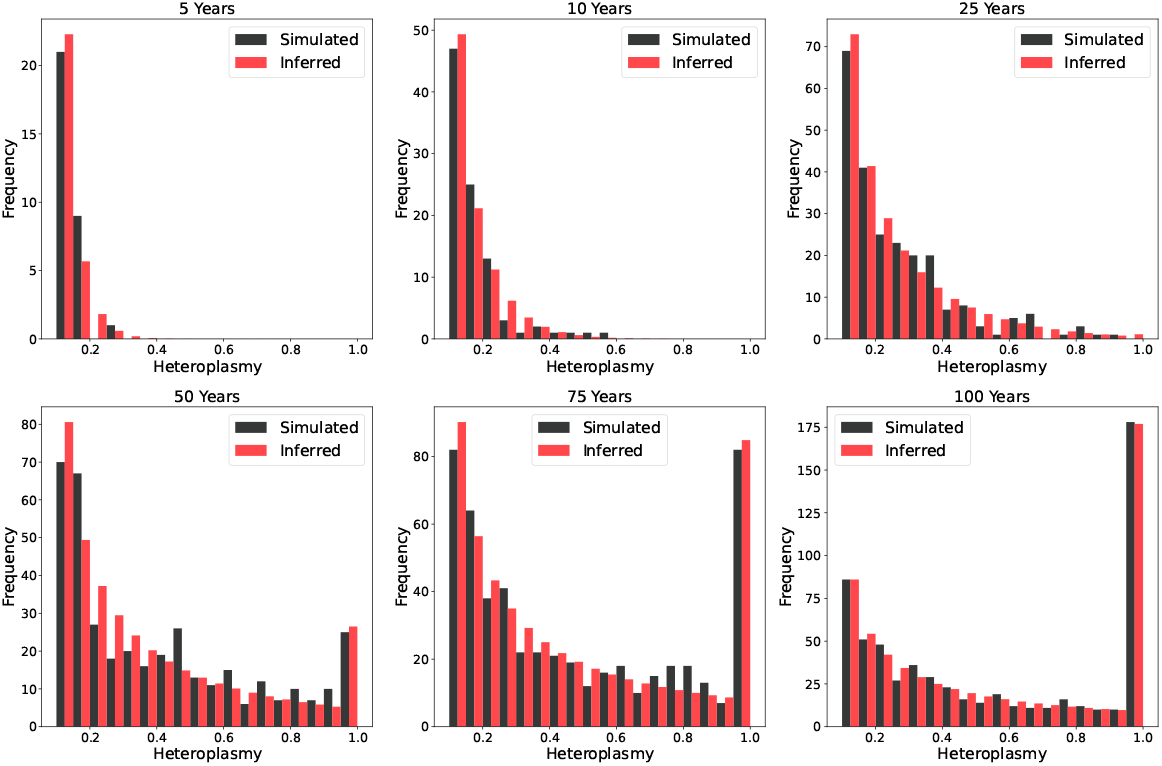
The predicted expected cSFS for the MAP estimates of *W* and *θ*, compared to the simulated cSFS for different ages.

**Supplementary Figure S4:**
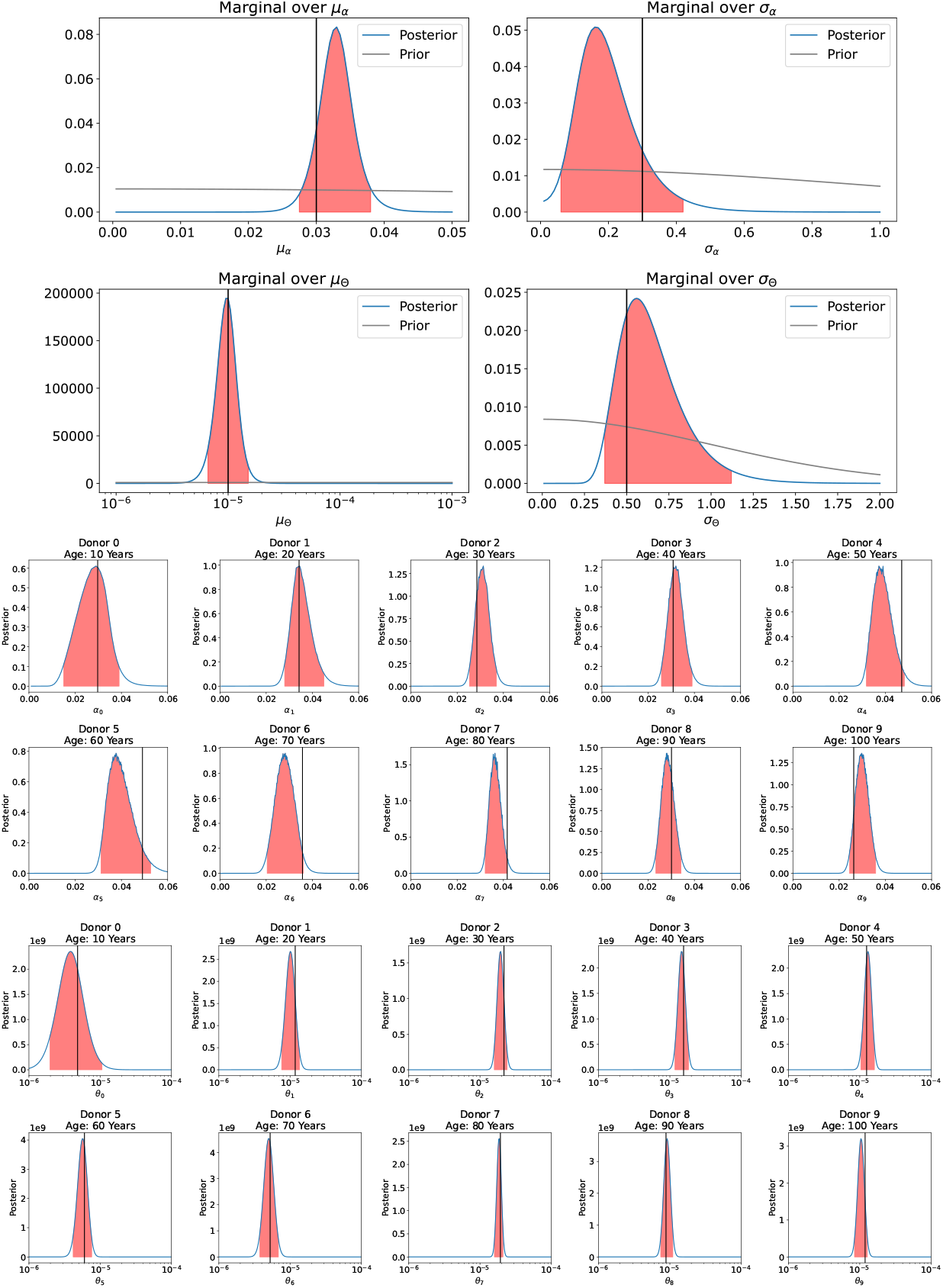
The marginal posterior distributions of the model parameters. The 95 % confidence intervals are marked in red, with the true parameter values shown with a black line. We see that the true value is contained by the 95 % confidence interval in all cases.

### S2.3 Model fitting to Data

We have established that the model is able to recapitulate parameters from simulated data accurately, and we now wish to apply it to our collected data. With this we have two goals -to show that mitochondrial age *W*, as inferred from the data, can be used as a marker of tissue age, and to find an estimate of the average mitochondrial mutation rate. We assume that all mutations which could be generated during sample and library preparation are found at heteroplasmies below 10% (see section S6.2), and that the sample size *n* should correlate with the average number of mtDNA a mutation is found from. Following the arguments from section S6.2 we expect there to be 7 reads for every true mtDNA sequenced. The average read depth of mutations is *≈* 640 reads and so we use *n* = 90 as the number of true mtDNA in our sample from the mtDNA population of the cell, though we find equivalent results for other values of *n*. In order to fit the data using our discrete likelihood, we round our heteroplasmy to values of 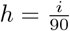 where *i ∈* [1,. ., 90], and then compute the full likelihood given all the data. The final likelihood form is:

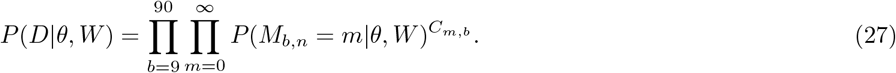

where *C*_*m,b*_ is the number of cells with *m* mutations at heteroplasmy 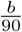.

### S2.4 Fitting to human data

For the human data we picked broad hyperprior distributions of *μ*_*α*_ *∼*Half-Normal(0.1), *μ*_Θ_ *∼*Half-Normal(0.01), *σ*_*α*_ *∼*Half-Normal(1), *σ*_Θ_ *∼*Half-Normal(1). Shown in Fig. 1h are the 95 % confidence intervals of the inferred mitochondrial age of every donor, which, as expected, increases with biological age. The fit shown on this plot is the MAP estimate of *μ*_*α*_ and its 95 % confidence interval. In Fig. 1i we show the marginal distribution of *μ*_Θ_, scaled by an estimate of mtDNA copy number, *N* = 1000 (*S14*) such that we are seeing the posterior of the median mutation rate per base per replication, *μ*_*ν*_. The full posteriors of both the hyperparameters and the donor specific parameters are shown in Figs. S5, S6, S7. We find that similar parameter values (though with broader posteriors) are recapitulated when the model is fit only to the most abundant cell type from the Enge pancreas dataset (*S5*) (see Fig. S23c).

Using our MAP estimates of *W* and *θ* for each donor, we can compare the expected site frequency spectrum for these values to the observed (Figure S8). We find that there is broad agreement between our expected site frequency spectra and those observed. Adding an explicit error model of sequencing and PCR to the likelihood, and allowing thresholds to vary based on the data quality would improve the inference, though the trends seen would likely remain unchanged. The fits presented in the main text for changes in relative probability of high heteroplasmy mutations (Fig. 1j) and for changes in the number of homoplasmic mutations (Fig. 1k-m) are done using the MAP estimates of *μ*_*α*_ and *μ*_Θ_, corresponding to the median of population distribution.

**Table S4:** The credible intervals and MAP estimate for the hyperparameters for all datasets. We see that mice have significantly higher mitochondrial ageing rate, *μ*_*α*_, and coalescent mutation rate, *μ*_Θ_.

| Parameter | Data used | MAP estimate | 95 % Credible Interval |
| --- | --- | --- | --- |
| $\mu_\alpha$ | All human (S5, S10, S15) | 0.036 | (0.029, 0.045) |
| $\mu_\alpha$ | Mouse liver (S16) | 0.456 | (0.300, 0.768) |
| $\mu_\alpha$ | Mouse pancreas (S16) | 0.252 | (0.156, 0.528) |
| $\mu_\Theta$ | All human (S5, S10, S15) | $2.29 \times 10^{-5}$ | $(1.38 \times 10^{-5}, 4.79 \times 10^{-5})$ |
| $\mu_\Theta$ | Mouse liver (S16) | $2.19 \times 10^{-4}$ | $(1.58 \times 10^{-4}, 3.31 \times 10^{-4})$ |
| $\mu_\Theta$ | Mouse pancreas (S16) | $1.51 \times 10^{-4}$ | $(9.55 \times 10^{-5}, 2.51 \times 10^{-4})$ |
| $\sigma_\alpha$ | All human (S5, S10, S15) | 0.42 | (0.33, 0.67) |
| $\sigma_\alpha$ | Mouse liver (S16) | 0.54 | (0.39, 1.14) |
| $\sigma_\alpha$ | Mouse pancreas (S16) | 0.67 | (0.49, 1.42) |
| $\sigma_\Theta$ | All human (S5, S10, S15) | 1.25 | (0.99, 1.82) |
| $\sigma_\Theta$ | Mouse liver (S16) | 0.34 | (0.22, 0.85) |
| $\sigma_\Theta$ | Mouse pancreas (S16) | 0.31 | (0.16, 1.28) |

**Supplementary Figure S5:**
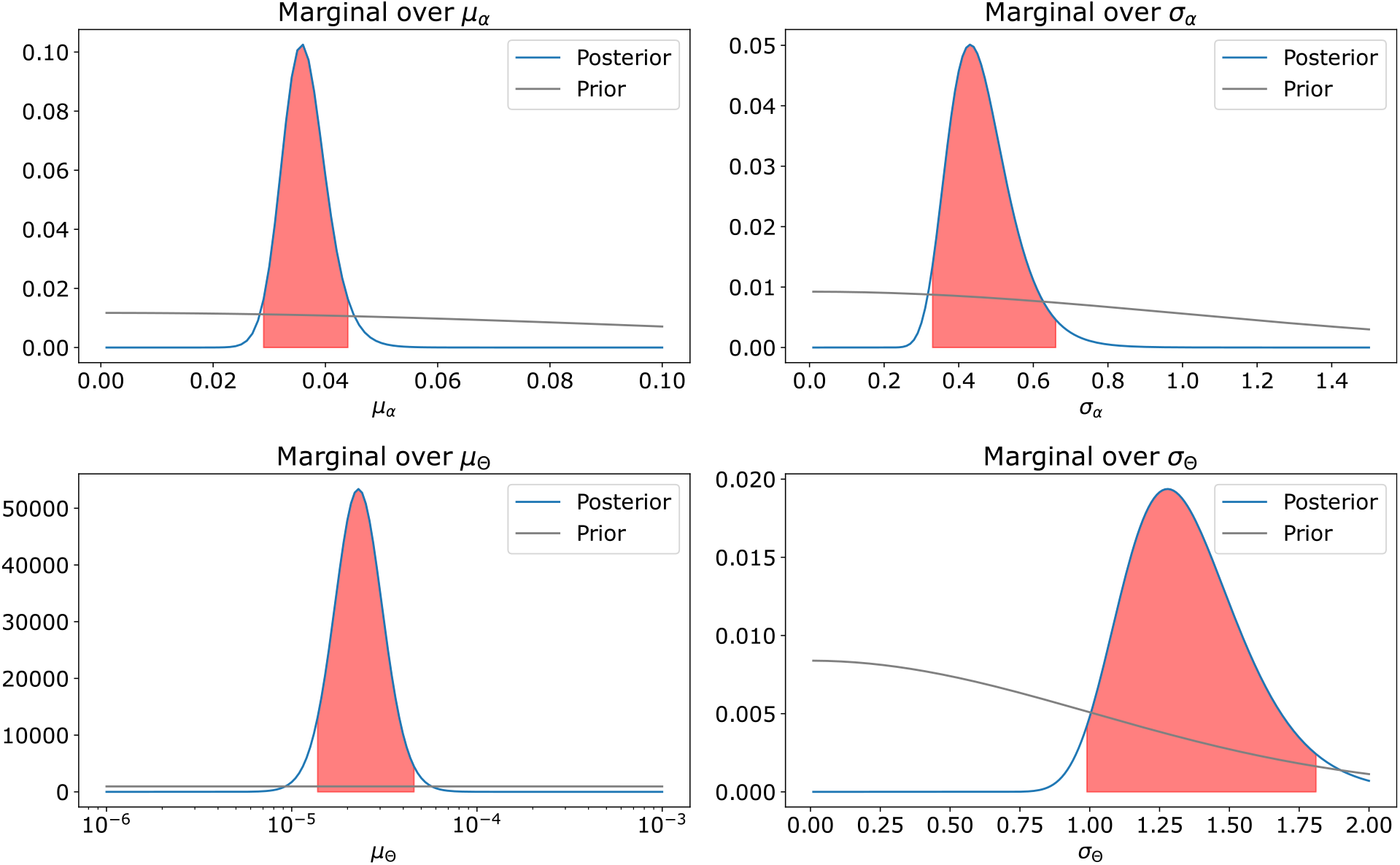
Unnormalised posteriors of the human hyperparameters. Marked in red are the 95 % confidence intervals. The large inferred variance on the mutation rate, *σ*_Θ_, is driven by the youngest donor’s large inferred mutation rate. When this donor is excluded most parameters remain the same, with *σ*_Θ_ reducing significantly (see Fig. S9 for full posteriors in this case).

**Supplementary Figure S6:**
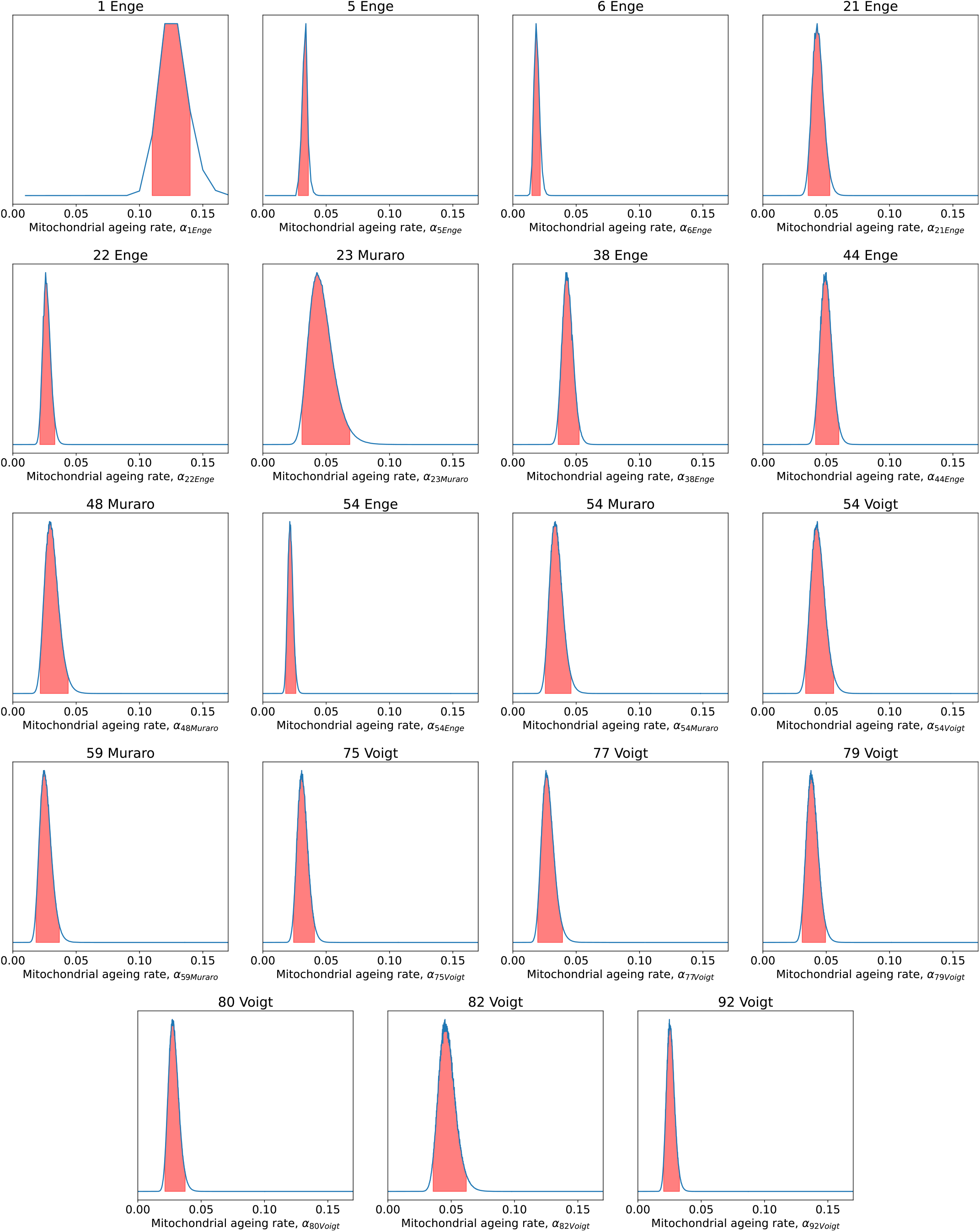
Unnormalised posteriors of the donor specific mitochondrial ageing rates. Marked in red are the 95 % confidence intervals. To convert these posteriors into effective mitochondrial ages (as seen in Fig. 1h, we multiply *α*_*d*_ by the donor age. Due to the grid evaluation of the posterior being over mitochondrial age, we see fewer evaluated points across the same range of *α* for younger donors.

**Supplementary Figure S7:**
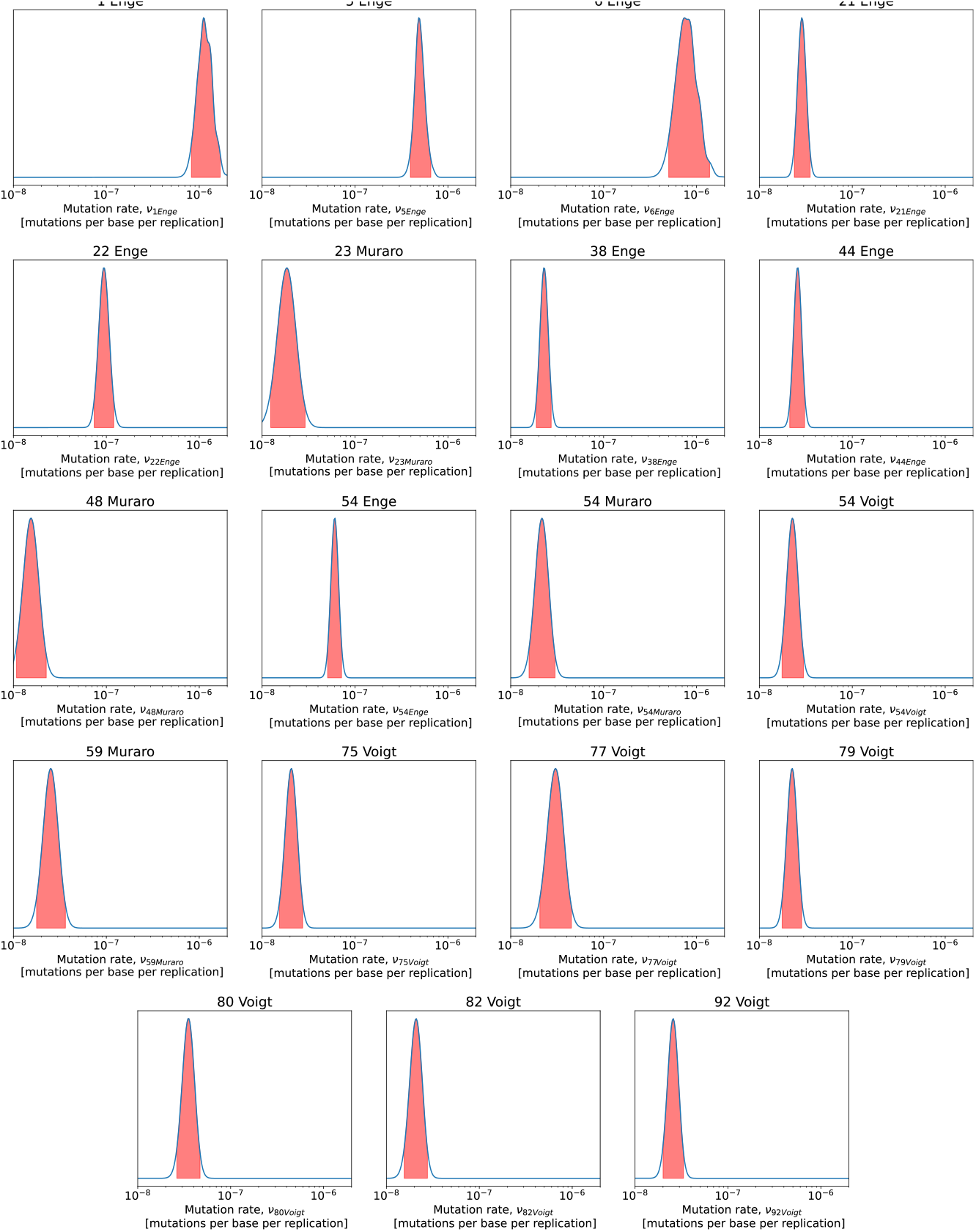
Unnormalised posteriors of the donor specific mutation rates. Marked in red are the 95 % confidence intervals. We see that the youngest 3 donors have much higher inferred mutation rates that the other donors. This could be due to low heteroplasmy mutations being dominated by errors from PCR, and very few high heteroplasmy mutations being present which can be used to compensate due to their young age.

**Supplementary Figure S8:**
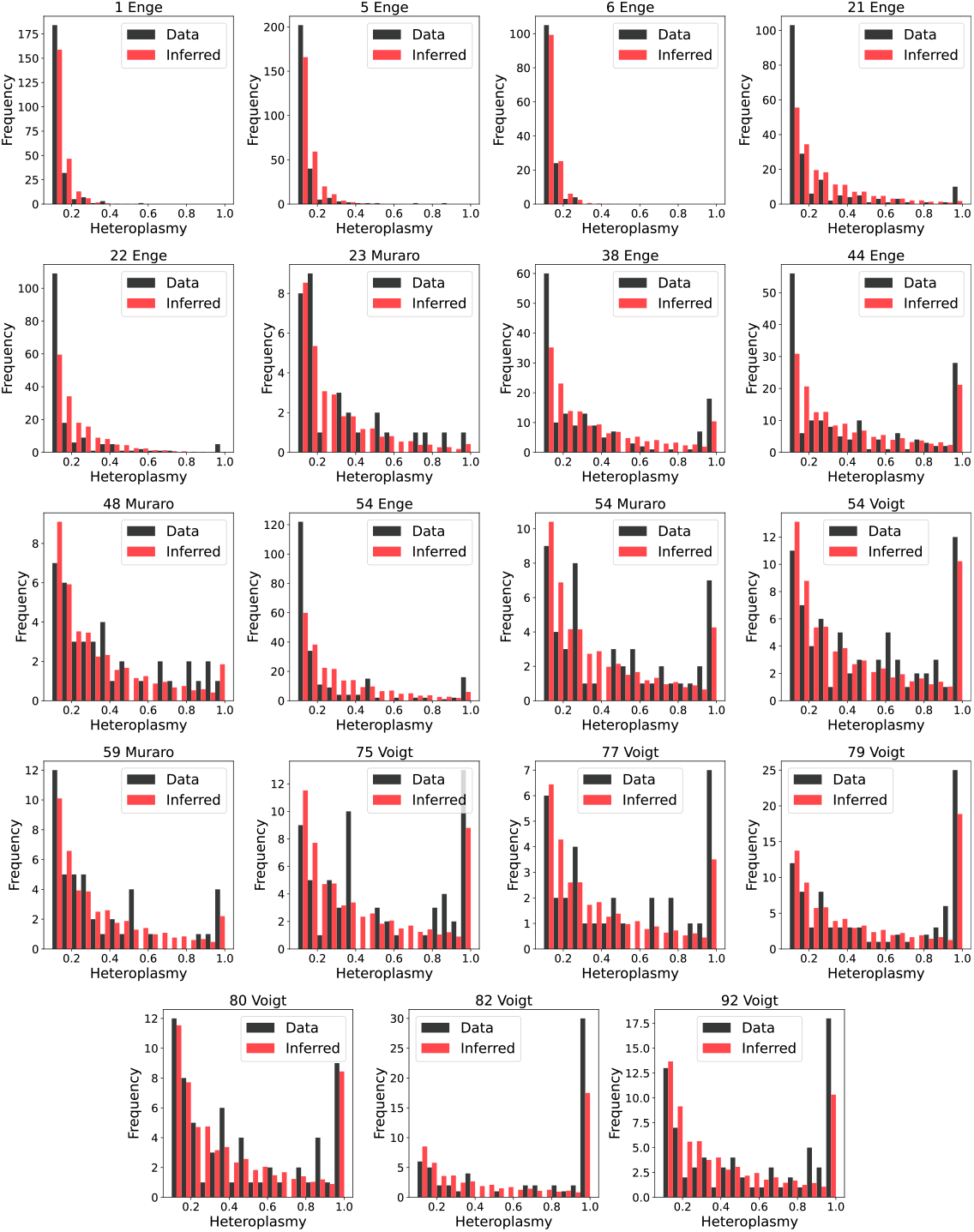
Comparison between the site frequency spectrum of our donors and the expected site frequency spectrum given our MAP estimate of W. The title of each subplot in the format *Age (Years) Dataset*. We see that the model performs better for Muraro and Voigt datasets, possibly due to the additional rounds of PCR in the high depth Enge dataset introducing more errors at low heteroplasmies.

**Supplementary Figure S9:**
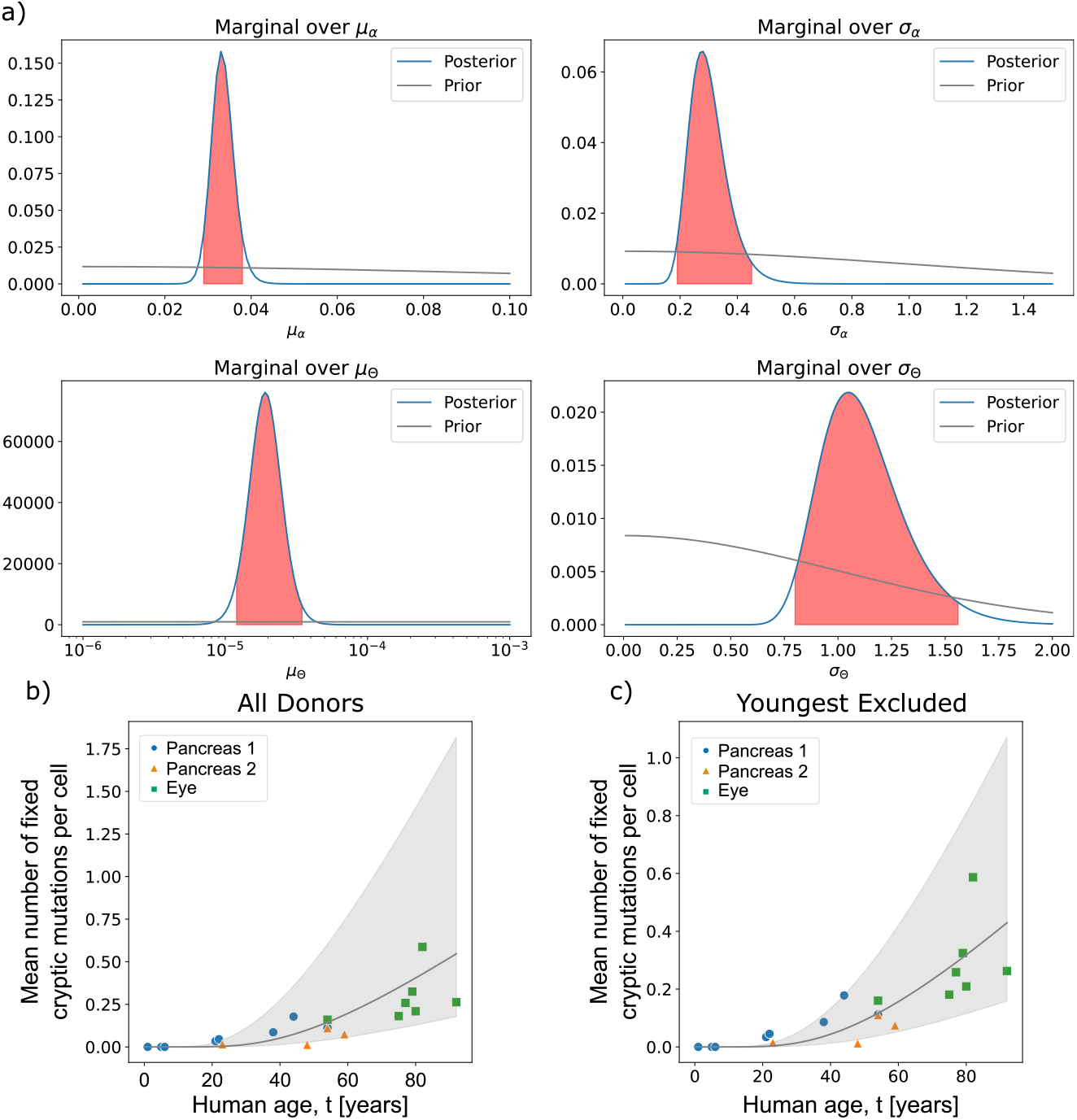
By excluding the youngest donor in the dataset, we see that posterior inferences tighten. a) The posteriors on the hyperparameters when the youngest donor is excluded from the dataset. b-c) Shown are the fits to the expected number of cryptic homoplasmic mutations. Marked in grey are the 95 % confidence intervals on the median of this fit. We see that by excluding the youngest donor the confidence interval shrinks substantially. We note that these are not fits done only to the homoplasmic mutations, but to the full set of mutations across the cSFS.

### S2.5 Fitting to mouse data

Because of the difference in proliferation rate between pancreas and liver tissue, we fitted the two mouse tissue types separately. For both mouse tissues we picked hyperprior distributions of *μ*_*α*_ *∼*Half-Normal(1.2), *μ*_Θ_ *∼*Half-Normal(0.01), *σ*_*α*_ *∼*Half-Normal(1), *σ*_Θ_ *∼*Half-Normal(1). We found that both the population mitochondrial ageing rate, *μ*_*α*_, and the population coalescent mutation rate, *μ*_Θ_ of both tissues to be much higher than that of the human data (see table S4). Both parameters depend on multiple biological parameters such as copy number, turnover rate and mutation rate, and so while we see the net effect of any differences between humans and mice results in a faster mitochondrial ageing rate and a higher coalescent mutation rate, we cannot disambiguate what is causing this difference. The posterior inferences for the hyperparameters of mouse data are particularly broad due to the lack of sufficient aged data (see Fig. S10), however the difference between the mouse and human parameters are still evident.

### S2.6 Comparing diabetic to healthy tissue

To demonstrate the applicability of our model to comparative studies, we analysed single cell RNA seq data taken from both healthy and diabetic pancreas tissue (*S17*). We fit the model separately on 1977 cells from 8 healthy donors and 1732 cells from 6 diabetic donors. First we found that, for the healthy donors, we credibly recapitulated the parameter values that we found in table S4. We further found there was evidence for a difference in the mitochondrial ageing rate, with a 93 % probability that *μ*_*α*_ was lower for the diabetic donors (Fig. S11b). To increase the power of the study, we merged the healthy donors from (*S17*) with one other healthy human pancreas dataset already analysed (*S5*) and found that the trend was preserved, with an improved 99 % probability that the mitochondrial ageing rate of the pancreas is slower for patients with diabetes compared to controls (Fig. S11c).

**Supplementary Figure S10:**
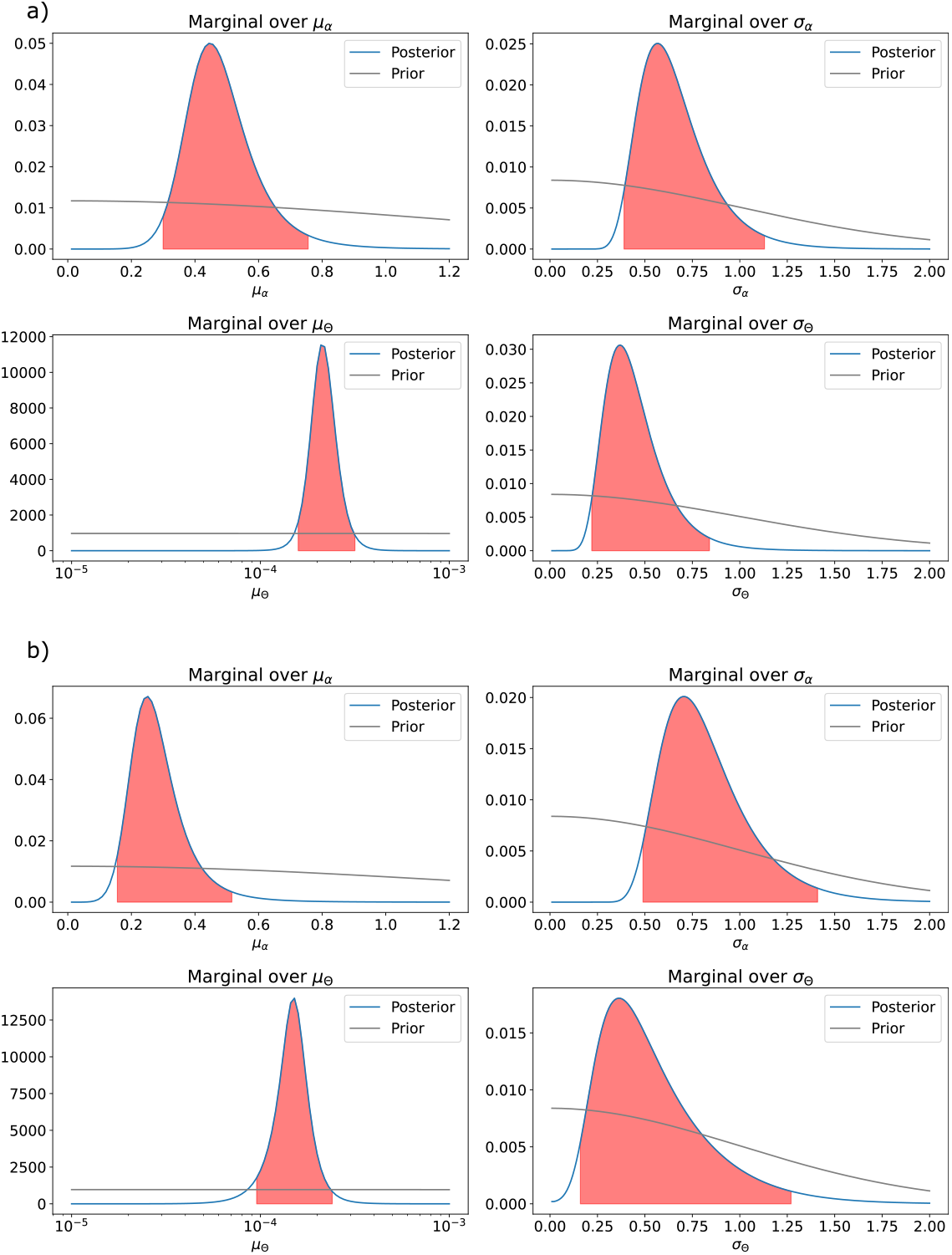
The posteriors on the hyperparameters for both mouse tissues. We see that for both tissues the posteriors are wider than those inferred for the human datasets, due to a reduced amount of single cell data from these tissues. a) The marginal posterior distributions for the mouse liver. b) The marginal posterior distributions for the mouse pancreas.

**Supplementary Figure S11:**
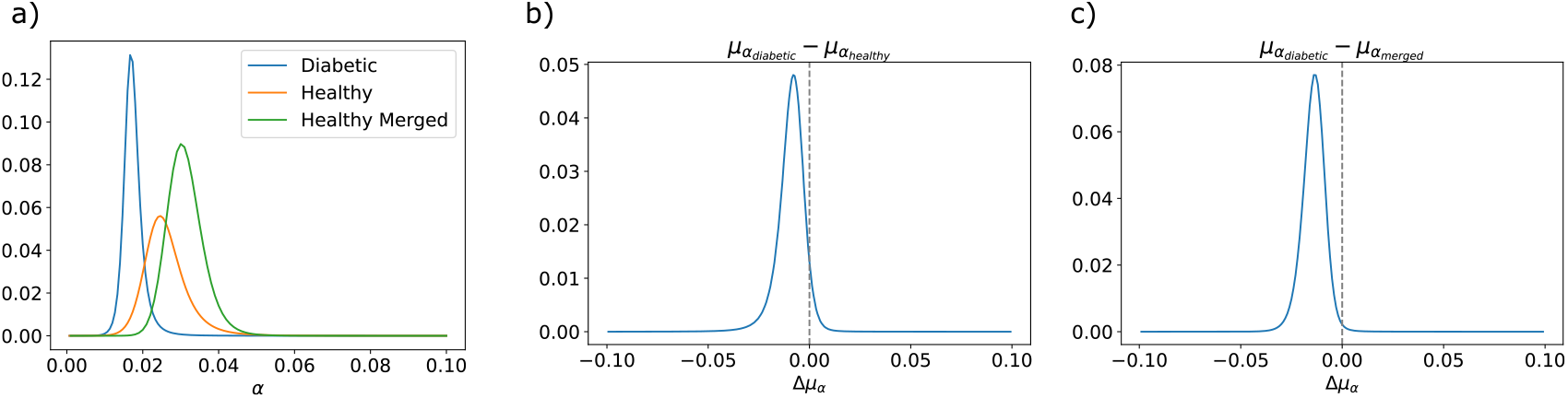
Diabetic pancreas tissue has a decreased mitochondrial ageing rate compared to healthy tissue. a) The marginal posterior distributions of the mitochondrial ageing rate, *μ*_*α*_. Shown are the posteriors on the diabetic donors (*S17*), the healthy donors (*S17*), and the healthy donors merged with data taken from healthy human pancreas from (*S5*). b) The distribution of the difference in *μ*_*α*_ between the diabetic and healthy tissue from (*S17*). 93 % of the probability mass lies below zero. c) When the healthy data (*S17*) was merged with another healthy pancreas dataset (*S5*), we found an increased effect on the value of *μ*_*α*_. For this merged difference, 99 % of the probability mass lies below zero, raising the hypothesis of a difference in ageing rate between healthy and diabetic pancreas.

## S3 Supplementary Discussion: Fisher’s method for DEG identification

### S3.1 Fisher’s method for donors

In the maintext, we identified differentially expressed genes (DEGs) aggregated for all cells. This has the advantage that even modest shifts in gene expression are identifiable. It comes with the disadvantage, however, that confounding factors might create stronger shifts than actually present. While the consistency of our significant GO terms across different organs, species, sequencing techniques, and ages (see Fig. 3 in the main text) suggests that the aggregate approach we use in the maintext is appropriate, we nonetheless, here, present a complementary approach to eliminate the possibility that the DEGs that we identified in the main manuscript are exclusively age-dependent genes. For this, we calculate DEGs for each donor separately and then aggregate the *p*-values of differential expression in a ‘meta’-analysis. Specifically, for a gene with *p*_*d*_ indicating the *p*-value of differential expression for donor *d*, we aggregate these *p*-values with Fisher’s combined probability test to a test statistic

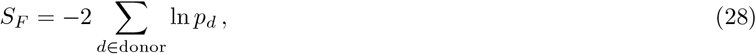

where *S*_*F*_ follows a *χ*^2^ distribution, which we may lookup to obtain a *p*-value.

We use this approach for the Enge (human pancreas data) and the two mice data sets (liver and pancreas). This means that *we only make comparisons between cells which have the same age* (in the narrow sense of all the cells coming from the same individual). For all three datasets we obtain significant genes that are differentially expressed (after multiple-testing correction). We then perform a GO-term enrichment analysis for the obtained DEGs and demonstrate that we obtain similar GO-terms (see Fig. S12).

Finding similar GO terms enriched indicates that the identified biological perturbations are not predominantly driven by a difference between donors. Rather, it leaves open the possibility that mitochondrial mutation load could induce a genetic perturbation that is identifiable at the resolution of each donor.

### S3.2 Fisher’s method for cell types

Similarly to the ‘meta’-analysis in Subsection S3.1 that only compares cells of the same donor, we can also perform a analysis that only compares cells of the same cell type. Specifically, for a gene with *p*_*c*_ indicating the *p*-value of differential expression for cell type *c*, we aggregate these *p*-values with Fisher’s combined probability test to a test statistic

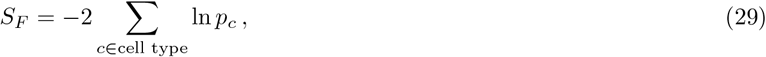

where *S*_*F*_ follows a *χ*^2^ distribution, which we may lookup to obtain a *p*-value.

Here, we analyse the Human pancreas data (Enge), for which the authors provide annotations that associate each cell with one of seven cell types (‘acinar’, ‘alpha’, ‘beta’, ‘delta’, ‘ductal’, ‘mesenchymal’, and ‘unsure’). Using this method, we find 628 differentialy expressed genes after multiple-testing correction. We then perform a GO-term enrichment analysis for the obtained DEGs and demonstrate that we obtain similar GO-terms to the maintext analysis (see Fig. S12). This indicates that the mitochondrial mutation load could induce a genetic perturbation that is identifiable at the resolution of each cell type. We, note, of course that this simple approach has the substantial caveat that cell-type assignments are themselves extracted from the full gene-expression data and do not constitute independent experiments: for this reason, combining across cell-types, our results combining p-values should be understood as indicative.

**Supplementary Figure S12:**
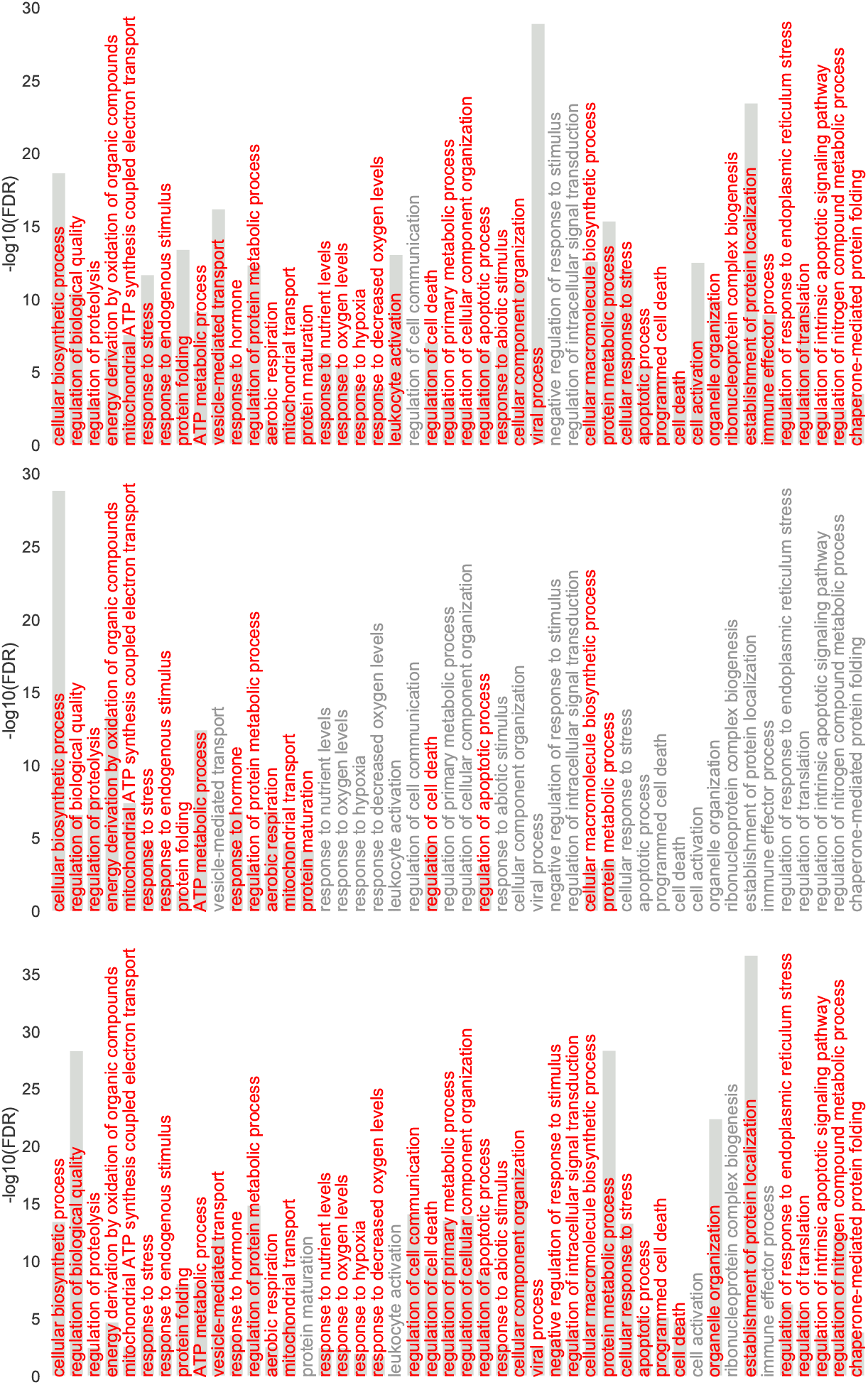
Transcriptional change correlated to the presence of cryptic mtDNA mutations is also present when making comparisons between cells of the same donor. Human pancreas (Enge, top panel) data, the mice liver (middle panel), and the mice pancreas (bottom panel). We show the highlighted GO terms from the main manuscript and highlight them in red if significant with the Fisher’s method.

**Supplementary Figure S13:**
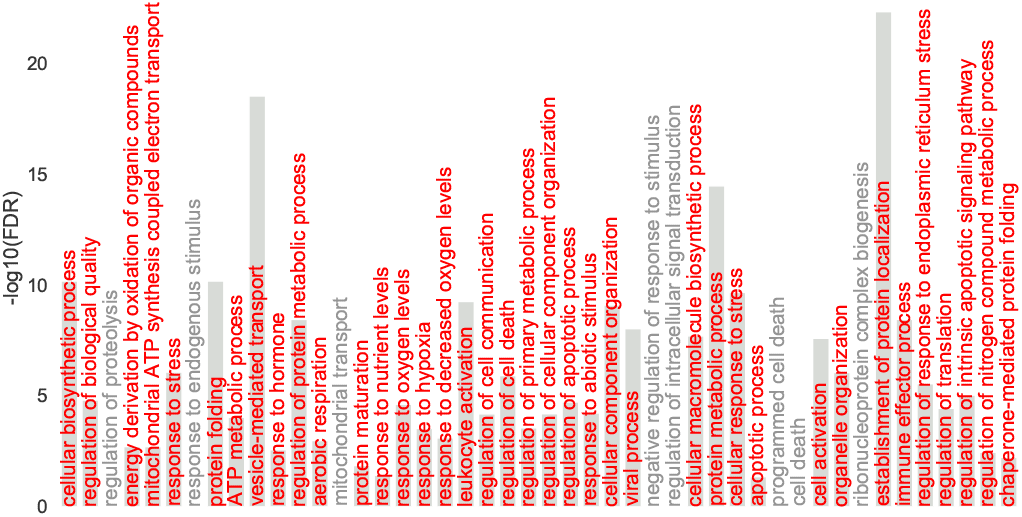
Transcriptional change correlated to the presence of cryptic mtDNA mutations is also present when making comparisons between cells of the same celltype for the Human pancreas data (Enge). We show the highlighted GO terms from the main manuscript and highlight them in red if significant with the Fisher’s method.

## S4 Supplementary Discussion: The role of the number of mitochondrial reads

Cells vary in their ratio Γ of the number of mitochondrial reads to the total number of reads, which could be a sign of varying mtDNA copying numbers. It is known that external factors, such as, oxygen tension may modulate the amount of mtDNA, as well as, heteroplasmy e.g. (*S18*). It is also long known that mtDNA mutations can alter copy-number (*S19*). The interplay between copy-number and mtDNA mutation is thus nuanced. For the human pancreas data (Enge *et al*.), we find that cells with *μ >* 0 (i.e., cells with a cryptic mutation above 10 %) tend to have a slightly higher percentage of mitochondrial reads than cells without such a mutation (⟨Γ⟩_*μ*=0_ *≈* 11.07 % and ⟨Γ⟩_*μ>*0_ *≈* 12.63 %). Therefore, it is a hypothesis that the differences in gene expression that we identify is exclusively driven by the variability of the ratio Γ between these two groups.

To test this hypothesis, we use a sampling procedure that constructs a new data set with ⟨Γ⟩_*μ*=0,sampled_ ≈⟩Γ⟩_*μ>*0_ *≈* 12.63 % (i.e. both sets of cells are controlled to have the same average number of reads) but otherwise the same characteristics (i.e., the same number of cells with and without cryptic mutations). To achieve this, we keep all mutated cells but sample with replacement from the cells without mutations such that ⟨Γ⟩_*μ*=0,sampled_ ≈⟨Γ⟩_*μ>*0_. Specifically, we construct a histogram with *n*_bin_ = 8 bins, depending on their Γ and then use an importance sampling procedure where the sampling weight for each bin is given by the ratio of the number of cells with *μ >* 0 to the number of cells with *μ* = 0 in this bin. This procedure yields a random sample with ⟨Γ⟩_*μ*=0,sampled_ *≈* 12.61. Repeating this procedure or increasing the number *n*_bin_ of bins yields similar results.

With this sampled data, we obtain a list of 1836 DEGs and the GO enrichment shows similar biological processes enriched as without the importance sampling procedure (see Fig. S14). This rejects the hypothesis the identification of DEGs is exclusively driven by a difference in the ratio Γ of the number of mitochondrial reads to the total number of reads.

**Supplementary Figure S14:**
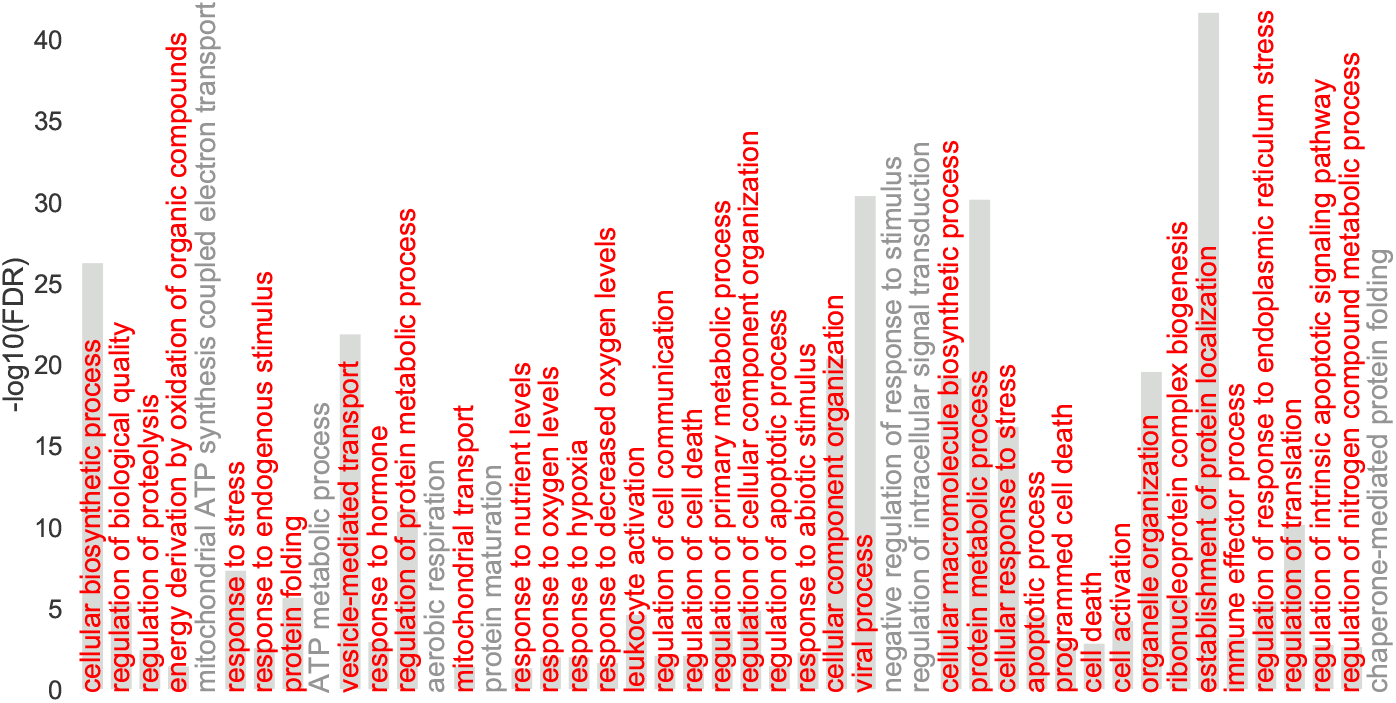
Transcriptional change from presence of cryptic mtDNA mutations is also present after correcting for the ratio Γ of the number of mitochondrial reads to the total number of reads (human pancreas data, Enge). We show the highlighted GO terms from the main manuscript and highlight them in red if significant after the correction.

## S5 Parkinson’s disease and Alzheimer’s disease single-nucleus RNA-seq data

In the main manuscript, we link cryptic mitochondrial load to gene expression in neuronal cells and in particular with genes linked to neurodegeneration. We showed this for a single-nucleus RNA-seq (snRNA-seq) data consisting of donors with Parkinson’s disease (PD) and age-matched controls. Here, we investigate a second data set consisting of patients with Alzheimer’s disease (AD) donors and age-matched controls (*S20*). Given the small number of mitochondrial reads in snRNA-seq data (see Fig. S21), we call heteroplasmies at 10 reads and mark cells with a cryptic mutation which is not synonymous above 95 % as ‘mutated’. Our aim is to identify whether the expression of genes is linked to the presence of cryptic mutations.

This data set is considerably smaller (3,759 cells after quality control, in comparison to 27,539 in the PD data) and therefore, we investigate only the top 6000 genes, to reduce the influence of the Benjamini–Hochberg procedure for multiple testing correction.

We find only MALAT1 to be differentially expressed after the multiple-testing correction. The functional role of the lncRNA MALAT1 was first identified in cancer (*S21*) but has since then also been associated with neurodegenerative diseases (*S22*). More generally, the identification of differentially expressed lncRNAs in the different data sets indicates that their involvement in response to cryptic mutations might be similar to their dysregulation in human disorders (*S23*).

Heat shock 70 kDa protein 1 (HSPA1A) and Glial fibrillary acidic protein (GFAP) are differentially expressed but not significant after the multiple testing correction. For both, however, there is external evidence that suggests a neurodegenerative function. The expression of GFAP in astrocytes is linked with old age and the onset of AD (*S24*). Heat shock proteins interfere with apoptosis and these homeostatic functions are especially important in proteinopathic neurodegenerative diseases (*S25*).

Overall, our results demonstrate that one can call high-heteroplasmy mutations from nuclei instead of whole cells. Despite the low coverage of mtDNA reads in snRNA-seq, we are able to identify cells with a high mitochondrial mutation load. We find evidence of a link between gene expression and cryptic mutations. In particular, the identified DEGs indicate that there is a change in inflammatory processes and lncRNAs which may be linked to neurodegeneration and senescence.

**Supplementary Figure S15:**
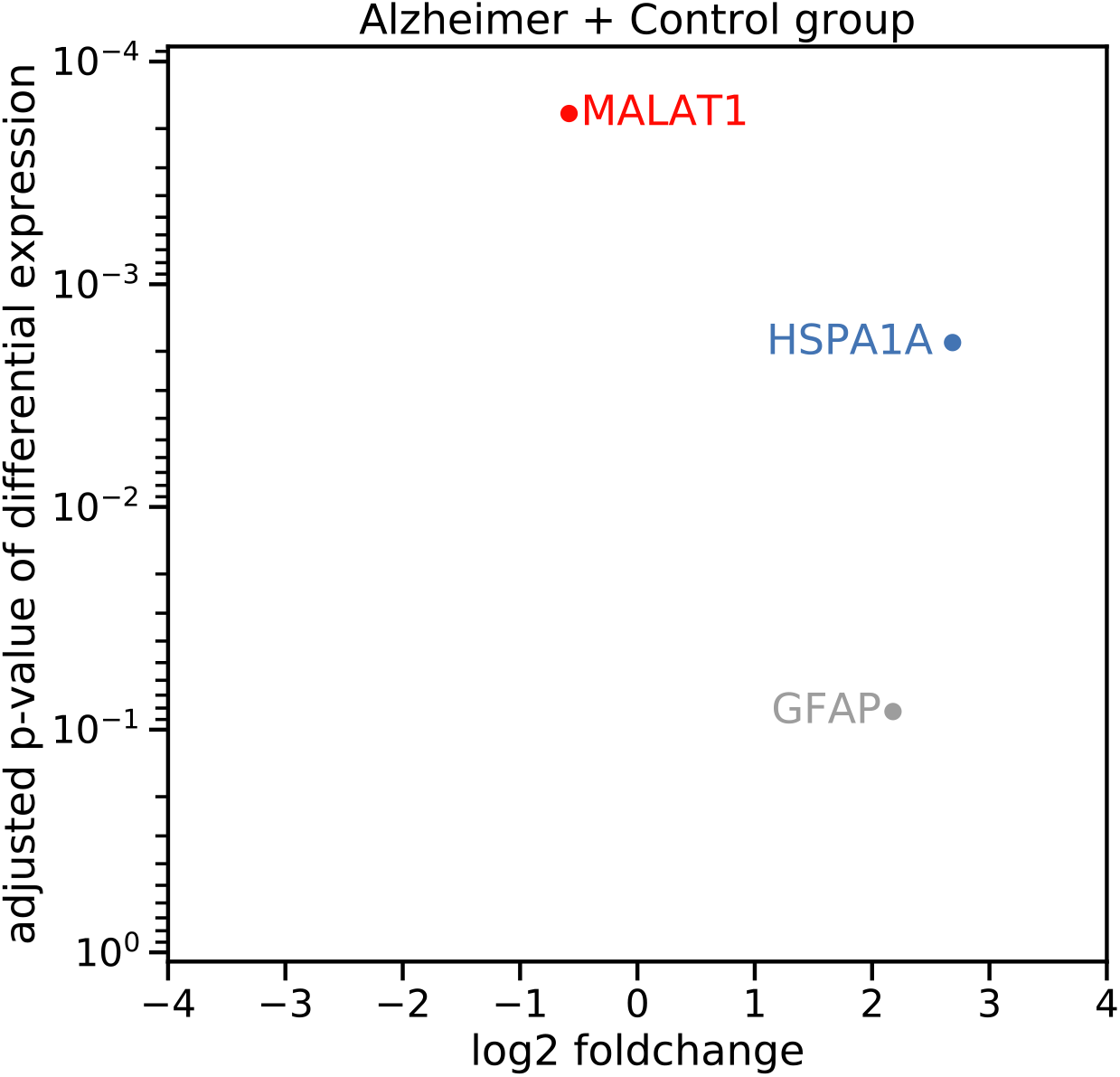
Differentially expressed genes in the Alzheimer’s disease (AD) data. We aggregate AD and Control group cells to identify genes that are perturbed in both groups. For greater statistical power, we only investigate the top 6000 most variable genes, which reduces the influence of the Benjamini–Hochberg procedure for multiple testing correction.

## S6 Supplementary Discussion: Variant Calling from RNA

### S6.1 Empirical corroboration

Throughout the manuscript, as well as using scATAC-seq, we analyse mitochondrial variants identified using scRNA-seq data, following similar efforts using scRNA-seq to do lineage tracing with mtDNA variants (e.g., (*S26* –*S28*)). We perform our own analysis of how accurately scRNA data reflects variants on the underlying DNA using paired scATAC and scRNA data from the 384 cells taken cultured from 4 different cell lines (*S29*). Both scATAC-seq and scRNA-seq libraries were aligned using STAR (*S30*) and then we identified variants using our custom variant calling script. The libraries were generated with a paired sequencing protocol which involved 7 more rounds of PCR that the standard smartseq2 protocol, and these rounds were held at higher temperatures for longer, which will results in more thermal errors (*S31*) spreading to higher heteroplasmies than the other datasets considered in the main text. In addition to our usual quality control filters, we only take forward positions for variant analysis if they pass were covered by at least 200 reads in both the scATAC-seq and scRNA-seq libraries. After all quality control was applied there were 314 cells taken forward for analysis. Despite the quality of the dataset, we found that 92.2 % of variants with heteroplasmy above 10 % identified using scATAC-seq were also found using scRNA-seq, and there was a strong correlation of heteroplasmy between mutations identified by both techniques (Pearson test *r*^2^ = 0.76).

In accord with Ludwig *et al*. (*S26*) we also find many RNA specific mutations which likely come from RNA editing events or sequencing errors. 76.9 % of these mutations occur in more than one cell, likely reflecting common RNA mutations and so will be filtered out when we look at cryptic mutations. In this study of data from (*S29*) we could (as we do in the maintext) improve the correlation between heteroplasmy called from RNA and DNA by using more stringent depth or heteroplasmy thresholds, but these trade off against the number of mutations detected reducing statistical power.

### S6.2 Robustness to variant calling errors

Our study calls variants from scRNA-seq and snATAC-seq and these are termed cryptic mutations: its conclusions are robust to errors in calling any specific variant. We ensure robustness through comparative and aggregate analysis and careful selection of quality-control thresholds to eliminate noise sources. Our results are consistent with theory and are consistent across multiple tissues and species.

#### Aggregation and comparison

Our principal conclusions depend on comparisons of the cryptic site frequency spectrum (cSFS) and on creating populations of cells which are enriched in cryptic mutations. The process of calling variants, and assigning heteroplasmies, convolves a latent ‘true’ SFS distribution with an error kernel (existing mutations are assigned erroneous heteroplasmies through sampling effects and equivalents) and then adds a separate error distribution (non-existent mutations are erroneously identified). By making comparisons *between* cSFS distributions we control for common additive noise effects and our rational use of quality control (discussed below) and success at discriminating between ages of individuals suggests that the noise kernel has sufficiently bounded-variance. In a similar manner our differential expression analysis (comparing cells with and without cryptic mutations) does not require that we accurately identify cryptic mutations in every cell: only that we successfully create a set of cells which have an enriched rate of cryptic mutations.

#### Quality control

As noted by others there exist recurring RNA-mutations which may or may not be functional (*S32*): we are able to eliminate this source of confounding from our study by restricting our study to cryptic mutations: those found in only one cell in the sample. As noted in our methods section these mutations might effect our analysis of selection effects in non-cryptic mutations (Fig. 2b) but we attempt to mitigate this by eliminating mutations that are found in 3 or more individuals. We also seek to eliminate noise from PCR and sequencing errors resulting in erroneous mutation calling, while maintaining the signal we wish to see. From our theory in S1 we find that the ageing effect of interest causes dynamics in the mid to high-heteroplasmy ranges of the SFS, and so we apply conservative filters to both the depth required for variant calling, and the heteroplasmy of variants we consider to eliminate noise which could obscure the signal of interest. To this end we consider sites with *>* 200 reads aligning and take forward variants with heteroplasmy *>* 10 %. In choosing these thresholds we not only focus on the physiologically relevant high-heteroplasmy range in the SFS, we also work to exclude PCR errors with the following argument. From the UMI data we have available, we find that the average number of reads per UMI is 7, and so, if we set the threshold for read depth as *N*, it represents on average ^*N*^*/*_7_ true initial RNA molecules. Assuming a lower bound of PCR efficiency of 90 %, we calculate the maximum heteroplasmy a mutation occurring in the first round of PCR (and hence initial heteroplasmy 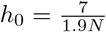) will reach after 37 rounds of PCR to be 11.5 % when our read threshold is set at *N* = 200. Any positions with a higher read depth, or any mutation that occurs after the first round of PCR will have a lower maximum possible heteroplasmy, and so, by combining a heteroplasmy threshold of 10 % with minimum read-depth 200, we exclude the vast majority of PCR errors. We do not consider bases with a sequencing error probability of *>* 0.001 and so by choosing our thresholds to be this conservative there is a *<* 10^*−*15^ probability of an erroneous heteroplasmy call from sequencing error.

#### Corroboration

We give a recapitulation of our main results using snATAC-seq in the next-but-one section. Our choices are also corroborated by the close accord between the our mathematical models of the extended site frequency spectrum (SFS) for a population of mtDNA and the cryptic site frequency spectrum (cSFS) we obtain. We find that young individuals have very few high frequency mutations in their cSFS (e.g. Fig. 1g and consistent with theory Fig. 1d) and that the cSFS also evolves with individual age in accord with theory (e.g. Fig 1h, Supplementary Material S1). Beyond finding results from the cSFS consistent with theory, our consistent results across species add further support. While our results on the dynamics and effects of cryptic mutations would be equally interesting if they were driven by mutations that only occurred in the mtRNA (and not in the mtDNA) it is, however, challenging to provide a theoretical account for the dynamics of the cSFS if it is not being driven by mutations in the mtDNA.

### S6.3 The effect of UMI collapsing on heteroplasmy calls

**Supplementary Figure S16:**
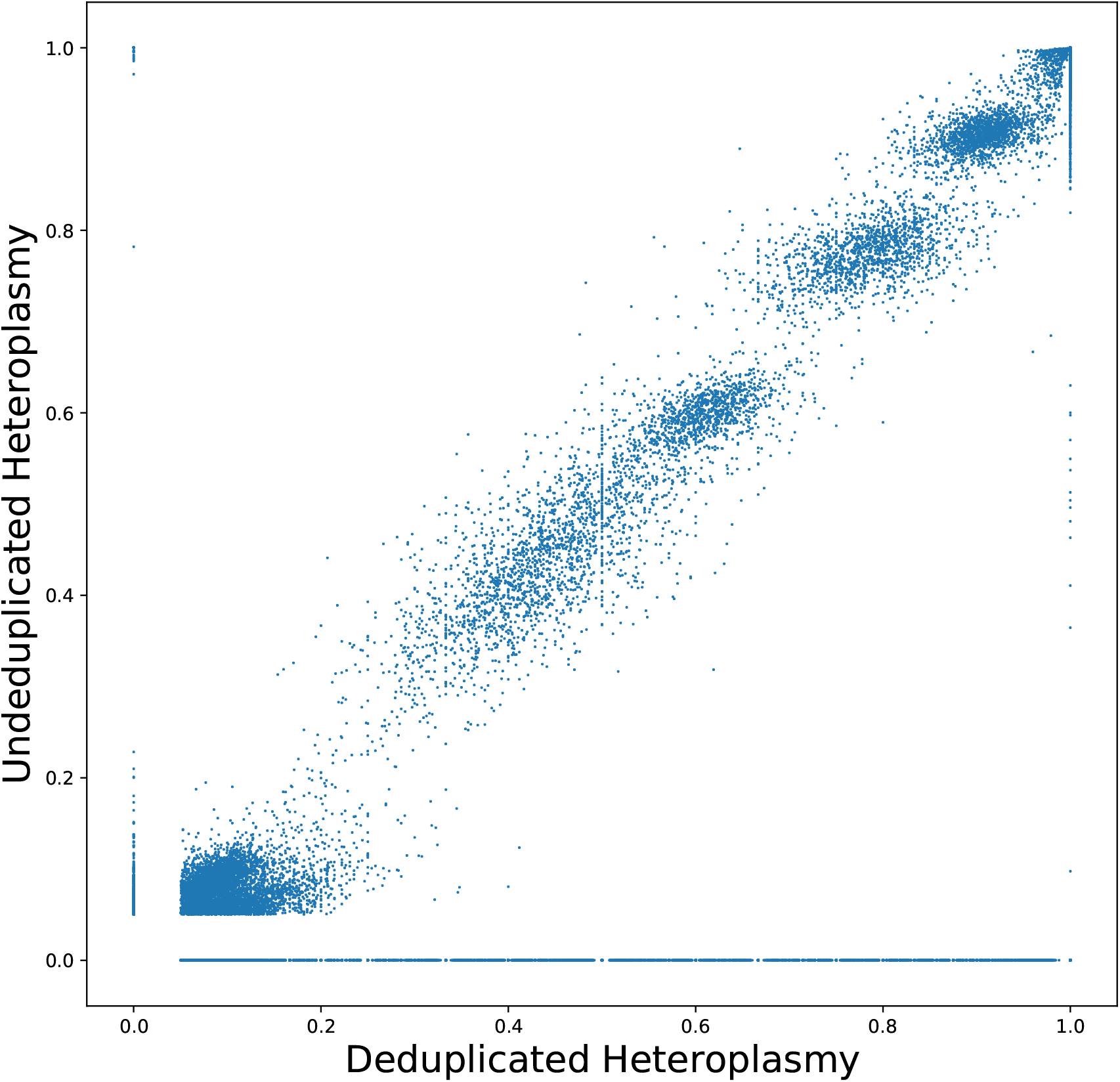
Heteroplasmy calling is in broad agreement both with and without UMI collapsing. Here we called heteroplasmic variants using both raw reads and also by picking a representative read for each UMI in each cell with umi tools dedup function (*S33*). We called variants for the undeduplicated data at 200 read depth and deduplicated at 5 UMIs and estimated heteroplasmies for the Voigt data. This was done at a 5% heteroplasmy threshold. It can be seen that many variants are observed at the line y = 0 in the deduplicated data. These variants are likely attributable to the sequencing error caused by using so few UMIs and choosing a random representative. Therefore, deduplicated heteroplasmy calls at y=0 were excluded from regression analysis. We find excellent agreement between the heteroplasmies upon deduplicating and not deduplicating UMIs with *R*^2^ = 0.991

#### S6.4 Mutations detected in snATAC data recapitulate results found from scRNA derived mutations

To further corroborate our findings from scRNA derived mutations, we reproduce key results of the main text using a 10x single-nucleus ATAC dataset (*S9*) taken from various brain tissues of 6 cognitively healthy individuals with no neuropathological hallmarks of Alzheimer’s. As the data was taken from a single nucleus dataset, it had much lower coverage of the mtDNA that our data used in the main text, and so we relax our depth threshold for variant calling to 10 reads, though we find that even with the extra noise this will introduce we replicate results regarding the evolution of the cSFS and selection of non-cryptic mutants from the main manuscript (Fig. 1f-h, 2b) as well as finding further evidence for tissue specific selection of non-synonymous cryptic mutations at heteroplasmies below *h <* 10 % (Fig. 2d).

**Supplementary Figure S17:**
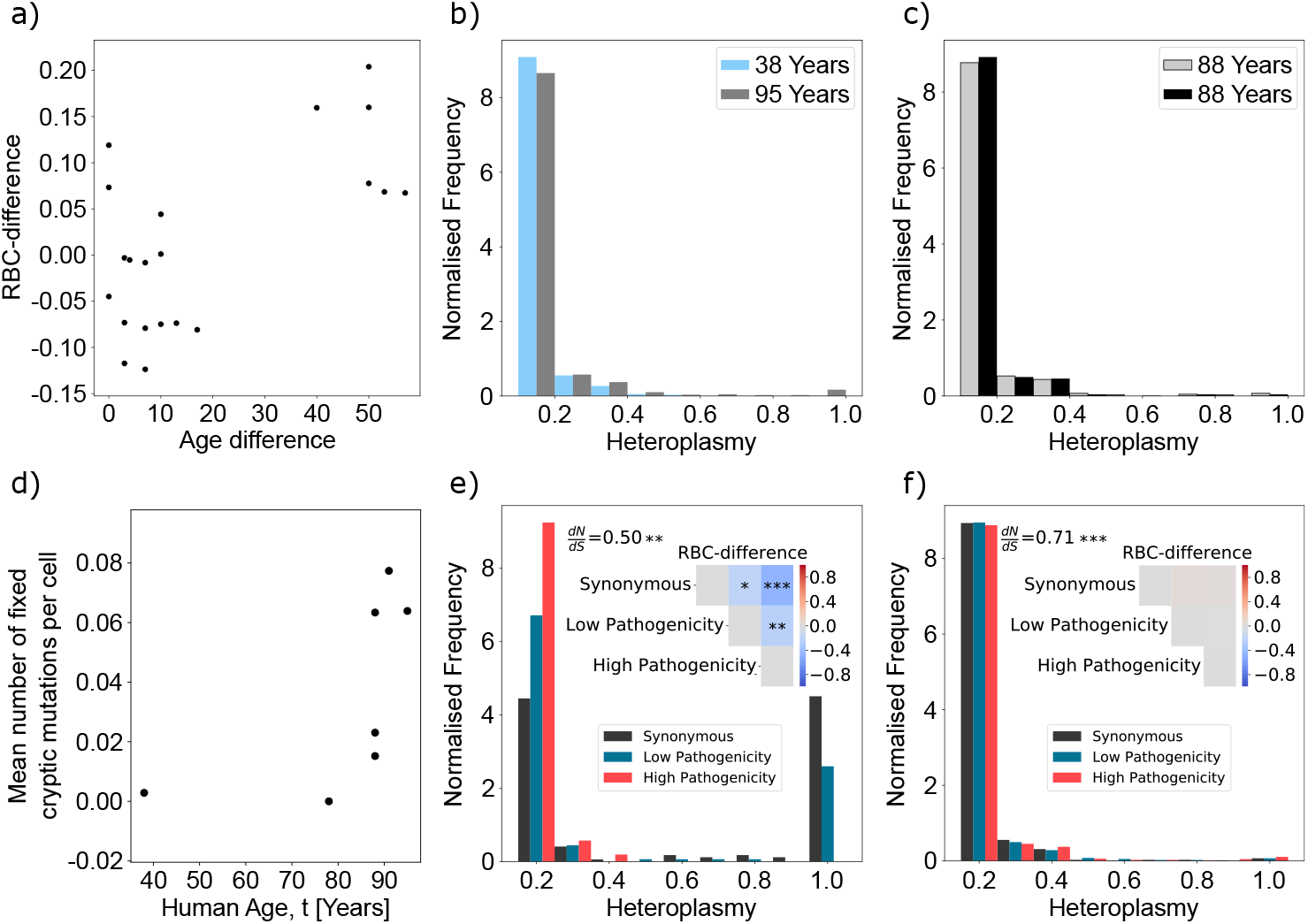
Results seen in the main manuscript are reproduced in a 10x single-nucleus ATAC dataset (*S9*) derived from human brain tissue from 6 donors. (a) The RBC-difference (see Methods 2.6) between donors of different ages significantly increases with age (Pearson correlation *r ≈* 0.64 and *p <* 0.01). (b-c) We present two pairs of cSFSs taken from donors of differing ages (b) and donors of the same age (c) to see that, as in the cSFSs shown in the main manuscript, older individuals have a cSFS shifted to higher heteroplasmies than younger individuals. (d) There is a significant increase of cryptic homoplasmic (heteroplasmy *>* 0.9) mutations with age (Spearman correlation *r ≈* 0.82 and *p <* 0.01). (e) We also look at selection effects first on all non-cryptic mutations and find a significant shift in the non-synonymous/synonymous ratio below 1 (Fisher’s exact test *p <* 10^*−*5^) as well as in the SFS of both high and low pathogenicity mutations when compared to synonymous mutations (*p <* 10^*−*10^, 0.01 respectively). Additionally we find a significant shift in the SFSs of low pathogenicity mutations compared to high pathogenicity mutations (*p <* 0.001), indicating that the degree of pathogenicity modulates the degree of selection affecting a mutation in the germline. (f) Cryptic mutations have a significant shift below 1 of their non-synonymous/synonymous ratio (Fisher’s exact test *p <* 10^*−*10^), but as seen in mouse data (Fig. 2d) there is no evidence of a shift in the cSFS hinting at selection occurring on mutations with a heteroplasmy *<* 10 %.

## S7 Supplementary Tables

### S7.1 Cell and mutation numbers for displayed normalised site frequency spectra

**Table S5:** For every displayed normalised site frequency spectrum we give the total number of cells that the mutations were found in as well as the total number of mutations which make up the SFS (for non cryptic SFSs we count each unique mutation once per individual and take the average heteroplasmy it is found at).

| Figure | SFS label | Cells | Mutations |
| --- | --- | --- | --- |
| Fig. 1c | N/A | 129 | 211 |
| Fig. 1g | 1 Month | 93 | 206 |
| Fig. 1g | 54 Years | 129 | 211 |
| Fig. 2a | Synonymous | 201 | 229 |
| Fig. 2a | Low Pathogenicity | 312 | 375 |
| Fig. 2a | High Pathogenicity | 290 | 337 |
| Fig. 2b | Synonymous | 2180 | 92 |
| Fig. 2b | Low Pathogenicity | 2166 | 48 |
| Fig. 2b | High Pathogenicity | 160 | 29 |
| Fig. 2c | Synonymous | 836 | 57 |
| Fig. 2c | Low Pathogenicity | 839 | 42 |
| Fig. 2c | High Pathogenicity | 478 | 19 |
| Fig. 2d | Synonymous | 585 | 709 |
| Fig. 2d | Non-synonymous | 1132 | 1797 |
| Fig. 4a | Young Ad Libitum | 50 | 50 |
| Fig. 4a | Old Caloric Restricted | 96 | 100 |
| Fig. 4a | Old Ad Libitum | 67 | 68 |
| Fig. 4b | Young Ad Libitum | 125 | 133 |
| Fig. 4b | Old Caloric Restricted | 145 | 154 |
| Fig. 4b | Old Ad Libitum | 58 | 58 |
| Fig. S10b | 38 Years | 471 | 3862 |
| Fig. S10b | 95 Years | 373 | 2985 |
| Fig. S10c | 88 Years (grey) | 395 | 2674 |
| Fig. S10c | 88 Years (black) | 125 | 706 |
| Fig. S10d | Synonymous | 1808 | 194 |
| Fig. S10d | Low Pathogenicity | 1493 | 180 |
| Fig. S10d | High Pathogenicity | 144 | 163 |
| Fig. S10e | Synonymous | 706 | 838 |
| Fig. S10e | Low Pathogenicity | 799 | 967 |
| Fig. S10e | High Pathogenicity | 724 | 849 |

### S7.2 LHON mutant haplogroups

**Table S6:** Haplogroups and accession numbers for bulk cell line data used in Fig. 4a-h

| Variant | Haplogroup | GeneBank Number |
| --- | --- | --- |
| 11778 | J1c1b | MH080306 |
| 3460 | J1c2e2 | MH080305 |
| Control1 | J1c8a | JN635299.2 |
| Control2 | J1c5 | JN635301.2 |
| Control3 | J1c1g | JN635302 |

## S8 Supplementary Figures

### S8.1 Differing sequencing techniques have varying coverage of the mitochondrial genome resulting in fewer possible variant locations

**Supplementary Figure S18:**
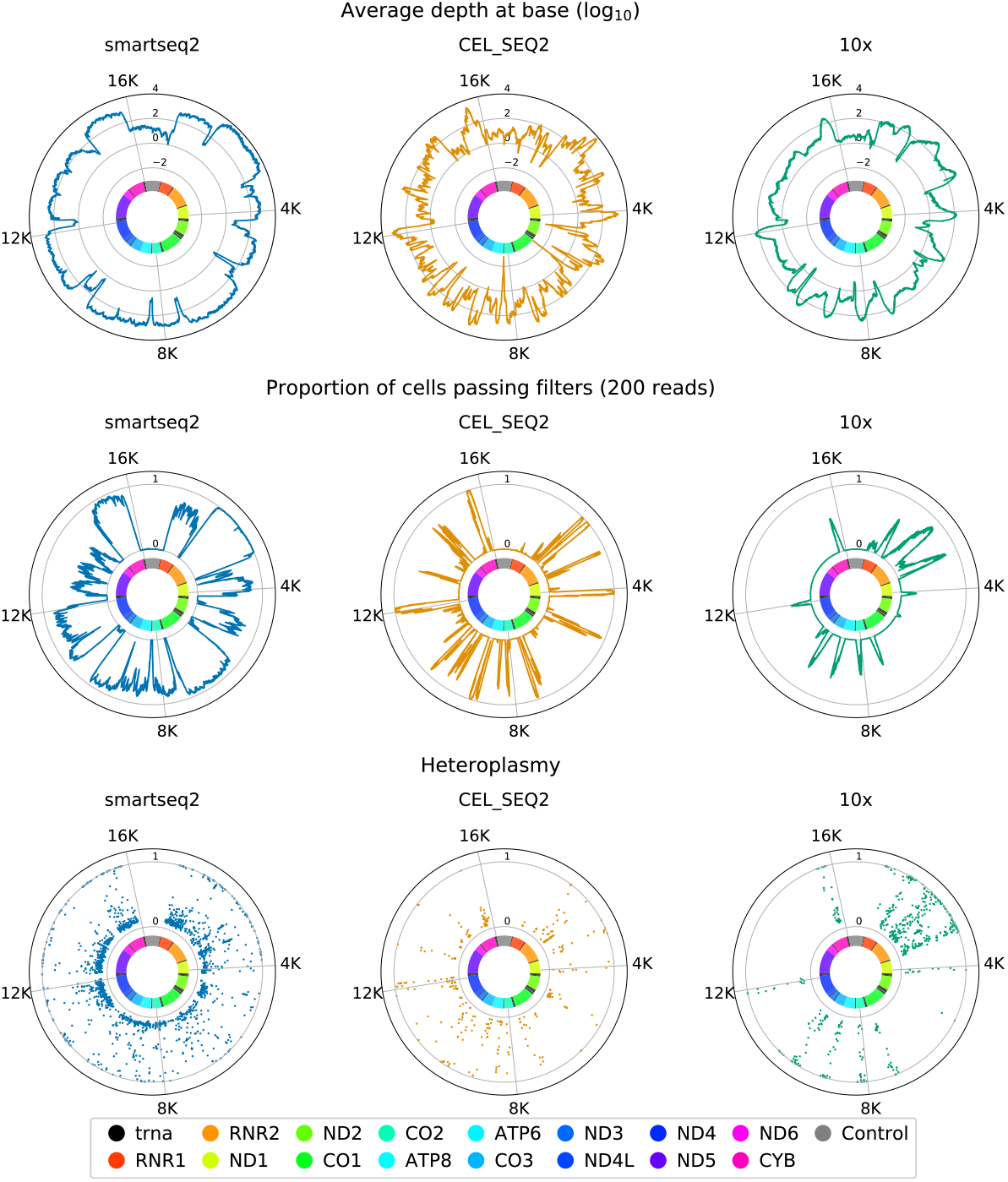
Throughout the paper we use data from three different scRNA sequencing technique, both full length and 3’. We show how the coverage of the mitochondrial genome differs for each technique, and how this affects the number of potential mutations we can identify. As expected full length sequencing techniques provide the most coverage across the mitochondrial genome, but for all techniques we observe that mutations are spread evenly across the sequenced regions

**Supplementary Figure S19:**
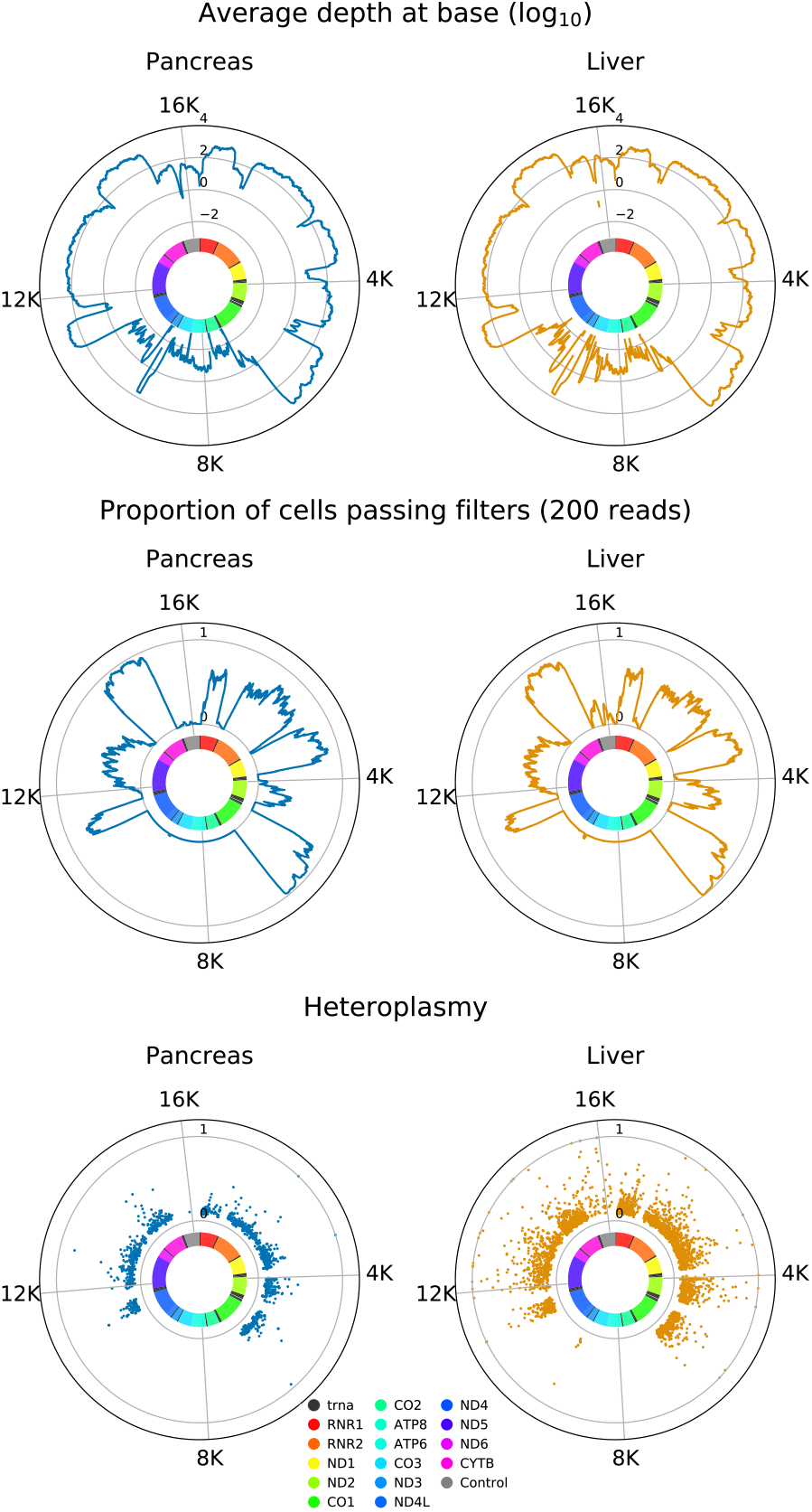
We show how the coverage of the Tabula Muris Senis dataset (*S16*) differs to that of human datasets - notably there is a drop in coverage between 6500-11000bp of the mtDNA of mice due to a large NUMT in the reference genome and our exclusion of multimapped reads

**Supplementary Figure S20:**
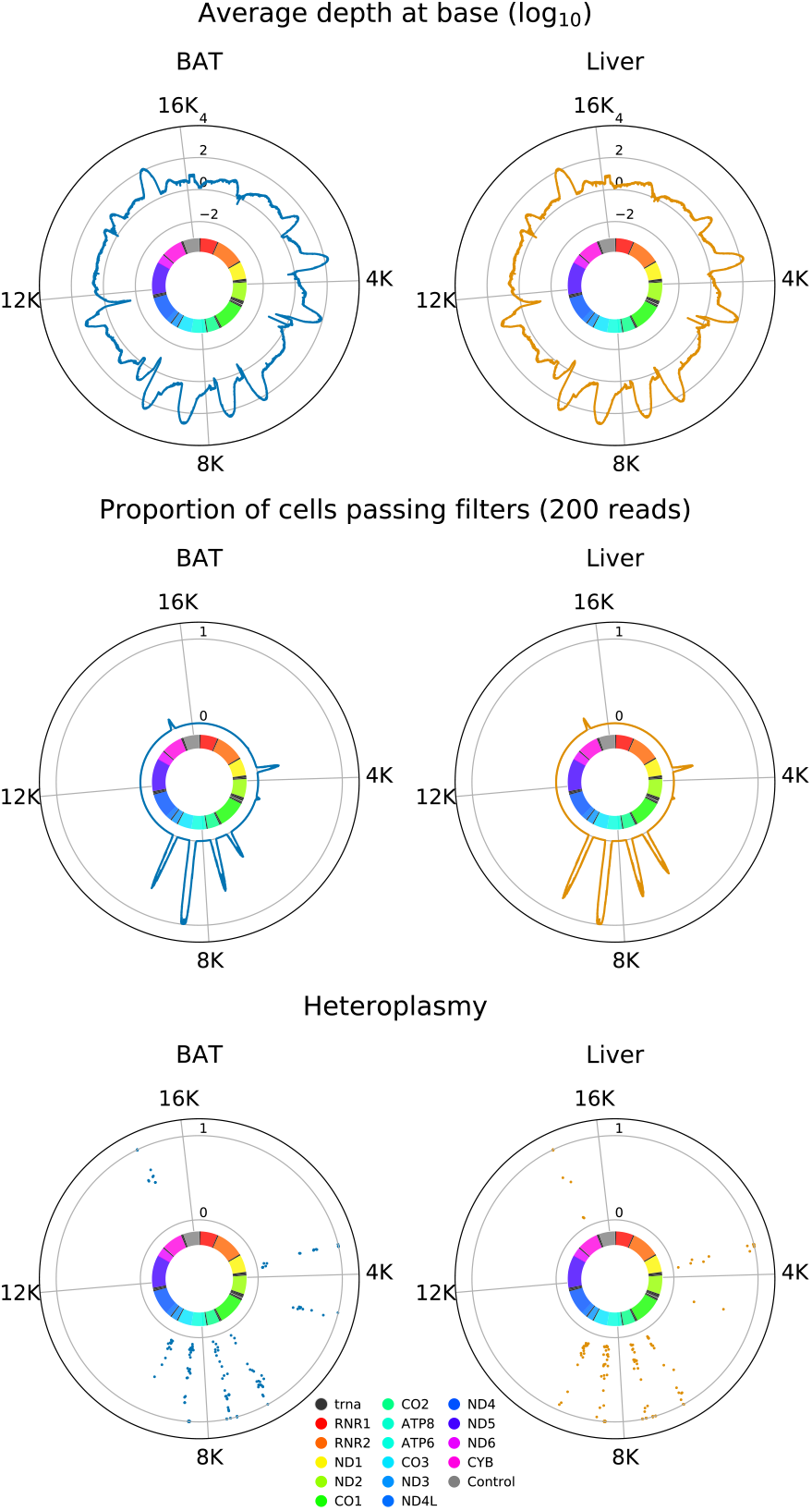
We also show the coverage of the rat caloric restriction dataset (*S34*) and find that the coverage of this dataset is an order of magnitude less than the equivalent 10x dataset in humans. In order to investigate caloric restriction, this dataset had variants called at a 5% heteroplasmy threshold, enabling greater quantities of variants to be detected in the dataset. Variant calling on both liver and brown adipose tissue at the 5% and 10% threshold had the mean heteroplasmy of mutations in young ad libitum and old calorically restricted mutations within 3% of each other, whilst the old ad libititum always had a mean heteroplasmy *>* 5% larger than both young ad libitum or old calorically restricted. Combining p-values from Mann-Whitney U tests between groups for both tissues for each difference separately using Fisher’s method gives the following p-values 0.381, 0.215, 0.131, for the parenthesised comparison pairs (young ad libitum, old calorie restricted), (young ad libitum, old ad libitum), (old calorie restricted, old ad libitum) respectively.

**Supplementary Figure S21:**
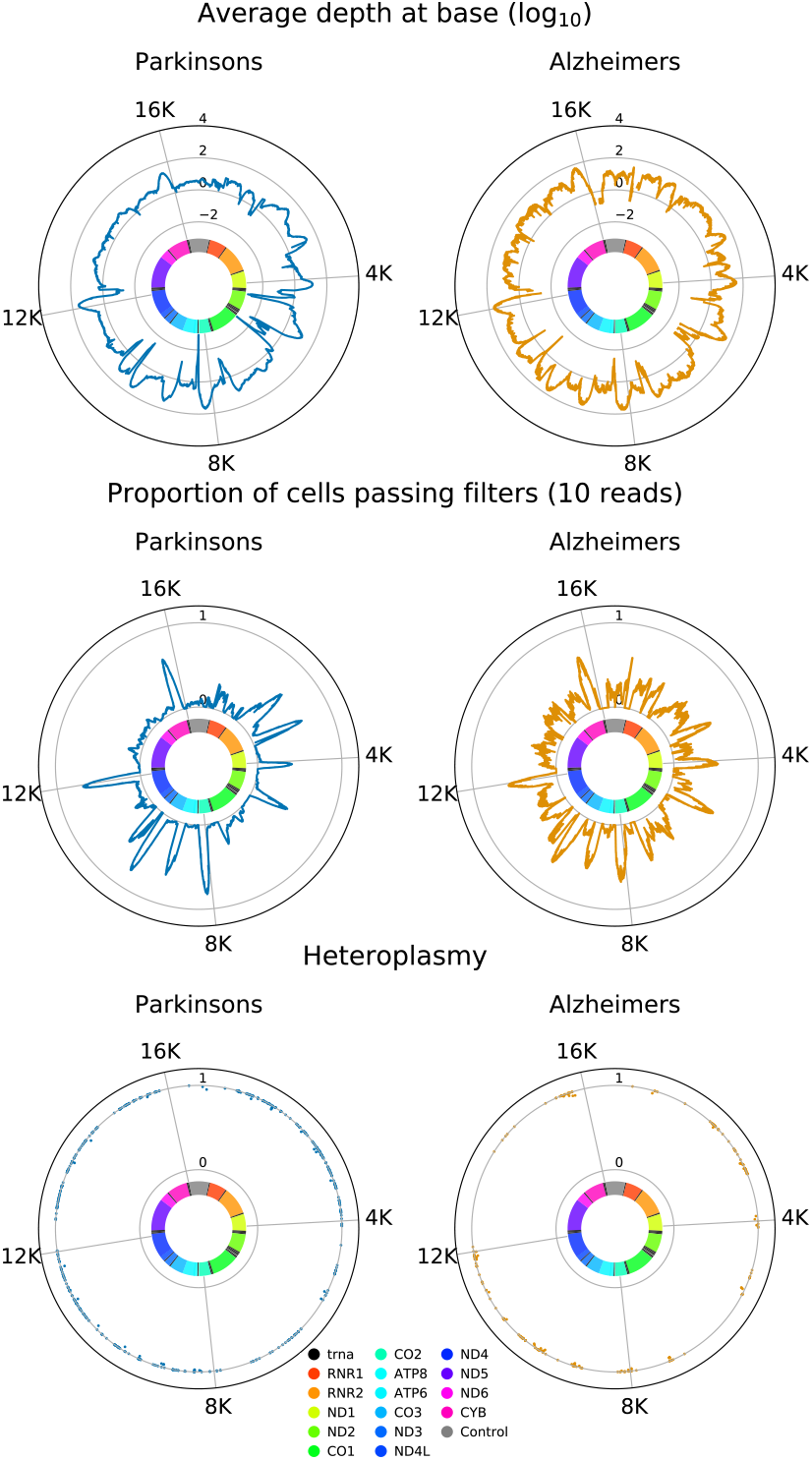
We show how the coverage from two single-nucleus datasets is consistently lower than that of single-cell experiments. To be able to use this data we relax our read depth threshold to 10 reads, and only call variants if their heteroplasmy *h >* 95 %

### S8.2 Increasing age difference evolves the cSFS further apart

**Supplementary Figure S22:**
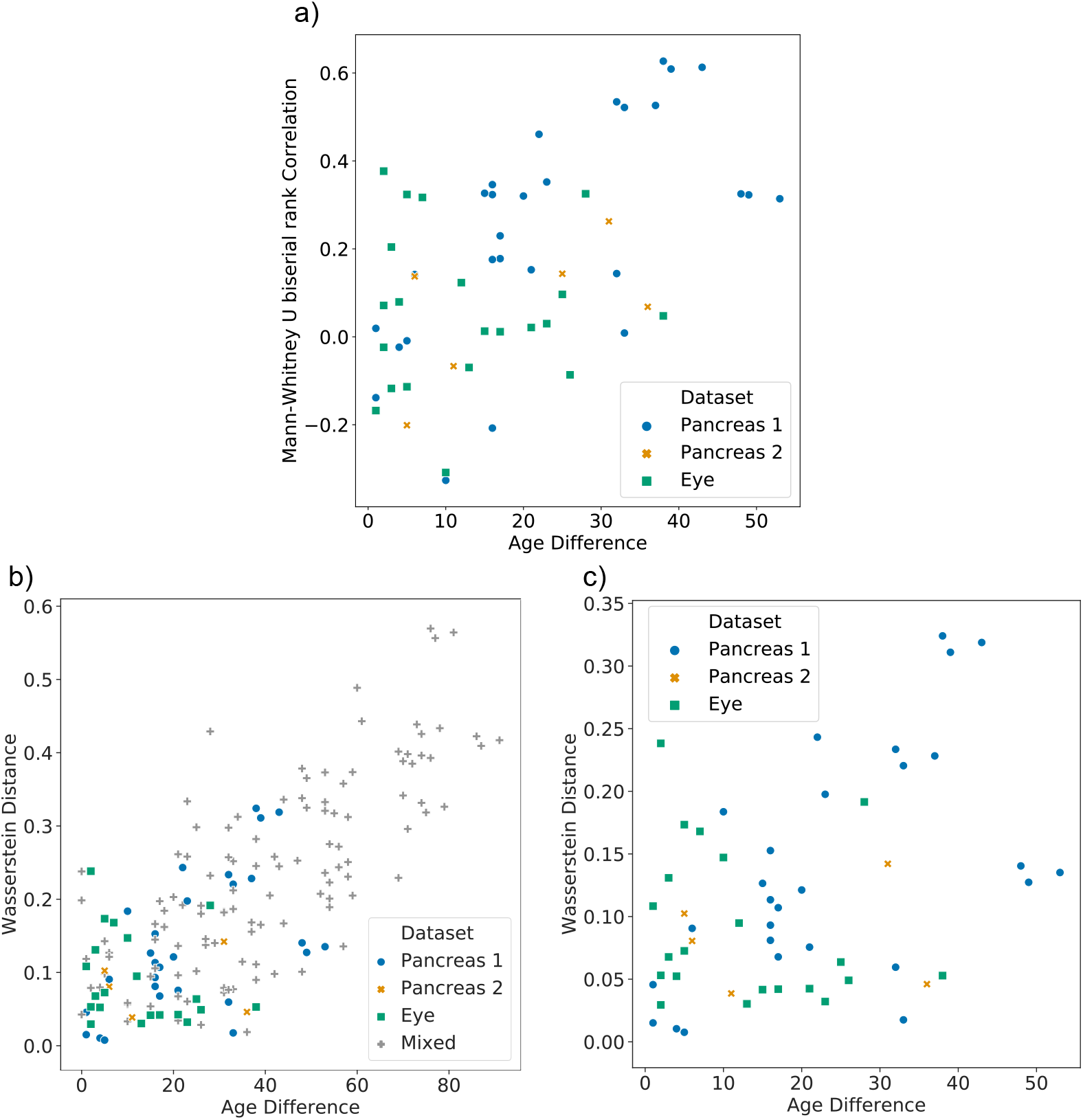
Using other metrics of distance between cSFSs we still find a significant correlation between distance between spectra and difference in age (a) When no comparisons between donors from different datasets is done, we still see a significant increase in the rank biserial correlation difference (RBC-difference, see methods 2.6) with difference in donor age (Spearman correlation *r ≈* 0.52 and *p <* 10^4^). (b) We can compare the cSFS of donors using the Wasserstein distance as an alternative measure of distance and find that the Wasserstein distance also significantly increases with age (Spearman correlation *r ≈* 0.70 and *p <* 10^*−*25^). (c) This significant increase also holds if we only consider comparisons of donors taken from the same datasets (Spearman correlation *r ≈* 0.33 and *p <* 0.05)

### S8.3 Restricting to a single cell type leaves results unchanged

**Supplementary Figure S23:**
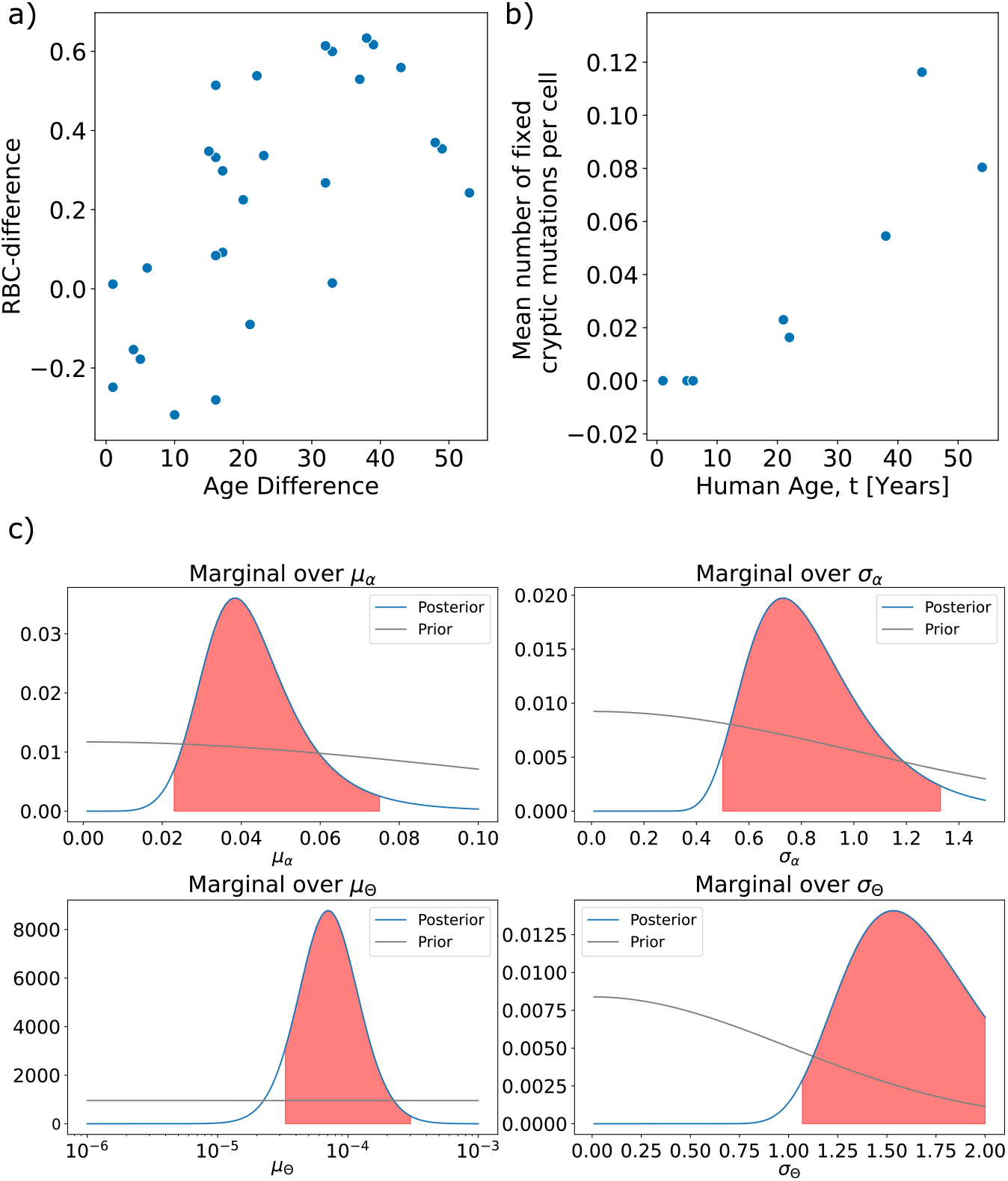
We restrict ourselves to only using the most abundant cell type of the Enge pancreas dataset (*S5*) (alpha cells) and find results are unchanged. a) The RBC-difference between cSFSs increases with the age difference between the donors (Spearman Correlation *r ≈* 0.64 and *p <* 10^*−*3^). b) The number of homoplasmies increase with age (Spearman Correlation *r ≈* 0.91 and *p <* 10^*−*2^). c) The inferred posteriors of the hyperparameters are wider due to the reduced amount of data, and the inferred population mutation rate is higher, driven by the youngest donors.

### S8.4 Mitochondrial ageing has multiple eras corresponding to mutations accumulating at different heteroplasmy levels

In the main text we show that the model predicts an early life equilibration of low heteroplasmy mutations, a mid life accumulation of mid-high heteroplasmy mutations and a late life accumulation of homoplasmic mitochondrial mutations. Shown in figure S24 are these potential “mitochondrial eras”, with all fits done using the MAP estimates from our Bayesian model (see Supplementary Discussion S2). The black line shows the Kullback–Leibler divergence of the out-of-equilibrium cSFS of heteroplasmies from its in equilibrium counterpart. Its fast decrease indicates that the majority of the cSFS, which is found at low heteroplasmies, equilibrates very quickly. The red line shows the relative probability of a heteroplasmic mutation being found at heteroplasmy *>* 60 % (excluding homoplasmy) when the out-of-equilibrium SFS is compared to the equilibrium SFS (60 % being an indicative heteroplasmy at which cellular dysfunction occurs). By 25 years old the relative probability is 0.25, increasing to 0.75 by 55 years. The blue line shows the mean number of homoplasmic mutations per cell, which undergoes non-linear dynamics up to around 55 years old, and then becomes a linear accumulation of mutations dependent only on the mutation and turnover rate of mtDNA. These eras align with early, mid and late life in humans.

**Supplementary Figure S24:**
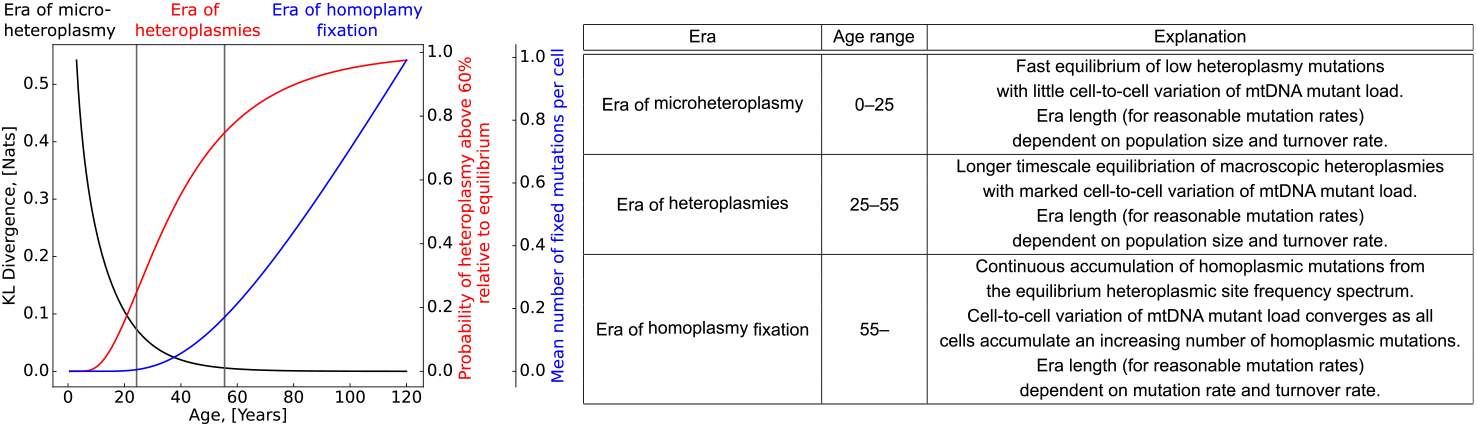
Three eras of mtDNA mutation accumulation. Table gives purely indicative ages. Notably it is only the very long term accumulation of homoplasmies, in late life, which resembles a linear accumulation of mutations. Turnover rate and cellular mtDNA population size define the equilibration phase (era of micro and macro heteroplasmies) whereas (for long times) mutation rate and turnover rates define the rate of accumulation of homoplasmic mutations (in manner independent of cellular mtDNA population size)

### S8.5 Correlation between donor age and mitochondrial load as a function of the heteroplasmy thresholds

In the main manuscript, we demonstrated that the cellular mitochondrial load *μ*^10 %^ of cryptic, not synonymous mutations above a heteroplasmy of 10 % is positively correlated with the age *t* of its donor. This indicates that cells accumulate high-heteroplasmy cryptic mutations throughout ageing in accord with our theory (see SI-section S1). To test whether this result depends on the heteroplasmy threshold *h ∈* [0, 1], we compute the Spearman correlation *Corr*(*μ*^10 %^, *t*) for various heteroplasmy thresholds (see Fig. S25). We find that for all heteroplasmy thresholds *h >* 10 %, the correlation is significant at a threshold of *σ* = 0.05. Thus, we choose *h* = 10 % as default heteroplasmy threshold to keep the most data. In Fig. S25, We observe that the *p*-value of correlation decreases for low thresholds because low-heteroplasmy mutations are present across all ages. For larger thresholds we filter out biologically significant mutations and therefore the correlation decreases, leading to an increase in the *p*-value with *t*. For *h* ∈ {10 %, 30 %, 40 %, 95 %*}*, we show the scatters for (*μ*^*h* %^, *t*) in Fig. S26. The correlation is still significant for *h* = 95 %, which is in accordance with the positive correlation between the mean number of fixed cryptic mutations per cell and the age of the donor (see Fig. 1i in main text).

**Supplementary Figure S25:**
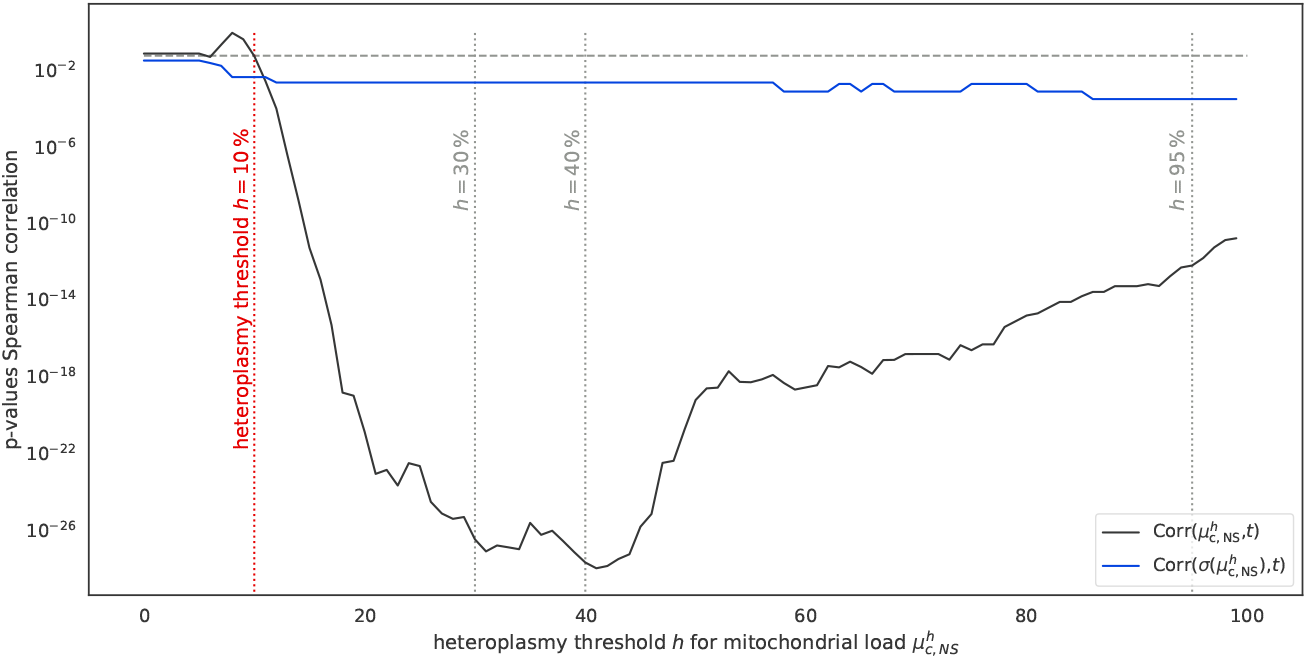
The correlation between mitochondrial load *μ*^*h* %^ and age *t* is statistically significant for a wide range of heteroplasmy thresholds *h* for the full-length RNA-seq human pancreas data (Enge (*S5*)). This also holds for the correlation between the standard deviation *σ*(*μ*^*h* %^) of the mitochondrial load and age *t*. The dashed horizontal line indicates a significance threshold of 0.05. We observe that the p-value of the correlation decreases for low thresholds because low-heteroplamsy mutations are present across all ages. For larger thresholds we filter out biologically significant mutations and therefore the correlation decreases. The correlations are still significant for *h* = 1 because the homoplasmic mutations increase with age, as show in Fig. 1i in the main text.

**Supplementary Figure S26:**
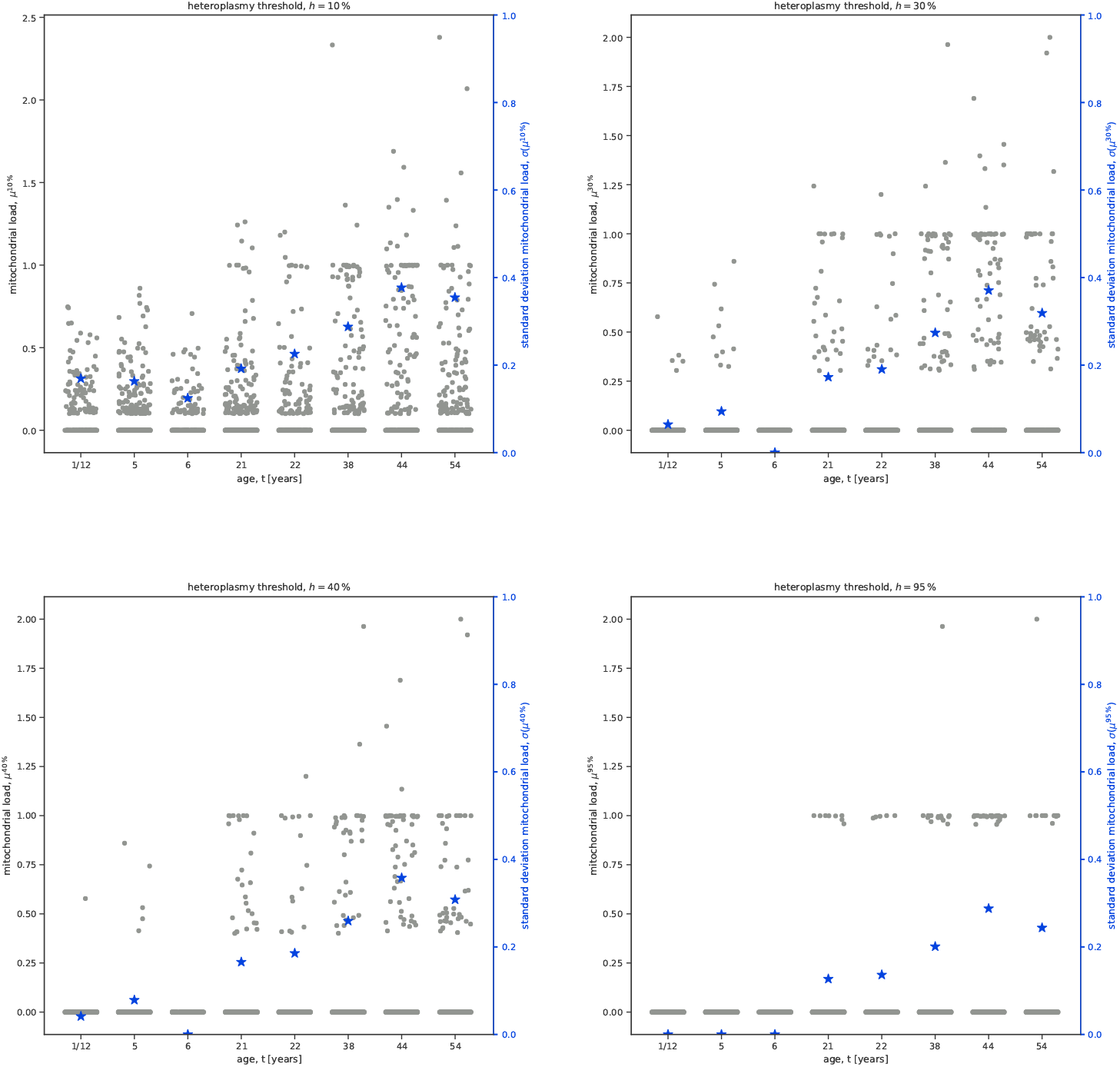
Scatterplots of mitochondrial load *μ*^*h* %^ versus donor age *t* for all cells in the full-length RNA-seq human pancreas data (Enge (*S5*)). We give results for four heteroplasmy thresholds *h* ∈ {10 %, 30 %, 40 %, 95 %} of which we use *h* = 10 % in the main manuscript.

### S8.6 Volcano plots for full-length human pancreas data at different heteroplasmy thresholds *h*

In the main manuscript, we show and discuss the differentially expressed genes (DEGs) in the full-length human pancreas data (*S5*) at a heteroplasmy threshold *h* = 10 %. Here, we show DEGs for four heteroplasmy thresholds *h* ∈ {10 %, 30 %, 40 %, 95 %} (see Fig. S27).

For all these heteroplasmy thresholds, we observe a large number of DEGs. Notably, see that different genes are covary with the presence of mtDNA mutations at different heteroplasmy thresholds *h*. This is in accord with the well-established notion of mitochondrial threshold effects, which mean that genetic effects manifest often only if they are present in sufficiently many mtDNA molecules (*S35, S36*). Nevertheless, we also find a substantial number of genes deferentially expressed across thresholds.

Surprisingly, we find that despite the small number of (almost) homoplasmic mtDNA mutations, we identify DEGs at a heteroplamsy threshold of 95 %. This indicates, that these mutations alter the gene expression substantially.

**Supplementary Figure S27:**
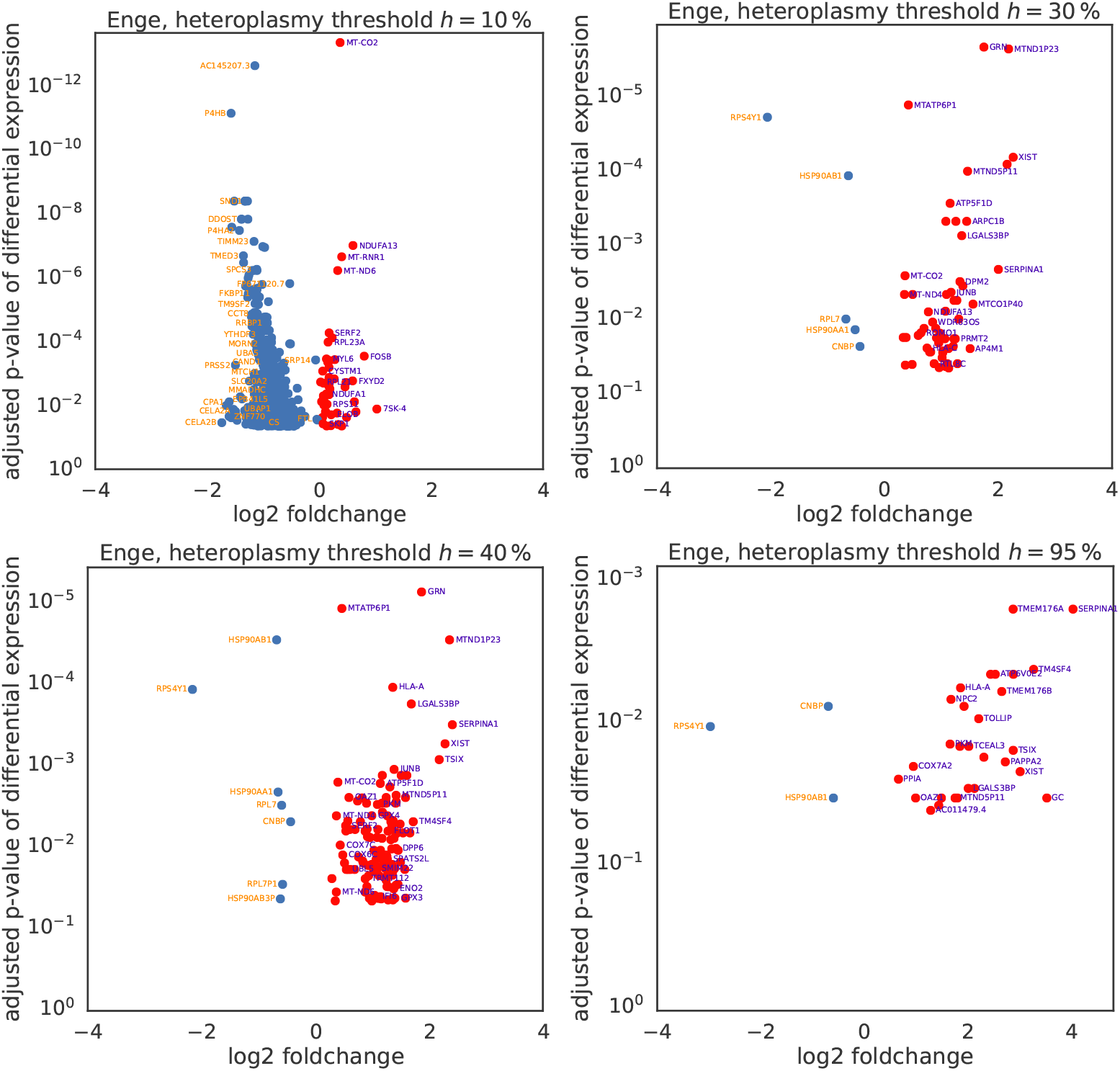
Differentially expressed genes in the full-length human pancreas data for four heteroplasmy thresholds *h ∈ {*10 %, 30 %, 40 %, 95 %*}*.

### S8.7 Volcano plots for all data sets

In the main manuscript, we show DEGs for the Enge data set (human pancreas). Here we show analogous plots for all data sets.

**Supplementary Figure S28:**
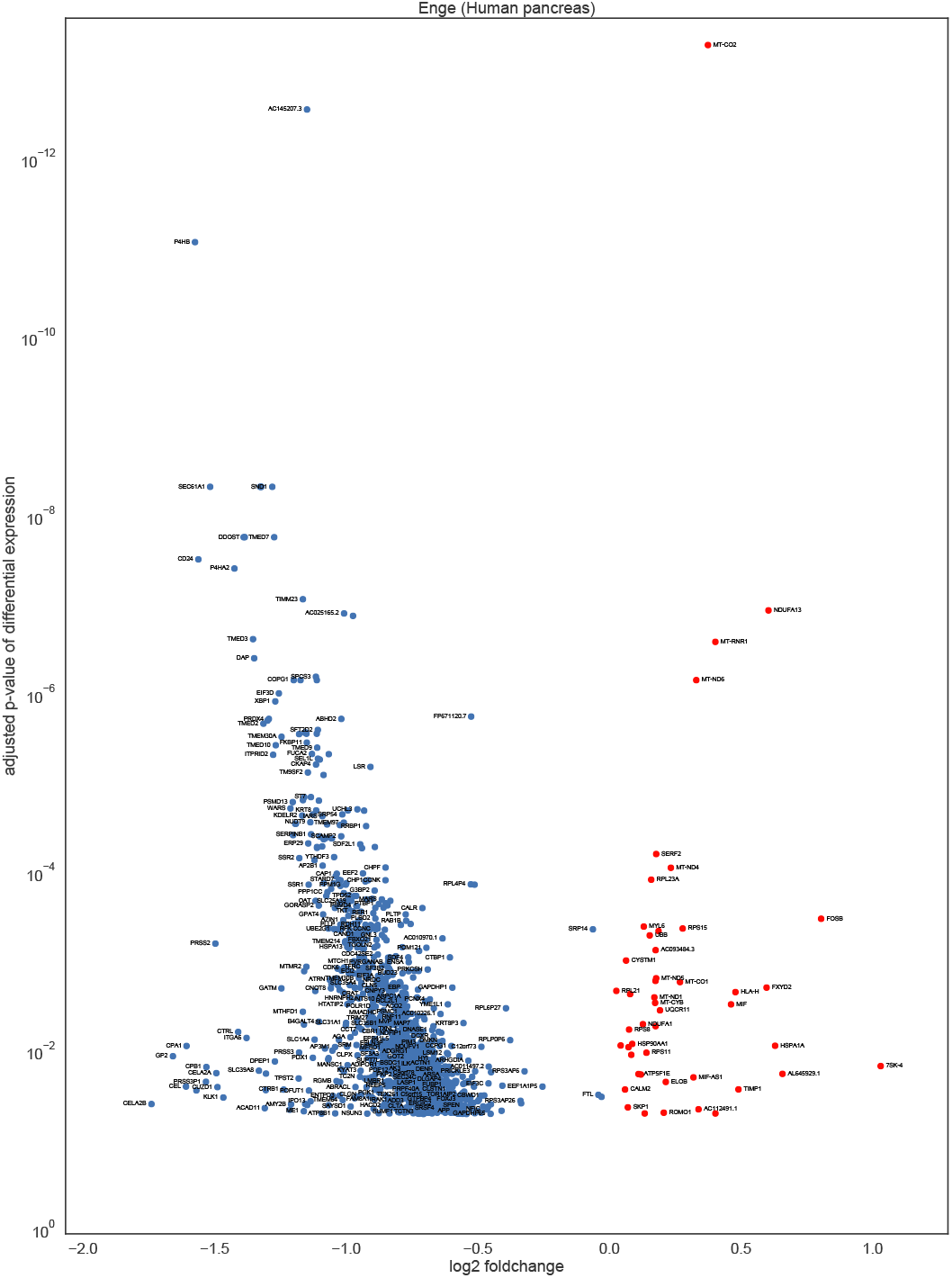
We show the 1342 differently expressed genes (DEGs) in the full-length human pancreas data (Enge (*S5*)). For each DEG, we show the log2-foldchange and the adjusted *p*-value of differential expression, as computed by a Wilcoxon rank sum test. We label selected genes.

**Supplementary Figure S29:**
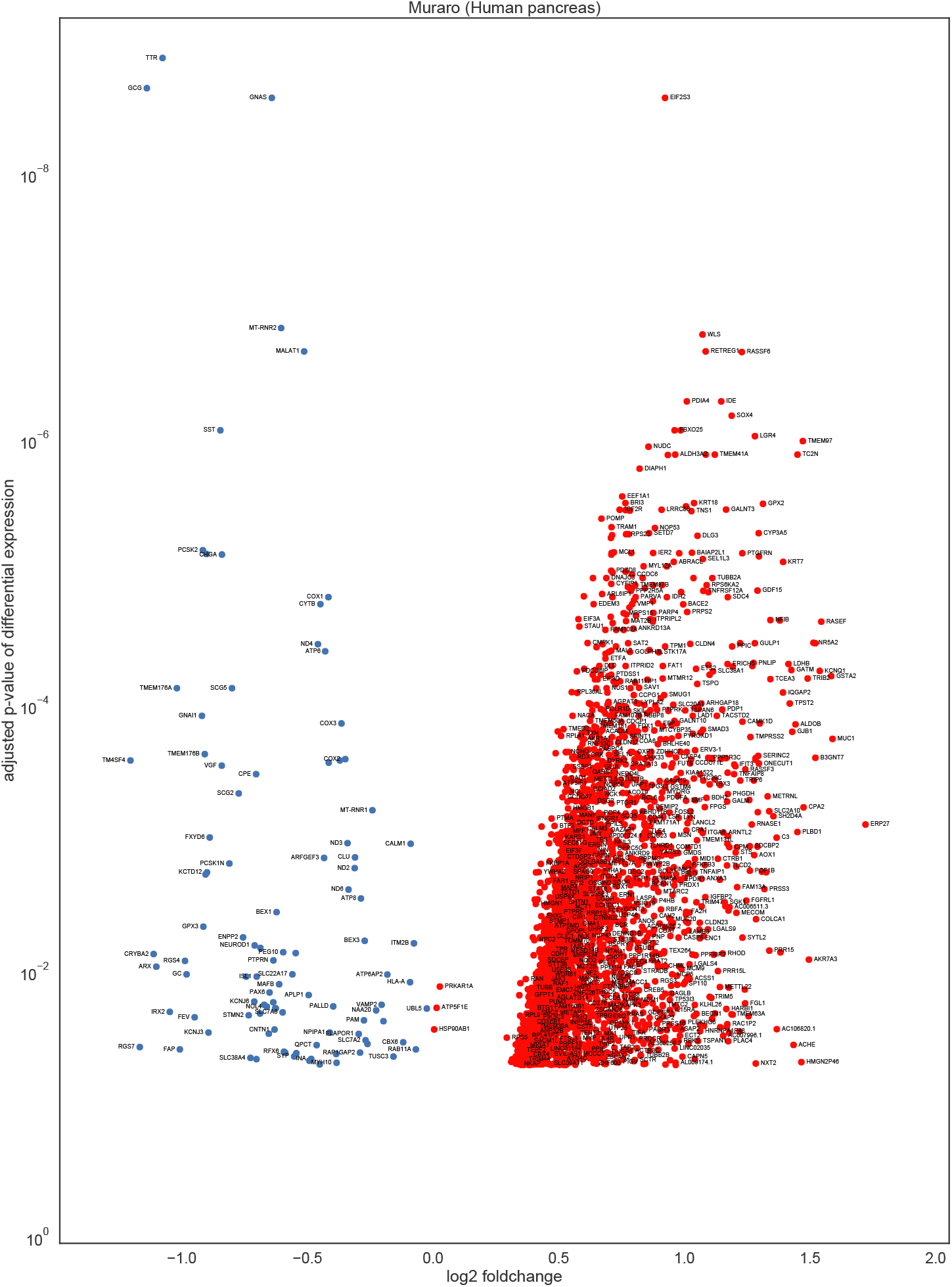
We show the 3319 differently expressed genes (DEGs) in the human pancreas data (Muraro (*S15*)). For each DEG, we show the log2-foldchange and the adjusted *p*-value of differential expression, as computed by a Wilcoxon rank sum test. We label selected genes.

**Supplementary Figure S30:**
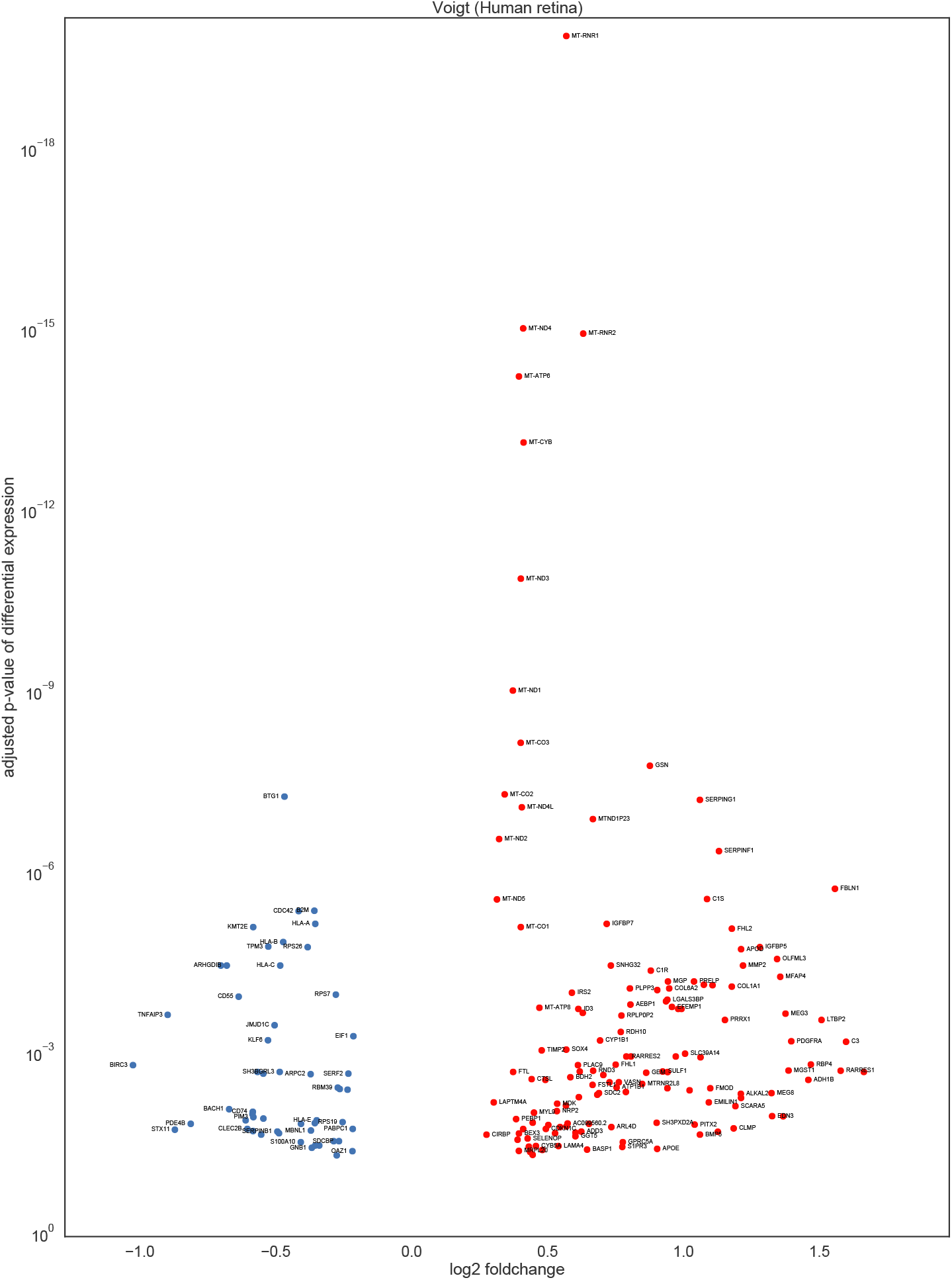
We show the 189 differently expressed genes (DEGs) in the human retina data (Voigt (*S10*)).

**Supplementary Figure S31:**
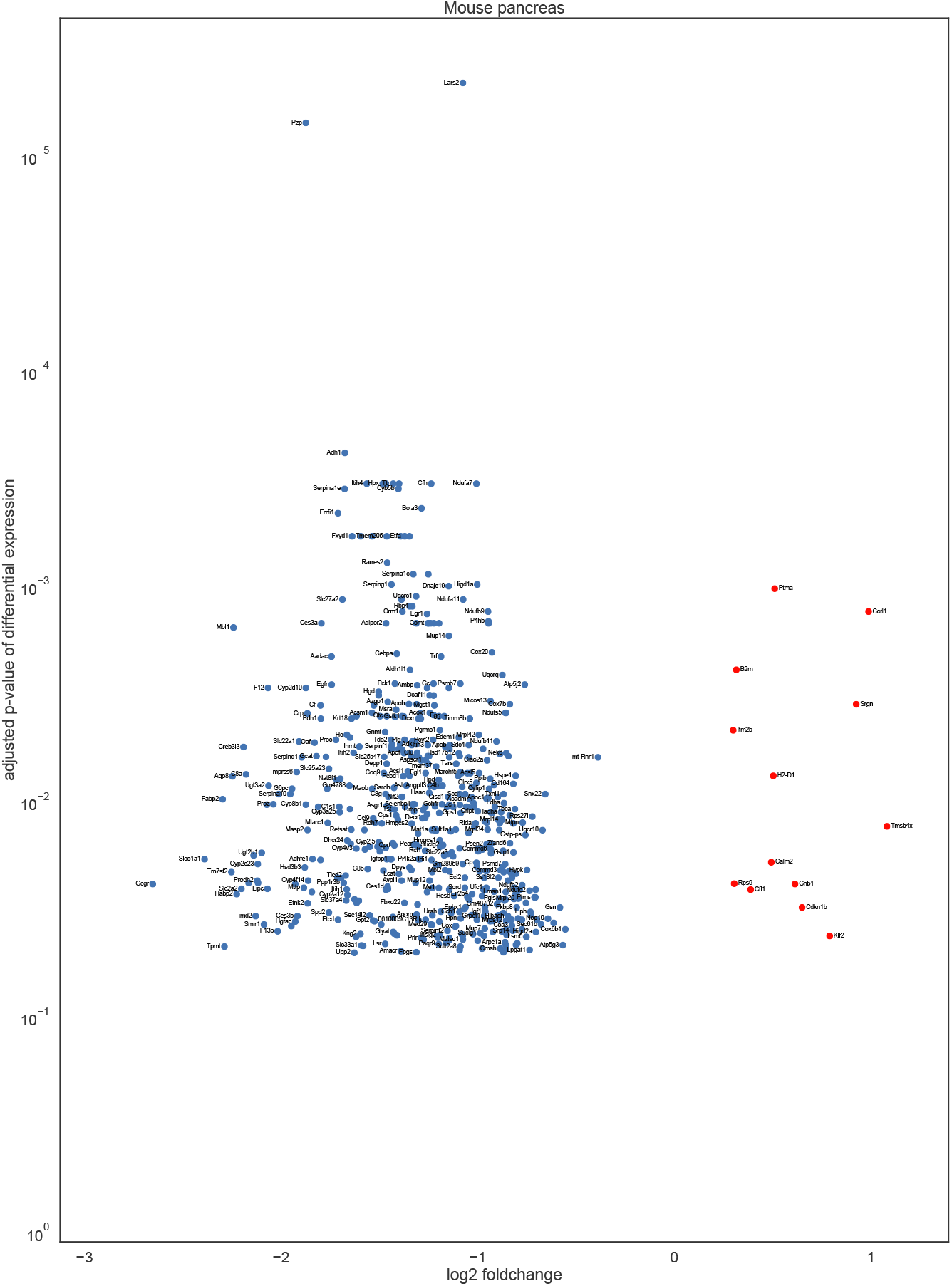
We show the 546 differently expressed genes (DEGs) in the mouse liver data (Tabula Muris (*S16*)).

**Supplementary Figure S32:**
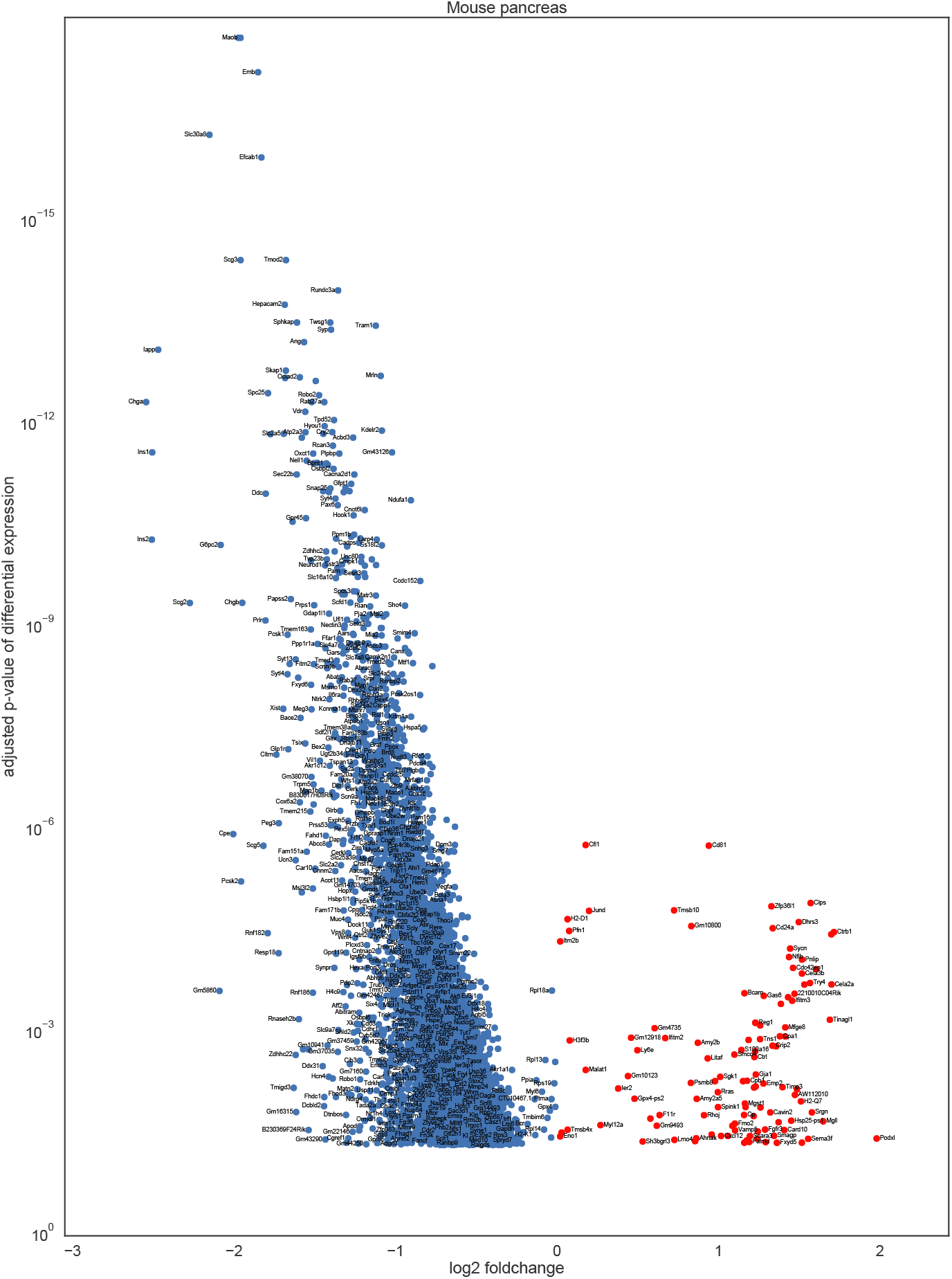
We show the 5430 differently expressed genes (DEGs) in the mouse pancreas data (Tabula Muris (*S16*)). For each DEG, we show the log2-foldchange and the adjusted *p*-value of differential expression, as computed

**Supplementary Figure S33:**
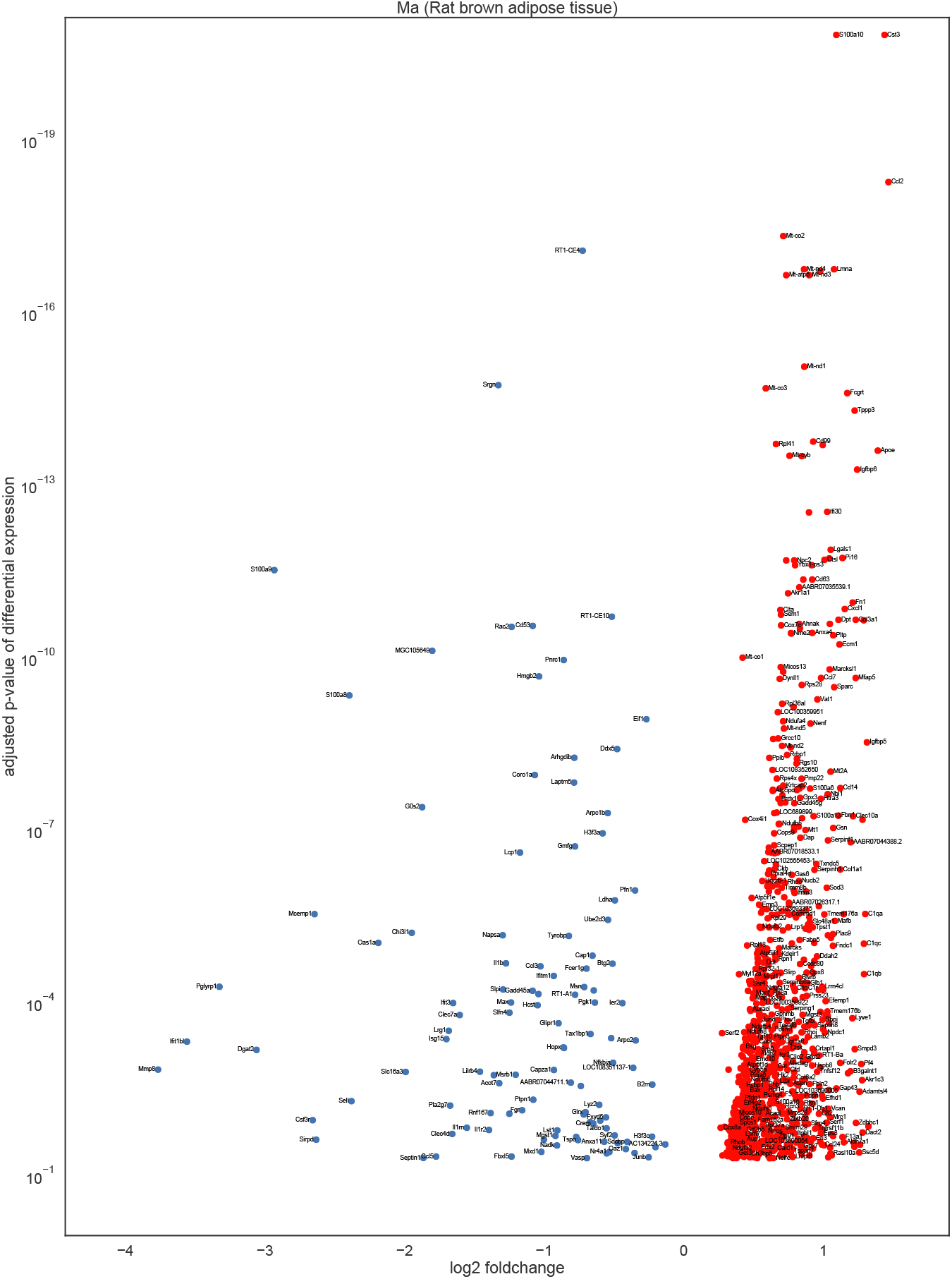
We show the 1691 differently expressed genes (DEGs) in the rat brown adipose tissue data (Ma *et al*. (*S34*)). For each DEG, we show the log2-foldchange and the adjusted *p*-value of differential expression, as computed by a Wilcoxon rank sum test. We label selected genes.

**Supplementary Figure S34:**
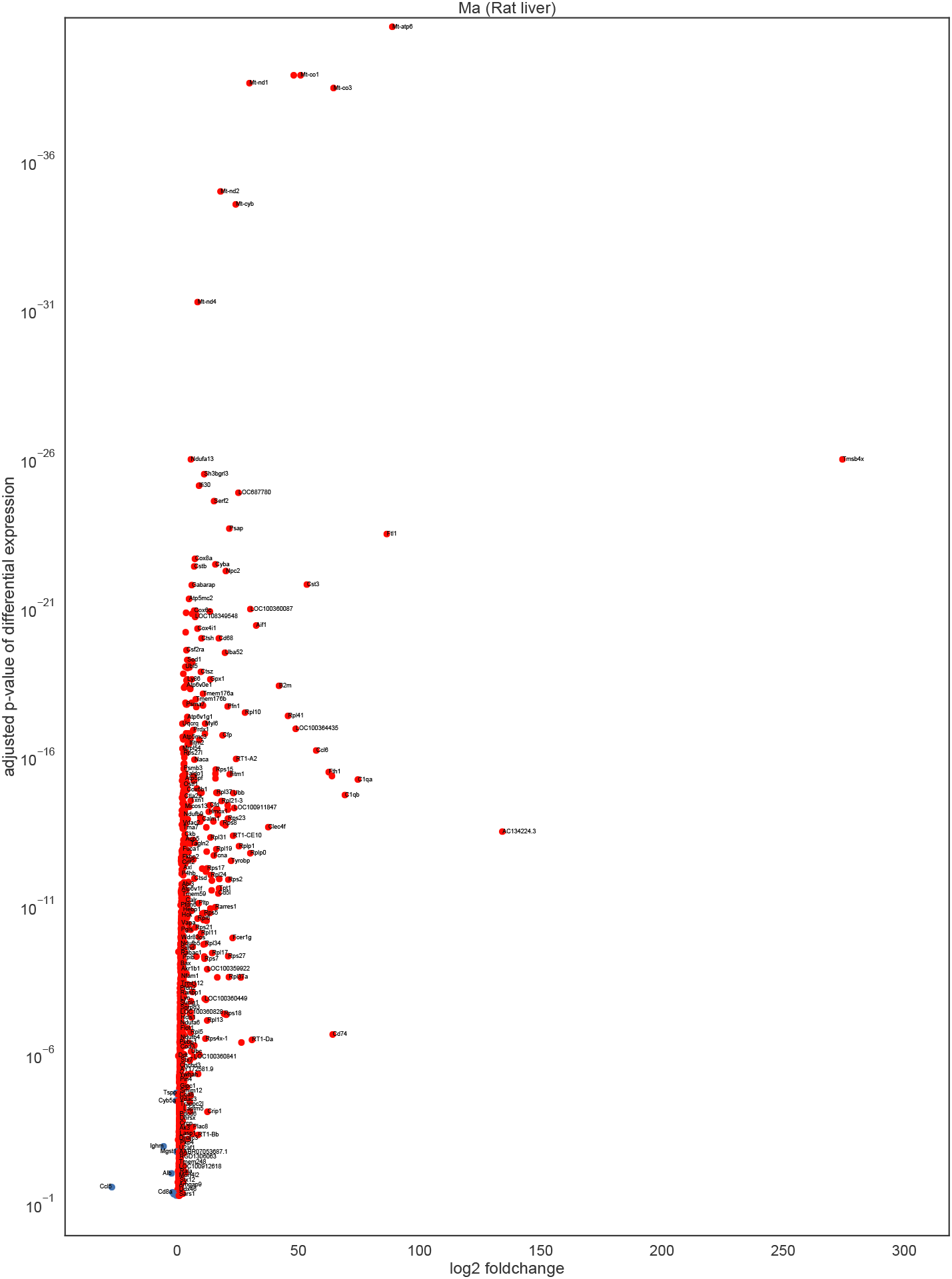
We show the 1003 differently expressed genes (DEGs) in the rat liver data (Ma *et al*. (*S34*)). For each DEG, we show the log2-foldchange and the adjusted *p*-value of differential expression, as computed by a Wilcoxon rank sum test. We label selected genes.

### S8.8 Processing pipeline

**Supplementary Figure S35:**
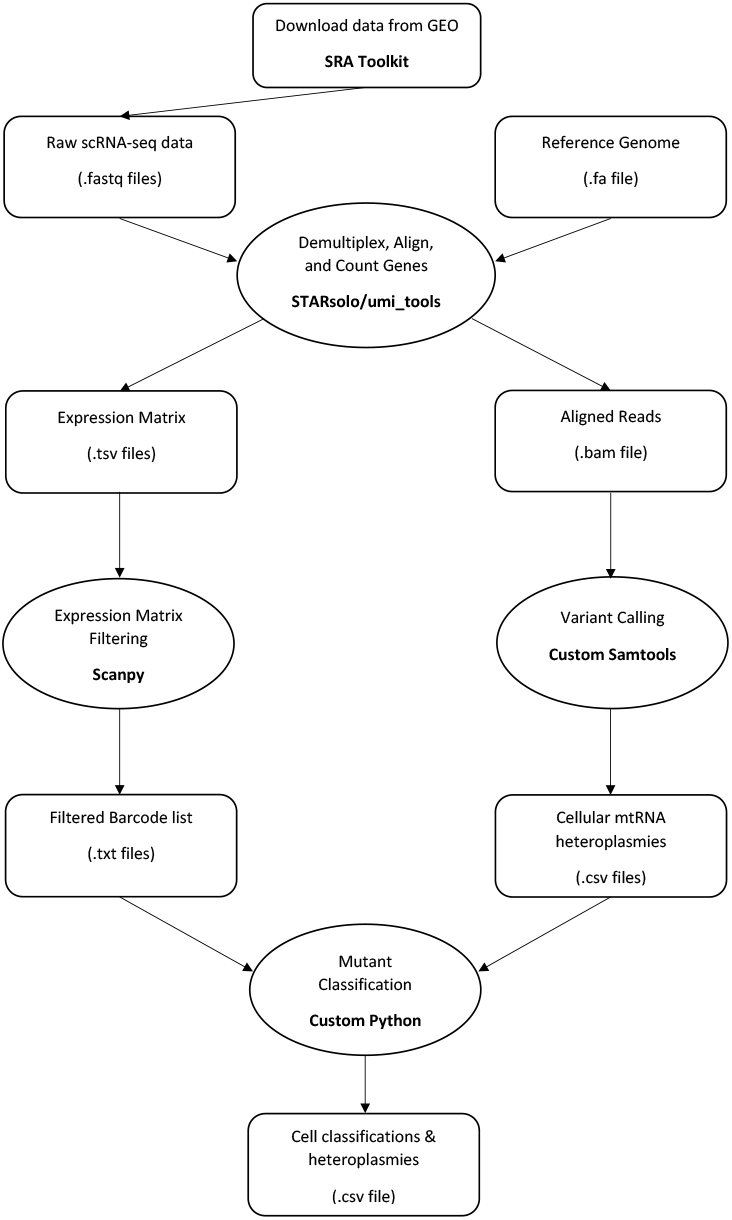
Directed acyclic graph representing the data processing pipeline. After downloading the raw scRNA-seq data as .fastq files, we align them to a reference genome, which yields aligned reads in a `.bam` file and an expression matrix. Using custom Python tools, this gives us for each cell the heteroplasmy of mitochondrial mutations and also the gene-expression data. Then using custom Jupyter notebooks we perform quality control on the mitochondrial content of cells before assigning mutant classes to every heteroplasmy based on the number of cells from a donor it is found in.

### S8.9 Differentially expressed genes for location-specific cryptic mtDNA mutations in human pancreas

**Supplementary Figure S36:**
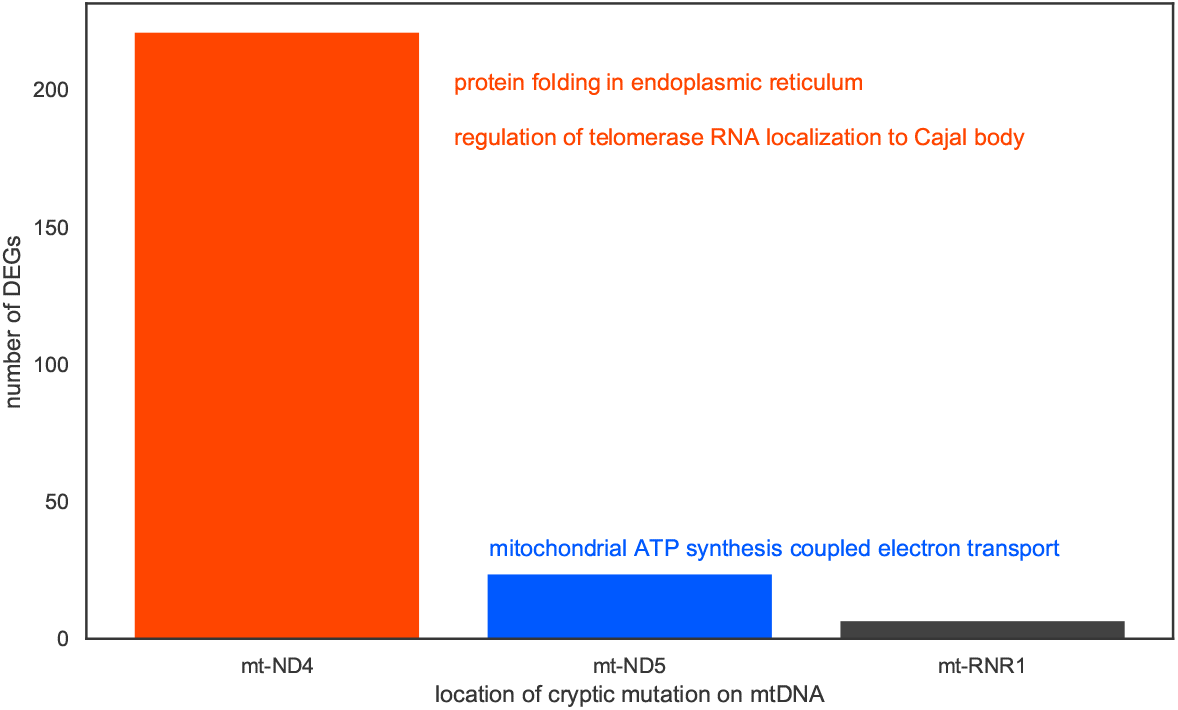
DEGs for cryptic mtDNA mutations are dominated by mutations at the mt-ND4, mt-ND5, and mt-RNR1 genes, each of which indicate distinct functional perturbations. For the full-length human pancreas data (Enge), we compute DEGs at a *t* = 10 % threshold for mutations that are located at each of the mitochondrial genes. For most mitochondrial genes, we do not find DEGs (not shown). For the mt-ND4, mt-ND5, and mt-RNR1 genes, however, we identify 220, 23, and 6 DEGs, respectively. A GO term-enrichment indicates that cryptic mt-ND4 mutations perturb ‘protein folding in the ER’ and ‘regulation of telomerase RNA localization to Cajal body’, which is consonant with the stress response that we discuss in the main manuscript. DEGs from cryptic mt-ND5 mutations in contrast are associated with various mitochondrial functions, particularly ‘mitochondrial ATP synthesis coupled electron transport’, indicating dysregulation of OXPHOS energy production. The six DEGs associated with cryptic mt-RNR1 mutations (TRAM1, CYSTM1, UBB, SSR2, AC093484.3, SF3B2) are not enriched for a GO term.

